# Avian influenza viruses in wild birds in Canada following incursions of highly pathogenic H5N1 virus from Eurasia in 2021/2022

**DOI:** 10.1101/2023.11.23.565566

**Authors:** Jolene A. Giacinti, Anthony V. Signore, Megan E. B. Jones, Laura Bourque, Stéphane Lair, Claire Jardine, Brian Stevens, Trent Bollinger, Dayna Goldsmith, British Columbia Wildlife AIV Surveillance Program (BC WASPs), Margo Pybus, Iga Stasiak, Richard Davis, Neil Pople, Larissa Nituch, Rodney W. Brook, Davor Ojkic, Ariane Massé, Gabrielle Dimitri-Masson, Glen J. Parsons, Meghan Baker, Carmencita Yason, Jane Harms, Naima Jutha, Jon Neely, Yohannes Berhane, Oliver Lung, Shannon K. French, Lawrna Myers, Jennifer F. Provencher, Stephanie Avery-Gomm, Gregory J. Robertson, Tatsiana Barychka, Kirsty E. B. Gurney, Jordan Wight, Ishraq Rahman, Kathryn Hargan, Andrew S. Lang, Michael G. C. Brown, Cynthia Pekarik, Trevor Thompson, Angela McLaughlin, Megan Willie, Laurie Wilson, Scott A. Flemming, Megan V. Ross, Jim Leafloor, Frank Baldwin, Chris Sharp, Hannah Lewis, Matthieu Beaumont, Al Hanson, Robert A. Ronconi, Eric Reed, Margaret Campbell, Michelle Saunders, Catherine Soos

## Abstract

Following detection of novel highly pathogenic avian influenza virus (HPAIV) H5N1 clade 2.3.4.4b in Newfoundland, Canada in late 2021, avian influenza surveillance in wild birds was scaled-up across Canada. Herein, we present results of Canada’s Interagency Surveillance Program for Avian Influenza in wild birds during the first year (November 2021 – November 2022) following the incursions of HPAIV from Eurasia. Key objectives of the surveillance program were to (i) detect the presence, distribution and spread of HPAIV and other avian influenza viruses (AIVs), (ii) detect wild bird morbidity and mortality associated with HPAIV, (iii) identify the range of wild bird species infected by HPAIV, and (iv) characterize detected AIV. A total of 6,246 sick and dead wild birds were tested, of which 27.4% were HPAIV positive across 12 taxonomic orders and 80 species. Geographically, HPAIV detections occurred in all Canadian provinces and territories, with the highest numbers in the Atlantic and Central flyways. Temporally, peak detections differed across flyways, though the national peak occurred in April 2022. In an additional 11,295 asymptomatic harvested or live captured wild birds, 5.2% were HPAIV positive across 3 taxonomic orders and 19 species. Whole genome sequencing identified HPAIV of Eurasian origin as most prevalent in the Atlantic flyway, along with multiple reassortants of mixed Eurasian and North American origins distributed across Canada, with moderate structuring at the flyway scale. Wild birds were victims and reservoirs of HPAIV H5N1 2.3.4.4b, underscoring the importance of surveillance encompassing samples from sick and dead, as well as live and harvested birds to provide insights into the dynamics and potential impacts of the HPAIV H5N1 outbreak. This dramatic shift in presence and distribution of HPAIV in wild birds in Canada highlights a need for sustained investment in wild bird surveillance and collaboration across One Health partners.

## Introduction

Since the detection of highly pathogenic avian influenza (HPAI) H5N1 virus clade 2.3.4.4b in Canada in December 2021, there have been significant impacts for wildlife and domestic poultry health. Globally, this clade is associated with unprecedented impacts on wild birds and mammals compared to previous H5Nx highly pathogenic avian influenza virus (HPAIV) in that it has a wider host range (1), a larger geographic range, facilitated by multiple instances of inter- and intracontinental spread (2–5), higher mortality levels of wild birds, mesocarnivores, and marine mammals (6–8), and longer persistence in wild bird populations in Europe (9). In Canada, this virus is also associated with unprecedented impacts on commercial, small flock, and other captive poultry facilities that far surpass the mortality and economic losses associated with the only other HPAIV incursion into North America in 2014/2015, which resulted in outbreaks on 16 premises in two provinces (10, 11). As of September 2023, 7.7 million domestic birds have been destroyed on 319 premises across nine Canadian provinces (12).

Canada’s Interagency Surveillance Program for Avian Influenza Viruses in Wild Birds (previously called Canada’s Interagency Wild Bird Influenza Survey) has been operating since 2005 (13). The program consists primarily of two core components: (i) morbidity and mortality surveillance in wild birds submitted opportunistically, often by members of the public, and (ii) surveillance in live and hunter-harvested wild birds sampled, often in conjunction with existing banding, research, or monitoring programs. Since late 2021, Environment and Climate Change Canada (ECCC) has worked with the Canadian Wildlife Health Cooperative (CWHC), provincial/territorial government agencies, other federal departments (Canadian Food Inspection Agency (CFIA), Public Health Agency of Canada, Parks Canada, and Indigenous Services Canada), and Indigenous and academic partners to increase surveillance for HPAIV in wild birds across the country (Supplemental Document 1).

Herein we describe the epidemiology of the HPAIV outbreak in wild birds in Canada from November 2021 to November 2022 by addressing several of the primary surveillance objectives related to reporting spatiotemporal dynamics, host taxonomic representation, and characterizing viral genetic diversity.

## Materials and Methods

### Morbidity and Mortality Surveillance

Morbidity and mortality surveillance of wild birds in Canada was largely opportunistic, requiring that sick or dead birds be found, reported, and samples submitted to the CWHC, provincial or territorial agencies or laboratories, often by the public. Avian carcasses were submitted fresh or frozen for processing. In some cases, sick birds were admitted to rehabilitation facilities prior to the submission of the carcass for AIV testing. Because of the increased volume of carcass submissions in 2021 and 2022, carcass testing was prioritized across Canada according to field and diagnostic lab capacity as well as funding (Supplemental Document 1, Appendix D). Oropharyngeal and cloacal swabs were collected from carcasses selected for avian influenza testing and pooled into a single vial containing appropriate transport medium. Vials were stored at temperatures of at least −20 °C until testing, and −70 to −80 °C when available (Supplemental Document 1, Appendix G). When resources and capacity were available, gross and histologic examination of carcasses with HPAIV positive swab results was undertaken at the CWHC or provincial or territorial laboratory to confirm HPAIV as the cause of death and to help rule out false positives particularly in the case of new species or new locations. In some cases, tissue samples (brain, lung, and intestine) collected during post-mortem examination were submitted in lieu of swabs for AIV testing.

### Live and Hunter-Harvested Bird Surveillance

Live wild birds were sampled by ECCC, the United States Fish and Wildlife Service, and provincial or territorial, and Indigenous or academic partners. Sampling opportunities were reviewed periodically to prioritize sample collection and ensure they were in line with surveillance objectives, sample size recommendations (Supplemental Document 1, Appendix E), capacity, and resources. All live bird sampling was performed in accordance with approved animal use protocols, appropriate federal or provincial wildlife permits where applicable, and appropriate safe work procedures. Samples from harvested birds were provided by permitted and Indigenous harvesters. For the purposes of this study, harvested birds are considered apparently healthy prior to harvest and are therefore categorized with live birds in our analyses. Live and harvested birds were sampled for AIV as described above.

### Laboratory Analyses

Real-time reverse transcriptase polymerase chain reaction (RT-PCR) testing of swab samples and tissue samples was performed at the diagnostic laboratories of the Canadian Animal Health Surveillance Network (CAHSN), which is a network of federal, provincial, and university animal health laboratories across Canada with the central reference laboratory operating from the National Centre for Foreign Animal Disease, Canadian Food Inspection Agency (NCFAD-CFIA) in Winnipeg, Manitoba. The CAHSN standard protocol for the detection of type A influenza viruses and avian H5 and H7 hemagglutinin subtypes by RT-PCR Assay (Version 3, January 2020) was utilized. In this protocol, a RT-PCR assay based on the use of fluorescent 5’ nuclease oligoprobes (hydrolysis probes) was used for rapid detection of group A specific Matrix (M1), H5, and H7 hemagglutinin subtype avian influenza virus sequences. The matrix assay also employs the use of an exogenous armored RNA-Enterovirus internal control for verification of the RNA extraction step and detection of PCR inhibitors. The matrix RT-PCR is designed to detect M1 gene sequences of all group A influenza viruses (birds and mammals). The H5NA/EA RT-PCR Assay in the CAHSN protocol is capable of detecting most North American and Eurasian lineage H5 avian influenza viruses, including Eurasian H5N1 viruses. The H7 2013 RT-PCR has been re-designed to detect H7 influenza viruses from the Americas as well as Eurasia.

Automated nucleic acid extraction from samples was performed using Magnetic Particle Processors (MagMax, KingFisher, Roche and others) and appropriate kits, while the manual nucleic acid extraction was done using Qiagen vacuum manifold and Qiagen Viral RNA MiniKit. The following RT-PCR Systems were used: Applied Biosystems 7500/7500 Fast, Roche Light Cycler 480, BioRad CFX 96 and Strategene MX3005. The interpretation of test results as outlined in the CAHSN standard protocol was followed.

All samples that were positive for Group A specific Matrix RT-PCR and positive, suspect, or non-negative for H5 or H7 were sent to NCFAD-CFIA for confirmatory testing and further genomic characterization.

We assigned AIV sample status as “confirmed or suspect HPAIV-positive” or “confirmed or suspect low pathogenicity avian influenza virus (LPAIV)-positive”. The former includes cases confirmed H5 HPAIV-positive (highly pathogenic virus of the subtype H5Nx confirmed by NCFAD-CFIA) and samples that were non-negative on H5 PCR at the regional laboratory or NCFAD-CFIA, but virus isolation and sequencing were not possible due to sample quality or in a few cases, testing is not yet completed. These samples were categorized as suspect H5 HPAIV-positive because there were few LPAIV H5 detections in Canada over the study period (Y. Berhane, personal communication). Confirmed or suspected LPAIV-positive include cases with low pathogenicity avian influenza virus confirmed by NCFAD-CFIA and those that tested non-negative on matrix PCR and negative on H5 PCR at the regional laboratory or NCFAD-CFIA. In the latter cases, additional virus isolation and sequencing was not possible or is not yet completed.

### Wild Bird Surveillance Data

Metadata and preliminary matrix, H5, and H7 RT-PCR results for sick and dead birds across Canada were received from surveillance partners. Metadata and diagnostic results associated with live and harvested wild birds were managed internally within ECCC. Confirmatory diagnostic results were compiled by NCFAD-CFIA. Data were regularly merged, structured, and samples were assigned additional identifiers (e.g., taxonomic family, flyway, watershed) resulting in a compiled national AIV surveillance dataset (12). Exact (Clopper-Pearson) confidence intervals were calculated in R version 4.2.2 (2022-10-31). The best available data are presented for surveillance conducted between November 2021 - December 2022, extracted from the full dataset on May 15, 2023.

### Influenza Virus Genome Sequencing and Assembly

Full genome segments of the AIVs were amplified either directly from clinical specimens or isolates as described previously (14). High-throughput sequencing was performed either on an Oxford Nanopore Technologies (ONT) GridION sequencer and R9.4.1 Flow Cell following library (n=921) construction using the ONT rapid barcoding kit (SQK-RBK004 or SQK-RBK110.96) or an Illumina MiSeq (n=257) using the Nextera XT Library Preparation kit (Illumina) following the manufacturers’ protocol. The Hamilton Microlab Star Robot was used for Illumina library preparation prior to sequencing with Illumina MiSeq Reagent Kits (300 cycle or 600 cycle) paired with either Illumina MiSeq V2 or V3 Flow Cells. The raw Nanopore signal data was basecalled and demultiplexed with the latest version of Guppy at the time of sequencing (v5.0.17 – v6.5.7) using the high-accuracy or super-high accuracy models. Basecalled Nanopore reads were analysed and assembled with the CFIA-NCFAD/nf-flu v3.3.6 Nextflow workflow (15, 16) which ran IRMA (v1.0.2) for initial genome assembly (17); nucleotide BLAST v2.14 (18, 19) search of IRMA assembled genome segment sequences against all Orthomyxoviridae sequences from the NCBI FTP site (https://ftp.ncbi.nlm.nih.gov/genomes/Viruses/AllNucleotide/; 1,070,105 sequences downloaded 2023-06-14); selection of appropriate reference sequence for each genome segment and H/N subtype prediction based on nucleotide BLAST results; Minimap2 v2.24 (20) read mapping to each genome segment reference sequence; Samtools v1.15 (21, 22) and Mosdepth v0.3.3 (23) for read mapping and sequencing coverage statistics; Clair3 v1.0.2 (24) variant calling; Bcftools v1.15.1 (22) variant filtering and depth-masked consensus sequence generation for each genome segment; and MultiQC v1.12 (25) for bioinformatics analysis summary report creation. Bcftools generated consensus sequences from Nanopore analysis with nf-flu were used for further analyses. For Illumina sequencing reads, IRMA (v1.0.2) Influenza genome assembly as part of the CFIA-NCFAD/nf-flu (v3.3.6) workflow was used to generate the consensus sequences used for further analyses. All viral genome sequences generated in this study will be deposited on GISAID before final peer-reviewed publication.

### Phylogenetic Analyses

Individual viral segments (PB2, PB1, PA, HA, NP, NA, M and NS) were trimmed of regions flanking the open reading frames and concatenated. The geographic origin (either Eurasian or North American) of each genome segment prior to concatenation was assessed by BLAST search similarity against reference sequences defined with segment-specific phylogenies from Alkie et al. (26). Concatenated HPAIV H5N1 sequences were aligned using MAFFT v7.49 (totaling 13,112 nucleotides in length) (27) and used to build a maximum likelihood phylogenetic tree using IQ-TREE v2.20 (28). A separate partition was designated for each viral segment, allowing each to have its own model of nucleotide substitution and model specific parameters as determined by ModelFinder (29). Node support for the resulting tree was assessed by 5000 ultrafast bootstrap replicates (30). The bootstrap consensus tree was re-rooted on the first H5N1 virus detected in Canada (A/Great_Black-Backed_Gull/NL/OTH-0114-1/2021) and sampling dates for each tip were used to time-scale the tree under a relaxed molecular clock rate in TreeTime (31). Reconstruction of the ancestral hosts’ taxomonic order in the time-calibrated phylogenetic tree was conducted using the TreeTime mugration model. The resulting phylogenetic tree, reassortment pattern, and host taxonomy was visualized using R package ggtree v3.7.2 (32). Inference of segment-specific phylogenies were used to identify unique genome constellations (i.e., reassortant genotypes) for a more detailed phylogeographic reconstruction in a separate manuscript (Signore et al. in prep.).

### Additional Data

Information on infected domestic bird premises in Canada was obtained from the Canadian Food Inspection Agency (12). Unusual wild bird mortality event information was obtained from the National Environmental Emergencies Centre situation report (33), which includes information received from surveillance partners across Canada throughout the course of the HPAIV epidemic in Canada. These data were supplemented by additional information from provincial/territorial wildlife agencies and information obtained through a regional collaborative effort to document HPAIV-related mortality estimates in Atlantic Canada (Avery-Gomm et al., in prep).

We simplified migratory flyway boundaries according to provincial/territorial divisions in Canada (34), acknowledging that migration does not precisely align with administrative boundaries and adjacent flyways overlap in some areas. ArcMap Pro v3.0.0 was used for mapping.

## Results

### HPAIV Outbreak Timeline

The presumed index case among wild birds of the 2021/2022 outbreak in North America was a first-winter Great Black-Backed Gull (*Larus marinus*; order Charadriiformes) from Newfoundland. This bird was found exhibiting neurologic signs including inability to fly, head tilt, ataxia, and depression. This and two other birds with similar histories were found alive between November 4 – 26, 2021, all three died within 24 hours of admission to a wildlife rehabilitation centre, were submitted to the CWHC in late December, and confirmed HPAIV positive the same month (Fig. 1). HPAIV was detected within the Atlantic flyway in geese (*Anatidae spp.*) or raptors (Accipitriformes, Strigiformes, Falconiformes) in January (Nova Scotia, Prince Edward Island), February (New Brunswick), and March (Quebec), 2022. A separate incursion of HPAIV was detected in the Pacific flyway in a Bald Eagle (*Haliaeetus leucocephalus*) in British Columbia in February 2022 (Fig. 1) (2), but there were no further detections in the province until April 2022. These detections represent bicoastal incursions by early 2022. The first detection in the mid-continental Mississippi flyway was a Red-Tailed Hawk (*Buteo jamaicensis*) in Ontario, in March 2022. Detections in southern Manitoba (Mississippi flyways) and Saskatchewan (Central flyway) were in Snow Geese (*Chen caerulescens*) and began in late March 2022 (Fig. 1). Presumed index cases in flyways, provinces, and new areas within each province typically were sick or dead or apparently healthy members of the order Anseriformes, sick or dead raptors followed by corvids (Passeriformes; Video 1). Within each flyway, the subsequent species detected through morbidity and mortality surveillance were often Charadriiformes, specifically gulls (*Larus* spp.; Video 1). The first detections in northern Canada occurred in early May in Yukon Territory, in a Canada Goose (*Branta canadensis*) and a Trumpeter Swan (*Cygnus buccinator*), and in the latter half of June in the Northwest Territories and Nunavut in Herring Gulls (*Larus argentatus*).

**Fig. 1.**
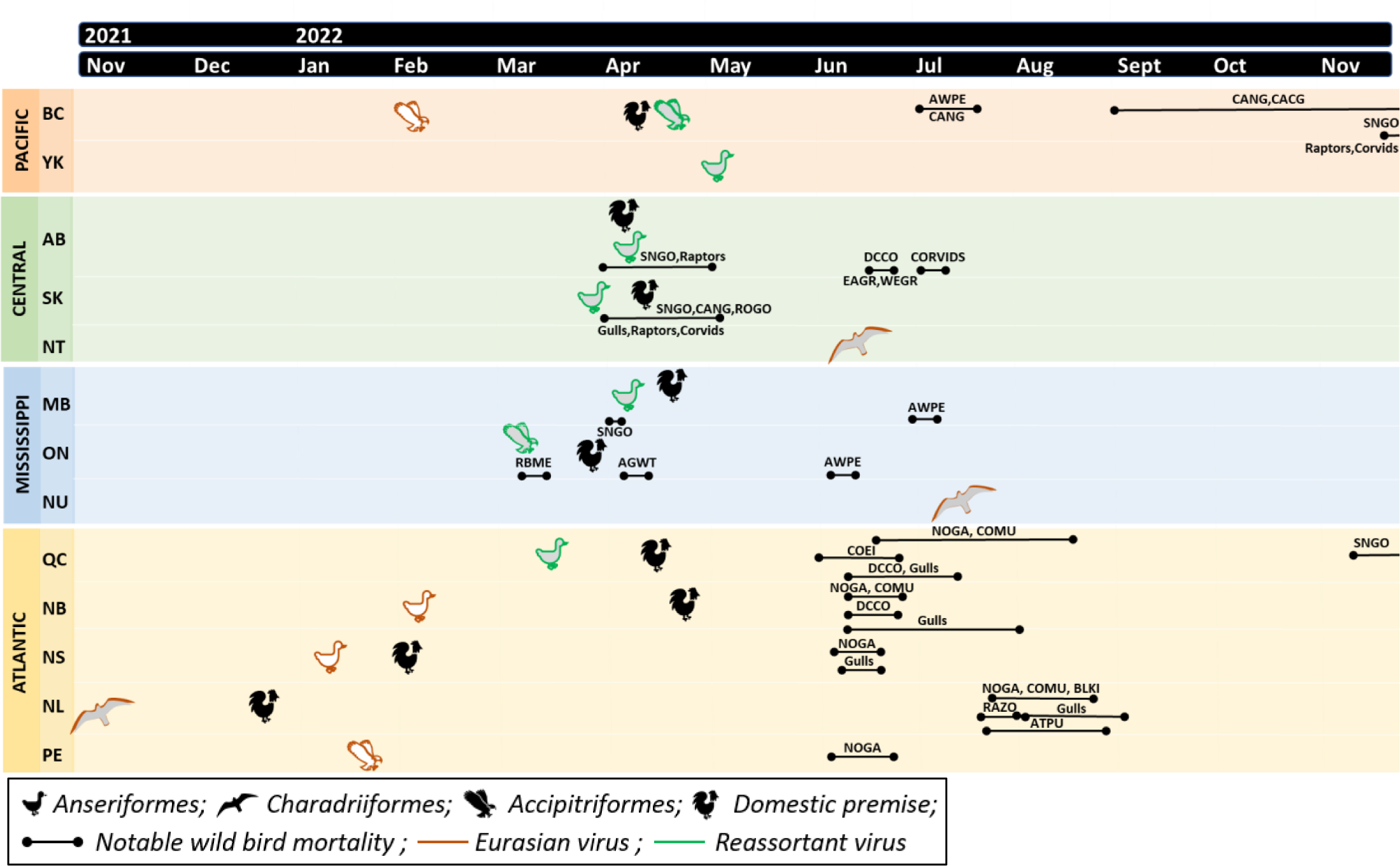
Timeline of events between November 2021 and 2022 following the first confirmed case of the clade 2.3.4.4b highly pathogenic avian influenza virus (HPAIV) in Canada. The timeline is stratified by flyway (Pacific, Central, Mississippi, and Atlantic) and province/territory (BC: British Columbia, YK: Yukon, AB: Alberta, SK: Saskatchewan, NT: Northwest Territories, MB: Manitoba, ON: Ontario, NU: Nunavut, QC: Quebec, NB: New Brunswick, NS: Nova Scotia, NL: Newfoundland and Labrador, PE: Prince Edward Island), which are denoted with colors. The first wild bird sample and domestic premise confirmed HPAIV-positive in each province is indicated with an icon. The identity of the HPAIV detected is indicated with color outline. Unusual wild bird mortalities are indicated with a solid black line spanning the relevant period of time (AGWT: American Green-Winged Teal, ARTE: Arctic Tern, ATPU: Atlantic Puffin, AWPE: American White Pelican, BLKI: Black-Legged Kittiwake, CACG: Cackling Goose, CANG: Canada Goose, COEI: Common Eider, COMU: Common Murre, COTE: Common Tern, DCCO: Double-Crested Cormorant, EAGR: Eared Grebe, NOGA: Northern Gannet, RAZO: Razorbill, RBME: Red-Breasted Merganser, ROGO: Ross’ Goose, SNGO: Snow Goose, WEGR: Western Grebe).

Multiple notable mortality events associated with HPAIV were reported across the country beginning in March 2022. Mortality was reported in Red-Breasted Mergansers (*Mergus serrator;* <100) in Ontario in mid-March 2022, followed by Green-Winged Teal (*Anas carolinensis*; <100) in early April 2022. In the central flyway beginning late March 2022, mortality events were reported in Snow Geese in southern Alberta (hundreds) and Manitoba (unreported number) and Snow Geese, Canada Geese, and Ross’ Geese in Saskatchewan (*Anser rossii*; hundreds; Fig. 1). Notable mortality in raptor species (e.g., eagles, owls, hawks) was reported in Alberta and Saskatchewan (<100) also beginning in late March 2022 in addition to gulls and corvids in Saskatchewan (<100) (Fig. 1).

Notable mortality events were also reported beginning in May 2022, but persisting throughout the spring and into early fall, predominantly in breeding colonial nesting seabirds and most prominently in eastern Canada. This included outbreaks at Northern Gannet (*Morus bassanus*) breeding colonies in Quebec and Newfoundland and at American Common Eider (*Somateria mollissima dresseri*) colonies in the Gulf of St. Lawrence, Quebec (Fig. 1; Avery-Gomm et al., in prep.) (35). Across eastern Canada, reported mortalities exceeding 40,000 wild birds, including >25,000 Northern Gannet, >8,000 Common Murre (*Uria aalge*), >1,700 Common Eider, along with numerous reports of dead gulls (>2,300), cormorants (*Phalacrocorax spp.*; >900), Atlantic Puffin (*Fratercula arctica*; >200), Black-legged Kittiwake (*Rissa tridactyla*; >200), Razorbill (*Alca torda*; >100), and terns (Fig. 1; Avery-Gomm et al., in prep). Notable mortality events were also reported in other aquatic species. This included Double-crested Cormorant (*Nannopterum auritum;* hundreds) in Alberta, Quebec, New Brunswick, and Nova Scotia in June 2022, in Eared Grebe (*Podiceps nigricollis;* hundreds) and Western Grebe (*Aechmophorus occidentalis; <100;* classified as special concern in Schedule 1 of the *Species at Risk Act* (36) in Alberta in June, and in multiple American White Pelican (<100, *Pelecanus erythrorhynchos*) colonies in the Mississippi and Pacific flyways in Canada in June and July 2022 (Fig. 1).

In fall 2022, notable mortality events were again reported in Geese. Canada Geese and Cackling Geese (*Branta hutchinsii)* mortalities were reported in September in the Pacific flyway (British Columbia) and in Snow Geese in mid-November 2022 in the Atlantic (Quebec) and Pacific (British Columbia) flyways. It is important to note that, in all cases, the reported wild bird mortality numbers will represent only a fraction of the total mortality.

### Morbidity and Mortality Wild Bird Surveillance Component

A total of 6,246 sick and dead wild birds were collected and tested for the presence of influenza A genomic material across Canada from November 2021 to November 2022 (Fig. 2A). Overall, 1,710 (27.4%; 95% confidence interval (CI): 26.3 – 28.5%) were confirmed or suspect positive for HPAIV (Table 1). Unless otherwise indicated (Table S1), species that tested HPAIV positive based on pooled swab samples and that underwent gross and histologic examination, had a majority of individuals with characteristic degenerative and inflammatory lesions consistent with HPAIV infection. A total of 62 (1.0%; 95% CI: 0.8 – 1.3%) sick and dead wild birds were positive for LPAIV. LPAIV was detected in members of the Charadriiformes, Anseriformes, and Accipitriformes (Tables 1 and S1).

**Fig. 2.**
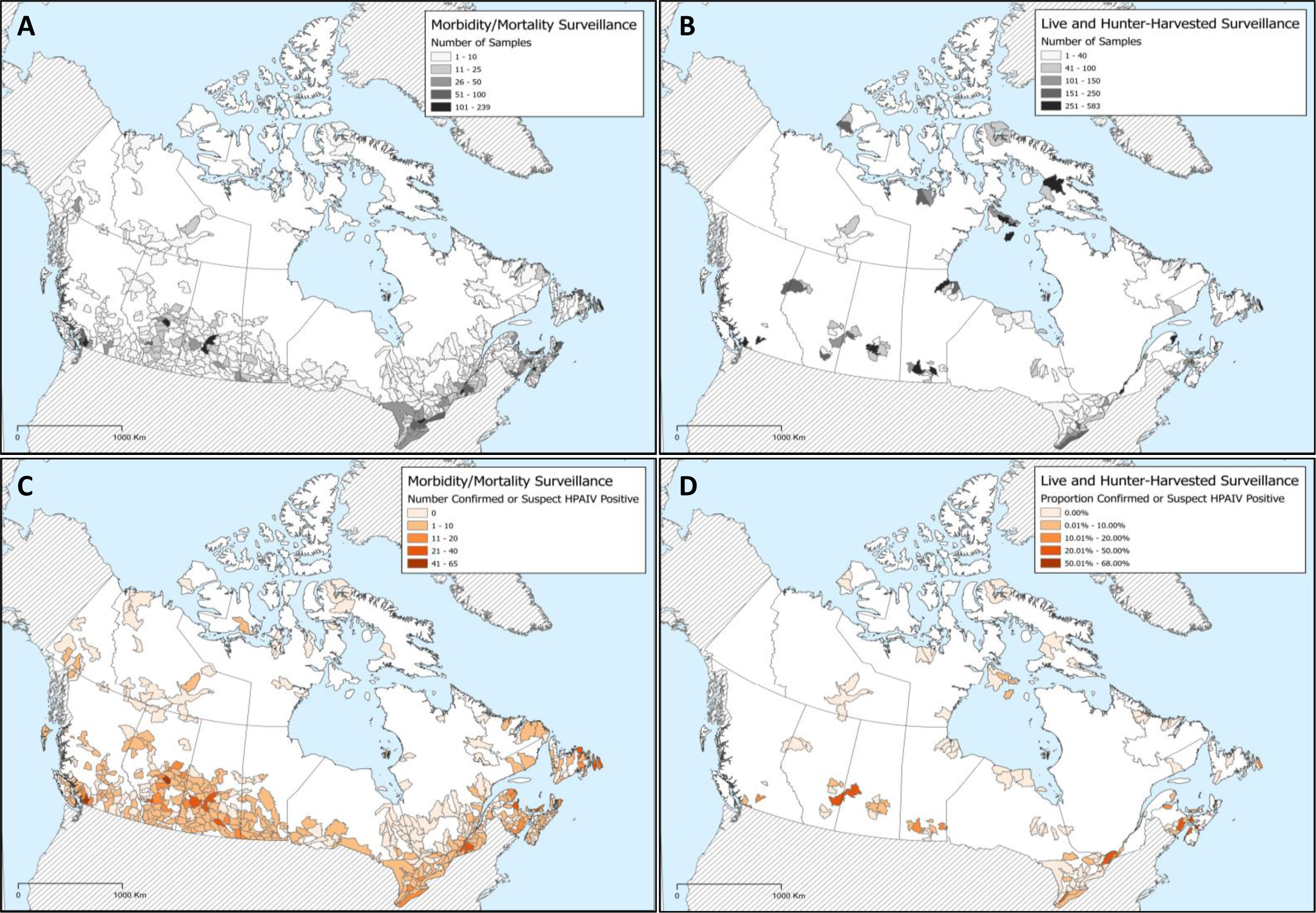
Distribution of A) sick and dead wild birds submitted and tested, B) live and hunter-harvested wild birds tested for avian influenza virus, C) the number of sick and dead wild birds confirmed or suspected to be positive for highly pathogenic avian influenza virus (HPAIV), and D) the proportion of live and hunter harvested wild bird samples confirmed or suspected to be positive for HPAIV, in Canada between November 2021 and December 2022, based on RT-PCR. Internal boundaries indicate watershed (i.e., sub-sub-drainage area) where samples were received for testing. Shapefile was downloaded from the National Hydro Network (73) on Oct 19, 2022 and was clipped to land boundary.

**Table 1.** Number of sick and dead wild birds submitted for testing and suspect or confirmed highly pathogenic avian influenza virus (HPAIV) or low pathogenicity avian influenza virus (LPAIV) positive in Canada between November 2021 – November 2022. Colour shading reflects general administrative migratory flyway routes (orange = Pacific, green = Central, blue = Mississippi, yellow = Atlantic).

| N suspect or confirmed positive/total sampled |  |  |  |  |  |  |  |  |  |  |  |  |  |  |
| --- | --- | --- | --- | --- | --- | --- | --- | --- | --- | --- | --- | --- | --- | --- |
| Taxonomic Order | British<br>Columbia | Yukon | Alberta | Saskatchewan | Northwest<br>Territories | Manitoba | Ontario | Nunavut | Quebec | New<br>Brunswick | Newfoundland<br>and Labrador | Nova<br>Scotia | Prince Edward<br>Island | Total |
| **HPAIV** |  |  |  |  |  |  |  |  |  |  |  |  |  |  |
| Accipitriformes | 44/164 | 2/14 | 33/63 | 48/84 | 0/1 | 33/46 | 43/135 | -- | 55/143 | 5/16 | 2/11 | 12/64 | 11/32 | 288/773 |
| Anseriformes | 114/192 | 4/7 | 78/134 | 85/125 | 0/9 | 20/32 | 89/306 | 0/26 | 160/288 | 8/58 | 9/38 | 21/105 | 1/6 | 589/1326 |
| Dabbling Ducks | 15/38 | 0/1 | 11/28 | 4/17 | -- | 6/13 | 15/56 | -- | 12/56 | 1/14 | 0/15 | 3/44 | 0/1 | 67/283 |
| Diving Ducks and Seaducks | 1/5 | -- | 0/7 | 0/6 | 0/4 | 1/1 | 4/30 | -- | 46/59 | 3/37 | 9/20 | 12/29 | 0/1 | 76/199 |
| Geese | 87/121 | 3/3 | 65/97 | 79/100 | 0/5 | 13/18 | 67/168 | 0/26 | 102/173 | 4/7 | 0/2 | 6/32 | 1/4 | 427/756 |
| Swans | 11/28 | 1/3 | 2/2 | 2/2 | -- | -- | 3/52 | -- | -- | -- | 0/1 | -- | -- | 19/88 |
| Apodiformes | 0/7 | -- | -- | -- | -- | -- | -- | -- | -- | -- | -- | -- | 0/4 | 0/11 |
| Caprimulgiformes | -- | -- | -- | -- | -- | -- | -- | -- | -- | -- | -- | 0/3 | -- | 0/3 |
| Charadriiformes | 4/58 | 0/5 | 10/47 | 2/25 | 1/6 | 0/2 | 6/74 | 2/4 | 44/77 | 27/77 | 87/241 | 20/220 | 4/14 | 207/850 |
| Columbiformes | 0/27 | -- | 0/10 | 2/94 | -- | 0/1 | 0/31 | -- | -- | 0/3 | 1/5 | 0/22 | 0/9 | 3/202 |
| Coraciiformes | -- | -- | 0/1 | -- | 0/1 | -- | 0/3 | -- | -- | -- | -- | 0/1 | -- | 0/6 |
| Falconiformes | 7/10 | 0/1 | 4/19 | 4/14 | 0/1 | 3/9 | 1/17 | 0/1 | 3/49 | 0/2 | 0/1 | 0/4 | -- | 22/128 |
| Galliformes | 0/2 | 0/1 | 0/4 | 0/5 | 0/2 | 0/1 | 0/29 | 0/1 | 2/13 | 0/16 | 2/13 | 1/49 | 0/7 | 5/143 |
| Gaviiformes | -- | -- | -- | 0/2 | 0/1 | 0/2 | 0/11 | -- | 0/7 | 0/1 | 0/4 | 0/10 | 0/1 | 0/39 |
| Gruiformes | 0/2 | -- | 0/4 | 0/1 | 0/3 | -- | -- | -- | -- | -- | -- | -- | 0/1 | 0/11 |
| Passeriformes | 10/96 | 1/28 | 31/190 | 71/328 | 0/20 | 26/81 | 7/261 | -- | 25/163 | 18/116 | 7/46 | 7/265 | 21/137 | 224/1731 |
| Pelecaniformes | 15/39 | -- | 6/12 | 3/6 | -- | 7/11 | 12/25 | -- | 1/8 | 0/2 | -- | 0/6 | 0/4 | 44/113 |
| Piciformes | 0/5 | 0/2 | 0/9 | 0/4 | -- | -- | 0/4 | -- | -- | 0/2 | -- | -- | -- | 0/26 |
| Podicipediformes | 1/3 | 0/1 | 27/35 | 2/6 | -- | 3/3 | 0/2 | -- | -- | -- | -- | -- | -- | 33/50 |
| Procellariiformes | -- | -- | -- | -- | -- | -- | -- | -- | -- | 0/4 | 2/30 | 0/34 | -- | 2/68 |
| Strigiformes | 21/98 | 0/10 | 50/88 | 29/59 | 0/3 | 9/16 | 9/62 | -- | 5/63 | 4/27 | 1/20 | 0/31 | 1/6 | 129/483 |
| Suliformes | 0/1 | -- | 19/23 | 9/14 | -- | 1/1 | 0/16 | -- | 23/57 | 21/23 | 50/79 | 16/39 | 25/30 | 164/283 |
| **Total** | **216/704** | **7/69** | **258/639** | **255/767** | **1/47** | **102/205** | **167/976** | **2/32** | **318/868** | **83/347** | **161/488** | **77/853** | **63/251** | **1710/6246** |

| LPAIV |  |  |  |  |  |  |  |  |  |  |  |  |  |  |
| --- | --- | --- | --- | --- | --- | --- | --- | --- | --- | --- | --- | --- | --- | --- |
| Accipitriformes | 2/164 | 0/14 | 3/63 | 2/84 | 0/1 | 1/46 | 1/135 | -- | 4/143 | 0/16 | 0/11 | 0/64 | 0/32 | 13/773 |
| Anseriformes | 0/192 | 0/7 | 1/134 | 4/125 | 0/9 | 0/32 | 2/306 | 0/26 | 3/288 | 3/58 | 0/38 | 0/105 | 0/6 | 13/1326 |
| Dabbling Ducks | 0/38 | 0/1 | 0/28 | 0/17 | -- | 0/13 | 1/56 | -- | 1/56 | 3/14 | 0/15 | 0/44 | 0/1 | 5/283 |
| Diving Ducks and Seaducks | 0/5 | -- | 1/7 | 0/6 | 0/4 | 0/1 | 0/30 | -- | 1/59 | 0/37 | 0/20 | 0/29 | 0/1 | 2/199 |
| Geese | 0/121 | 0/3 | 0/97 | 4/100 | 0/5 | 0/18 | 1/168 | 0/26 | 1/173 | 0/7 | 0/2 | 0/32 | 0/4 | 6/756 |
| Swans | 0/28 | 0/3 | 0/2 | 0/2 | -- | -- | 0/52 | -- | -- | -- | 0/1 | -- | -- | 0/88 |
| Apodiformes | 0/7 | -- | -- | -- | -- | -- | -- | -- | -- | -- | -- | -- | 0/4 | 0/11 |
| Caprimulgiformes | -- | -- | -- | -- | -- | -- | -- | -- | -- | -- | -- | 0/3 | -- | 0/3 |
| Charadriiformes | 0/58 | 0/5 | 3/47 | 3/25 | 0/6 | 0/2 | 6/74 | 0/4 | 3/77 | 1/77 | 3/241 | 1/220 | 1/14 | 21/850 |
| Columbiformes | 0/27 | -- | 0/10 | 1/94 | -- | 0/1 | 0/31 | -- | -- | 0/3 | 0/5 | 0/22 | 0/9 | 1/202 |
| Coraciiformes | -- | -- | 0/1 | -- | 0/1 | -- | 0/3 | -- | -- | -- | -- | 0/1 | -- | 0/6 |
| Falconiformes | 0/10 | 0/1 | 0/19 | 0/14 | 0/1 | 0/9 | 0/17 | 0/1 | 0/49 | 0/2 | 0/1 | 0/4 | -- | 0% |
| Galliformes | 0/2 | 0/1 | 0/4 | 0/5 | 0/2 | 0/1 | 0/29 | 0/1 | 0/13 | 0/16 | 0/13 | 0/49 | 0/7 | 0/143 |
| Gaviiformes | -- | -- | -- | 0/2 | 0/1 | 0/2 | 0/11 | -- | 0/7 | 0/1 | 0/4 | 0/10 | 0/1 | 0/39 |
| Gruiformes | 0/2 | -- | 1/4 | 0/1 | 0/3 | -- | -- | -- | -- | -- | -- | -- | 0/1 | 1/11 |
| Passeriformes | 0/96 | 0/28 | 1/190 | 0/328 | 0/20 | 1/81 | 1/261 | -- | 4/163 | 0/116 | 0/46 | 0/265 | 1/137 | 8/1731 |
| Pelecaniformes | 0/39 | -- | 0/12 | 0/6 | -- | 0/11 | 1/25 | -- | 0/8 | 0/2 | -- | 0/6 | 0/4 | 1/113 |
| Piciformes | 0/5 | 0/2 | 0/9 | 0/4 | -- | -- | 0/4 | -- | -- | 0/2 | -- | -- | -- | 0/26 |
| Podicipediformes | 0/3 | 0/1 | 0/35 | 0/6 | -- | 0/3 | 0/2 | -- | -- | -- | -- | -- | -- | 0/50 |
| Procellariiformes | -- | -- | -- | -- | -- | -- | -- | -- | -- | 0/4 | 1/30 | 0/34 | -- | 1/68 |
| Strigiformes | 0/98 | 0/10 | 0/88 | 1/59 | 0/3 | 0/16 | 0/62 | -- | 1/63 | 0/27 | 0/20 | 0/31 | 0/6 | 2/483 |
| Suliformes | 0/1 | -- | 0/23 | 0/14 | -- | 0/1 | 0/16 | -- | 0/57 | 0/23 | 1/79 | 0/39 | 0/30 | 1/283 |
| Total | 2/704 | 0/69 | 9/639 | 11/767 | 0/47 | 2/205 | 11/976 | 0/32 | 15/868 | 4/347 | 5/488 | 1/853 | 2/251 | 62/6246 |

#### Spatial

Sick and dead birds were submitted from all provinces and territories, with relatively fewer submissions and detections in sick and dead bird samples from northern regions of provinces or the Territories (i.e., northern Canada) (Fig. 2A and 2C). Suspect or confirmed HPAIV detections in sick and dead birds occurred across all flyways but were in the highest numbers in the

Atlantic (particularly Quebec) and Central flyways (particularly Alberta and Saskatchewan) (Table 1, Fig. 2C).

#### Temporal

Following the initial incursion in November 2021, carcass submissions and detections began increasing between January and February 2022 (Fig. 3A). At the national scale, the highest number of submissions was in April 2022 (Fig. 3A), but this varied by flyway with the number of submissions peaking earlier in February and March 2022 in the Atlantic flyway and again in June 2022 (Fig. S1). The highest number of HPAIV detections was also in April 2022 at the national scale. Within flyways, the number of detections in sick and dead birds also peaked in April in the Central and Mississippi flyways but peaked in May and June in the Pacific and Atlantic flyways, respectively, with a second small peak in the fall (September in the Mississippi flyway, September and October in the Central flyway, and November in the Pacific and Atlantic flyways) (Fig. S1).

**Fig. 3.**
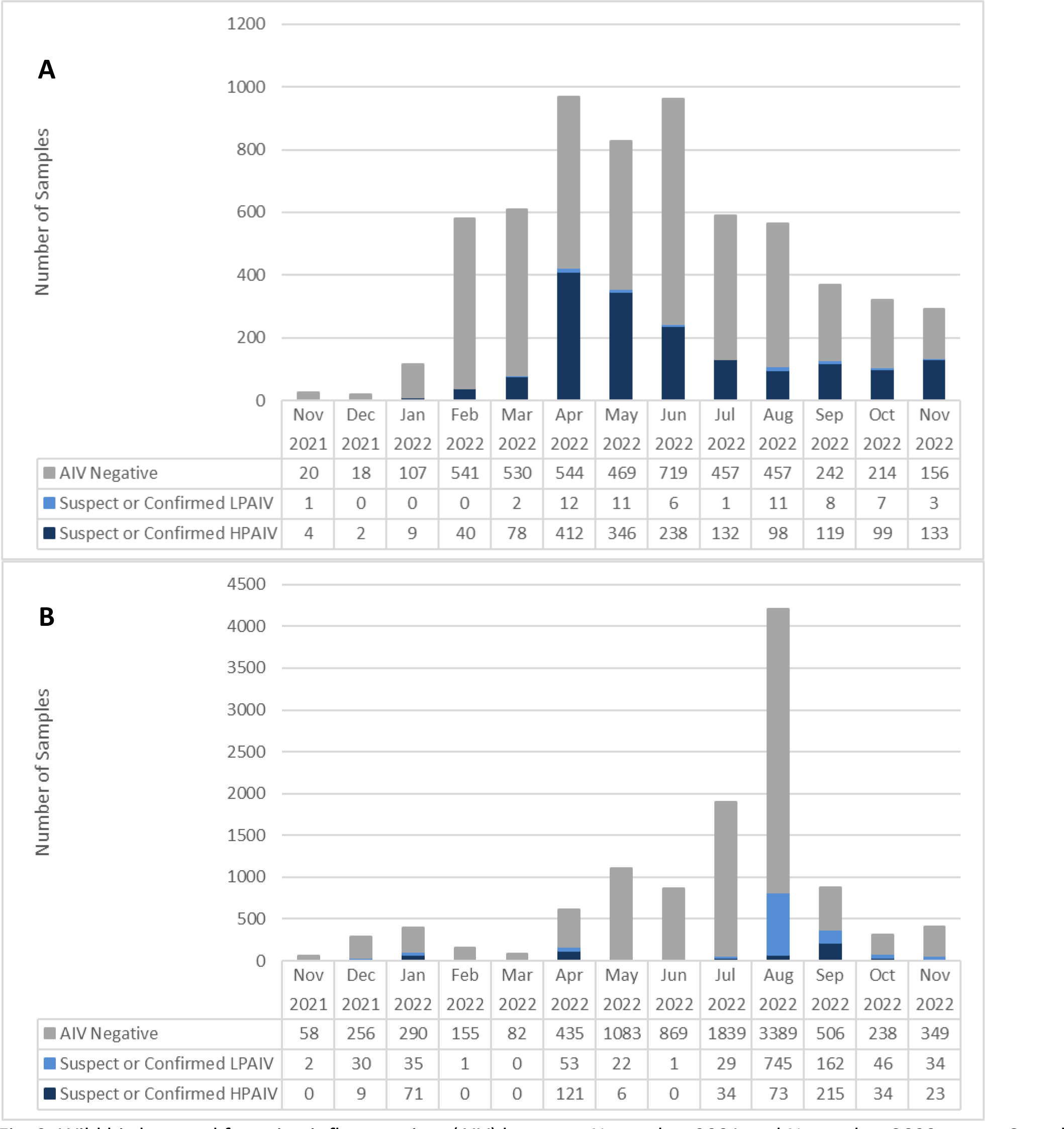
Wild birds tested for avian influenza virus (AIV) between November 2021 and November 2022 across Canada obtained through A) morbidity/mortality surveillance and B) live and hunter-harvested surveillance.

#### Taxonomic Order

Eighteen different taxonomic orders and 207 different species were screened through morbidity/mortality surveillance (Table S1). Fifty-five carcasses (0.9%) were only identified to the genus level (Table S1). HPAIV was confirmed or suspected in 12 taxonomic orders and 80 species (Fig. 4; Table S1). Taxonomic orders or functional groups with the largest number of detections included: Anseriformes (primarily geese, diving ducks and sea ducks, and dabbling ducks), raptors (i.e., Accipitriformes, Falconiformes, Strigiformes; primarily owls, hawks; eagles, and vultures), Passeriformes (primarily corvids), Charadriiformes (primarily gulls, terns, and murres), and Suliformes (primarily Northern Gannets and cormorants; Fig. 4; Table S1). Small numbers of suspect or confirmed HPAIV-positive Pelecaniformes (primarily in American White Pelicans) and Podicipediformes (primarily Western and Eared Grebes) were also detected (Fig. 4; Tables 1 and S1).

**Fig. 4.**
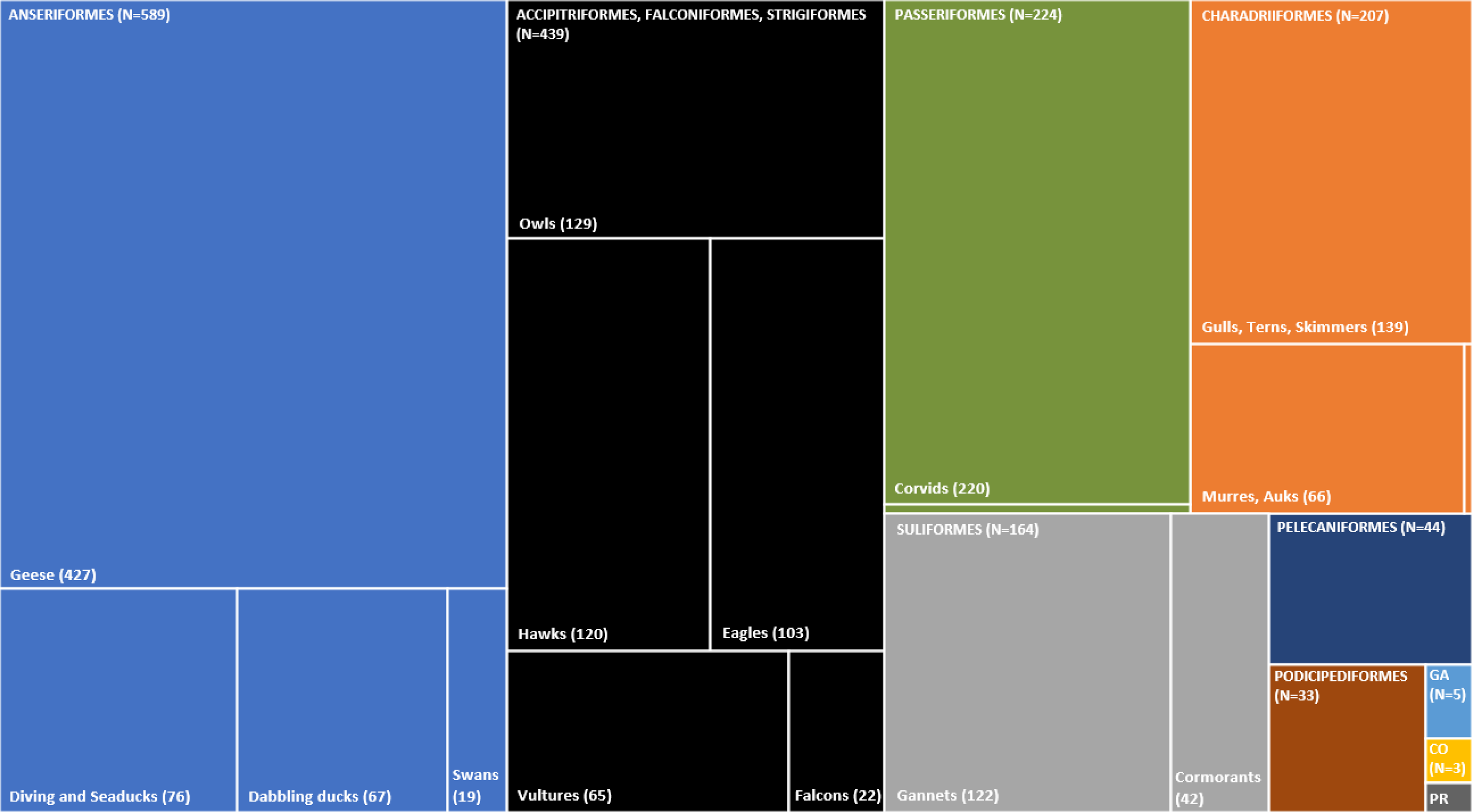
Breakdown of sick and dead wild bird samples that were confirmed or suspected highly pathogenic avian influenza virus (HPAIV) positive between November 2021 and November 2022 across Canada stratified by taxonomic order and species functional group. The data are shown as a treemap; the area of each rectangle is proportional to the number of wild bird samples that were confirmed or suspected HPAIV positive. (GA = Galliformes, CO = Columbiformes, PR = Procellariiformes (N=2), unlabeled Passeriformes = other (N=4), unlabeled Charadriiformes = Sandpipers and Allies (N=2)). Note: the data shown here reflect the samples that were tested and therefore do not represent the number of birds that died from HPAIV.

At the national scale and across the Pacific, Central and Mississippi flyways, sick and dead geese, primarily Snow Geese and Canada Geese, accounted for the most detections (Table 1). A first peak of detections in both Canada Geese and Snow Geese occurred in April 2022 (Fig. S3). A second peak in Canada Geese occurred in September 2022, although detections in this species occurred continuously from January through November 2022 across flyways. The second peak in Snow Geese occurred in November 2022 (Fig. S3); trends in Snow Geese were largely driven by detections in the Central and Atlantic flyways (Fig. S3).

At the national scale, peaks in morbidity and mortality for dabbling ducks occurred in the spring (April) and fall (September; Table 1). In diving ducks and seaducks, peaks corresponded with the breeding season in May and June and was largely driven by Common Eiders in eastern Canada (Fig. S4).

Peaks in morbidity and mortality for raptors and corvids occurred in the spring, in April and May, with a slight increase in detections in the fall for both functional groups (Fig. S5). The majority of HPAIV detections in corvids occurred in the Central, eastern Mississippi, and Atlantic flyways (Fig. S5). HPAIV detections in raptors occurred across all flyways (Fig. S5), however the majority of detections in Strigiformes were found in the Pacific and Central flyways (Table 1).

### Live and Hunter-Harvested Wild Bird Surveillance Component

A total of 11,295 live and hunter-harvested birds were tested for AIV across Canada between November 2021 and November 2022. Overall, 586 (5.2%; 95% CI: 4.8 – 5.6%) were confirmed or suspect positive for HPAIV (Table 2), and 1,160 (10.3%; 95% CI: 9.7 – 10.8%) were confirmed or suspect positive for LPAIV (Table 2). The following sections provide a more detailed breakdown of HPAIV and LPAIV detections in apparently healthy live or hunter-harvested wild birds.

**Table 2.**
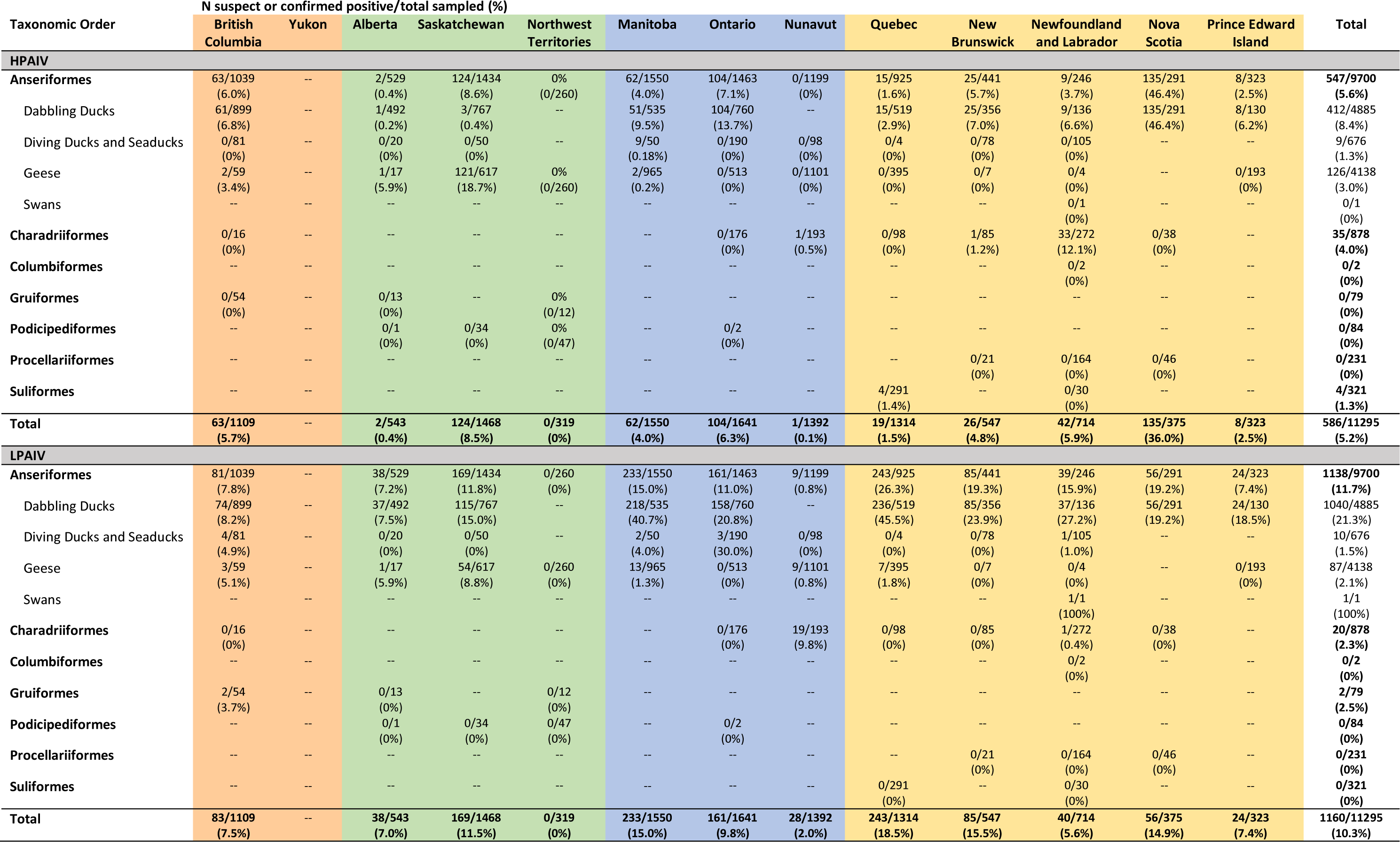
Number of live and hunter-harvested wild birds submitted for testing, and suspect or confirmed highly pathogenic avian influenza virus (HPAIV) or low pathogenicity avian influenza virus (LPAIV) positive in Canada between November 2021 – November 2022. Colour reflects migratory flyway routes (orange = Pacific, green = Central, blue = Mississippi, yellow = Atlantic).

#### Spatial

Samples from live and harvested birds were collected in all provinces and two territories (Fig. 2B). The Central and Atlantic flyways had the highest prevalence of HPAIV across multiple watersheds in which sampling occurred (Fig. 2D). In the northern portions of the flyways, prevalence of HPAIV in live and harvested birds ranged from 0% in Northwest Territories (n=319; 95% CI: 0 – 1.1%) to 0.07% in Nunavut (n=1392; 95% CI: <0.01 – 0.4%). In the southern Canadian portions of the Pacific, Central, Mississippi, and Atlantic flyways, apparent prevalences were 5.7% (n=1109; 95% CI: 4.4 – 7.2%), 6.3% (n=2011; 95% CI: 5.2 – 7.4%), 5.2% (n= 3191; 95% CI: 4.5 – 6.0%), and 7.0% (n=3273; 95% CI: 6.2 – 8.0%), respectively (Table 2, Fig. 2D). The highest proportion of confirmed and suspected HPAIV-positive samples occurred in Nova Scotia (Table 2). This high proportion was driven by two separate sampling events of Anseriformes; one that took place in January 2022 (68/100 HPAIV suspect or confirmed positive) in advance of but in close proximity (temporally and geographically) to infected premises (in Canada, defined as premises where HPAIV has been detected and confirmed through laboratory testing(37)) and the second in September 2022 (n=61/94).

In the northern portions of the flyways, LPAIV prevalence in apparently healthy birds ranged from 0% in the Northwest Territories (n=319; 95% CI: 0 – 1.1%) to 2.0% in Nunavut (n=1392; 95% CI: <1.3 – 2.9%), and appeared to increase from west to east across the southern Canadian portion of the flyways (Pacific: 7.5%, 95% CI: 6.0 – 9.2%; Central: 10.3%, 95% CI: 9.0 – 11.7%; Mississippi: 12.3%, 95% CI: 11.2 – 13.5%, and Atlantic: 13.7%, 95% CI: 12.5 – 14.9%; Table 2).

#### Temporal

At the national level and within each flyway, the majority of samples from live birds were collected in July and August (Fig. 3B; Fig. S2). There was a peak in prevalence of HPAIV in apparently healthy birds in January (17.9%; 95% CI: 14.3 – 22.1%), largely driven by sampling of mallards at a single open water pond in Nova Scotia in proximity to an infected premises (Atlantic flyway; 54.4%; 95% CI: 45.3 – 63.3%; Fig. S2D). A second peak in HPAIV prevalence in Canada occurred in April (19.9%; 95% CI: 16.8 – 23.3%) in association with spring migration (Fig. 3B), largely driven by the Central flyway (28.1%, 95% CI: 23.8 – 32.7%; Fig. S2B), and the highest peak in prevalence in Canada occurred in September 2022 (24.3%, 95% CI: 21.6 – 27.3%; Fig. 3B) during fall migration, largely in the Mississippi (26.2%; 95% CI: 22.1 – 30.6%) and Atlantic (22.9%; 95% CI: 18.8 – 27.4%) flyways, and to a lesser extent the Pacific flyway (57.1%, but note small sample size; 95% CI: 30.4 – 78.2%) (Fig. S2). There was no live bird surveillance in the Central flyway until April 2022, when the HPAIV prevalence was highest in that flyway (Fig. S2B).

LPAIV detections in apparently healthy birds were highest in August 2022 at the national scale, and prevalence peaked in August and September (17.7; 95% CI: 16.6 – 18.9% and 18.3%; 95% CI: 15.8 – 21.1%, respectively; Fig. 3B). LPAIV prevalence was highest in August in the Atlantic (29.7%; 95% CI: 27.0 – 32.5%) flyway, in September in the Central flyway (17.4%; 95% CI: 7.8 – 31.4%), and from August to October in the Mississippi flyway (17.2%; 95% CI: 15.4 – 19.2% to 21.8%; 95% CI: 15.6 – 29.1%; Fig. S2), largely driven by trends observed in Anseriformes, which had the highest LPAIV prevalence within each flyway (Table 2).

#### Taxonomic Order

Seven different taxonomic orders and 59 species were screened through live and hunter-harvested bird surveillance (Table S1). Twenty-seven individuals were only identified to the genus level (0.2%). HPAIV was confirmed or suspected in apparently healthy birds of 19 species from three taxonomic orders, including Anseriformes (5.6%; 95% CI: 5.2 – 6.1%), Charadriiformes (4.0%; 95% CI: 2.8 – 5.5%) and Suliformes (1.2%; 95% CI: 0.3 – 3.2%; Tables 2 and S1).

Within live or hunter-harvested Anseriformes, dabbling ducks had the highest HPAIV prevalence (8.4%; 95% CI: 7.7 – 9.2%, Table 2), with the highest found in American Black Duck (*Anas rubripes*, 13.3%; 95% CI: 10.0 – 17.3%), Northern Pintail (*Anas acuta*, 11.4%; 95% CI: 7.2 –16.9%), and Mallard (10.0%; 95% CI: 8.9 – 11.3%; Table S1). Lower HPAIV prevalences were found in Blue-Winged Teal (*Spatula discors*, 2.2%; 95% CI: 1.3 – 3.5%) and Green-Winged Teal (7.3%; 95% CI: 5.2 – 9.9%). Overall, 3.0% (95% CI: 2.5 – 3.6%) of apparently healthy geese were suspected or confirmed positive for HPAIV (Table 2). Out of 1,427 live or harvested Canada Geese across the country, only one sample was suspected or confirmed HPAIV positive in September 2022 in the Central flyway (Table S1). In contrast, out of 2,475 Snow Geese tested, 125 (5.1%; 95% CI: 4.2 – 6.0%) were suspected or confirmed positive for HPAIV, with the highest peak in prevalence detected in April 2022 (24.2%; 95% CI: 20.5 – 28.3%), largely in the Central flyway (Fig. S3, Table S1). Overall, 1.3% (95% CI: 0.6 – 2.5%) of apparently healthy diving ducks and sea ducks were suspected or confirmed positive for HPAIV (Table 2), with positives found in Canvasback (*Aythya valisineria*, 19.4%; 95% CI: 8.2 – 36.0%) and Redhead (*Aythya americana*, 2.0%; 95% CI: 0.2 – 7.0%; Table S1). All 272 Common Eiders, sampled in New Brunswick, Newfoundland, and Nunavut, tested negative for HPAIV (Table S1).

Within live or hunter-harvested Charadriiformes, the highest HPAIV prevalence was in Common Murre (61.9%; 95% CI: 45.6 – 76.4%; Table S1), of which 41 were sampled from a single colony in the Atlantic flyway and 31 were sampled over the course of three days during an active outbreak. In contrast, only one Thick-Billed Murre (*Uria lomvia*) sampled in Nunavut was suspected positive for HPAIV of 174 sampled nationally during the study period. Overall, of 397 gulls and terns (family Laridae) sampled, seven (1.8%; 95% CI: 0.7 – 3.6%) were confirmed or suspect positive for HPAIV, with positives found in only Black-Legged Kittiwake (12.8%; 95% CI: 4.3 – 27.4%) and Herring Gull (3.0%; 95% CI: 0.4 – 10.5%; Table S2). Of 132 apparently healthy shorebirds and waders sampled (families Charadriidae and Scolopacidae), all were negative for HPAIV (Table S1).

Within Suliformes, four of 321 apparently healthy Northern Gannets (1.3%; 95% CI: 0.3 – 3.2%) were confirmed or suspect positive for HPAIV (Table S1).

LPAIV was detected in three of the seven taxonomic orders sampled, including Anseriformes (11.7%; 95% CI: 11.1 – 12.4%), Charadriiformes (2.3%; 95% CI: 1.4 – 3.5%), and Gruiformes (2.5%; 95% CI: 0.3 – 8.8%; Table 2). Within Anseriformes, apparently healthy dabbling ducks had the highest LPAIV prevalence (21.3%; 95% CI: 20.1 – 22.5%; Table 2), with the highest found in American Black Duck (30.6%; 95% CI: 25.8 – 35.6%), Blue-Winged Teal (28.5%; 95% CI: 25.4 – 31.7%), Green-Winged Teal (20.7%; 95% CI: 17.2 – 24.4%), and Mallard (19.1%; 95% CI: 17.6 – 20.6%; Table S1). High LPAIV prevalence was observed in Northern Shoveler (41.7%; 95% CI: 15.2 – 72.3%), but only 12 individuals were sampled from this species (Table S1). Among the live and harvested geese sampled, 2.1% (95% CI: 1.7 – 2.6%) were positive for LPAIV, and most positives were found in Snow Geese (3.4%; 95% CI: 2.7 – 4.2%), with a few found in Canada Geese (0.2%; 95% CI: 0.04 – 0.6%) (Table S1). Only 1.5% (95% CI: 0.7 – 2.7%) of diving ducks or sea ducks were positive for LPAIV amongst several species sampled (Table 2 and S1). Among the apparently healthy Charadriiformes sampled, LPAIV was detected in only Thick-Billed Murre (19 of 174 tested; 10.9%; 95% CI: 6.7 – 16.5%) and one of 200 Ring-Billed Gulls sampled (0.5%; 95% CI: 0.01 – 2.8%; Table S1). All 132 shorebirds and waders were negative for LPAIV. Within the apparently healthy Gruiformes sampled, two of 64 American Coots (*Fulica americana*, 3.1%; 95% CI: 0.4 – 10.8%) were positive for LPAIV, and none of the 15 Whooping Cranes (*Grus americana*, classified as endangered in Schedule 1 of the *Species at Risk Act* (36) were positive for LPAIV or HPAIV (Table S1).

### Viral Reassortment and Phylogenetic Analysis

There was substantial genetic diversity in HPAIV viruses, as 341 (24.8%) were Eurasian AIVs with the remaining resulting from reassortments between Eurasian and North American viruses, with evidence of 10 different genome constellations (Table 3, Figs. 6 and 7). Identification of unique genome constellations was based on the pattern of monophyletic clades from individual gene segments (Signore et al., in prep.). The majority of reassortants detected involved one to four of the five internal gene segments (i.e., PB2, PB1, PA, NP, and NS) originating from North American LPAIVs, and none involved reassortment of the HA gene. Only a single virus (Pattern 10) in an apparently healthy Blue-Winged Teal in Manitoba in August 2022 involved reassortment of the NA and M genes. However, while this H5N6 virus showed reassortment involving all but the HA gene, it was sequenced directly from the swab material from this bird and virus isolation was unsuccessful. In swabs collected from birds at the same location during the same month, virus isolation yielded either Eurasian H5N1 or North American H4N6 virus. The most common gene segments involved in reassortment were NP, PB2, and PB1, which were involved in eight, seven, and five of the 10 genome constellation patterns, respectively (Table 3). The most common genome constellations detected in Canada in the first year since incursion included the Eurasian lineage along with Patterns 2, 4, and 5, collectively comprising 93.5% of all sequenced viruses (Table 3).

**Fig. 5.**
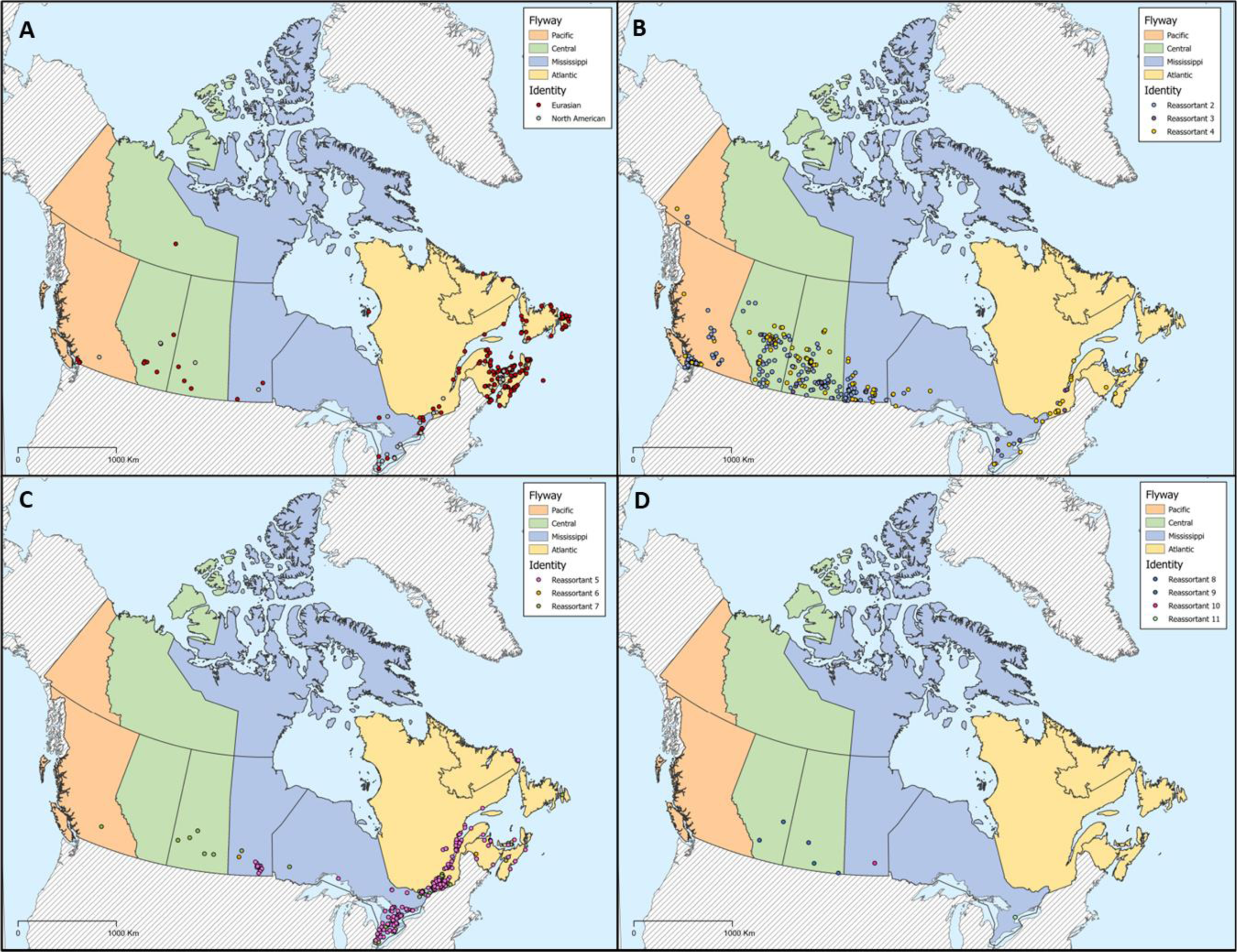
Distribution of avian influenza viruses characterized from wild birds sampled during the first year of the outbreak (November 2021 – November 2022) across Canada. Viruses detected included those with A) fully Eurasian or North American origins, and B)-D) and ten reassortant viruses, the genetic composition of which are described in Table 3. Provinces and territories are colored by the predominant migratory bird flyway.

**Fig. 6.**
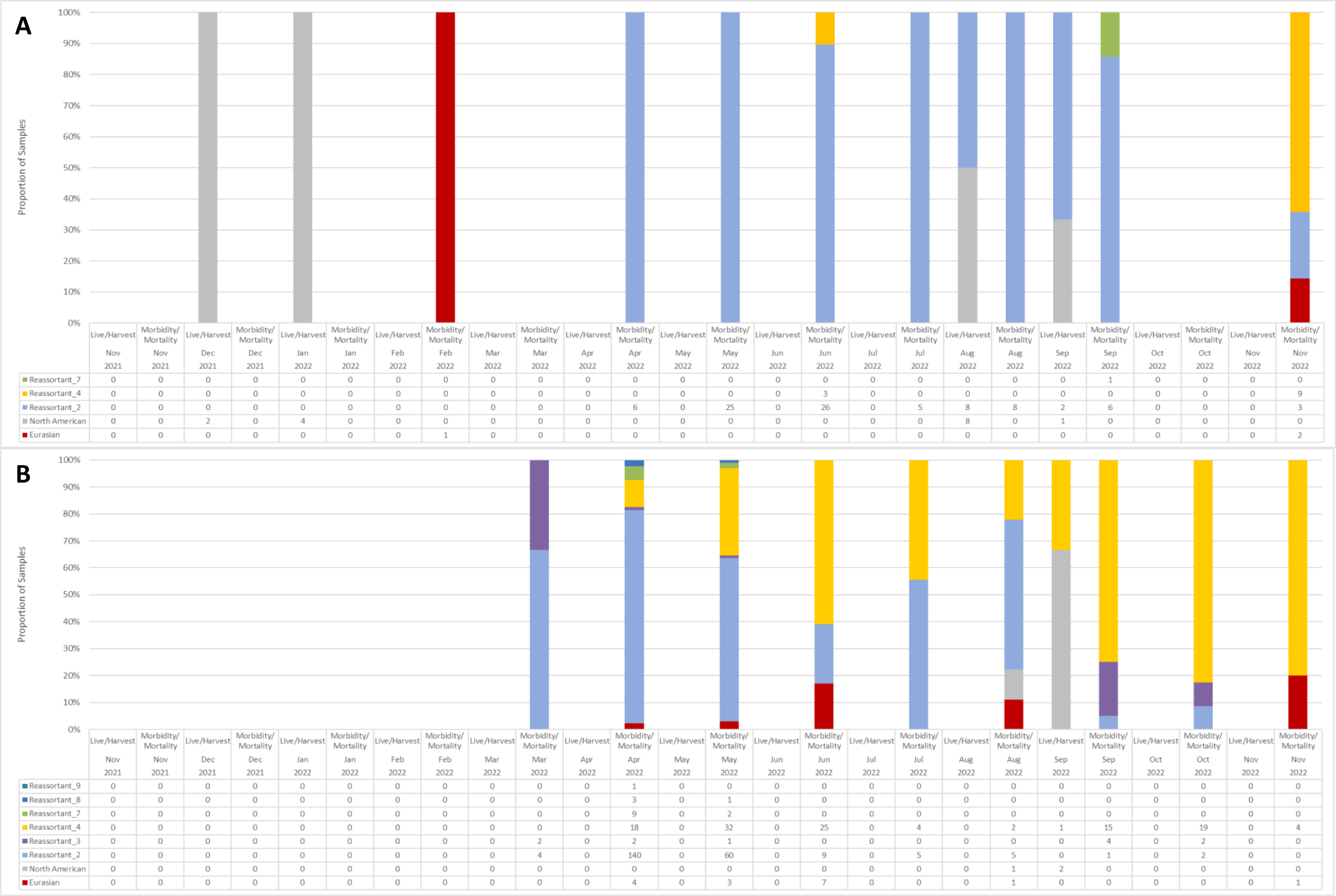
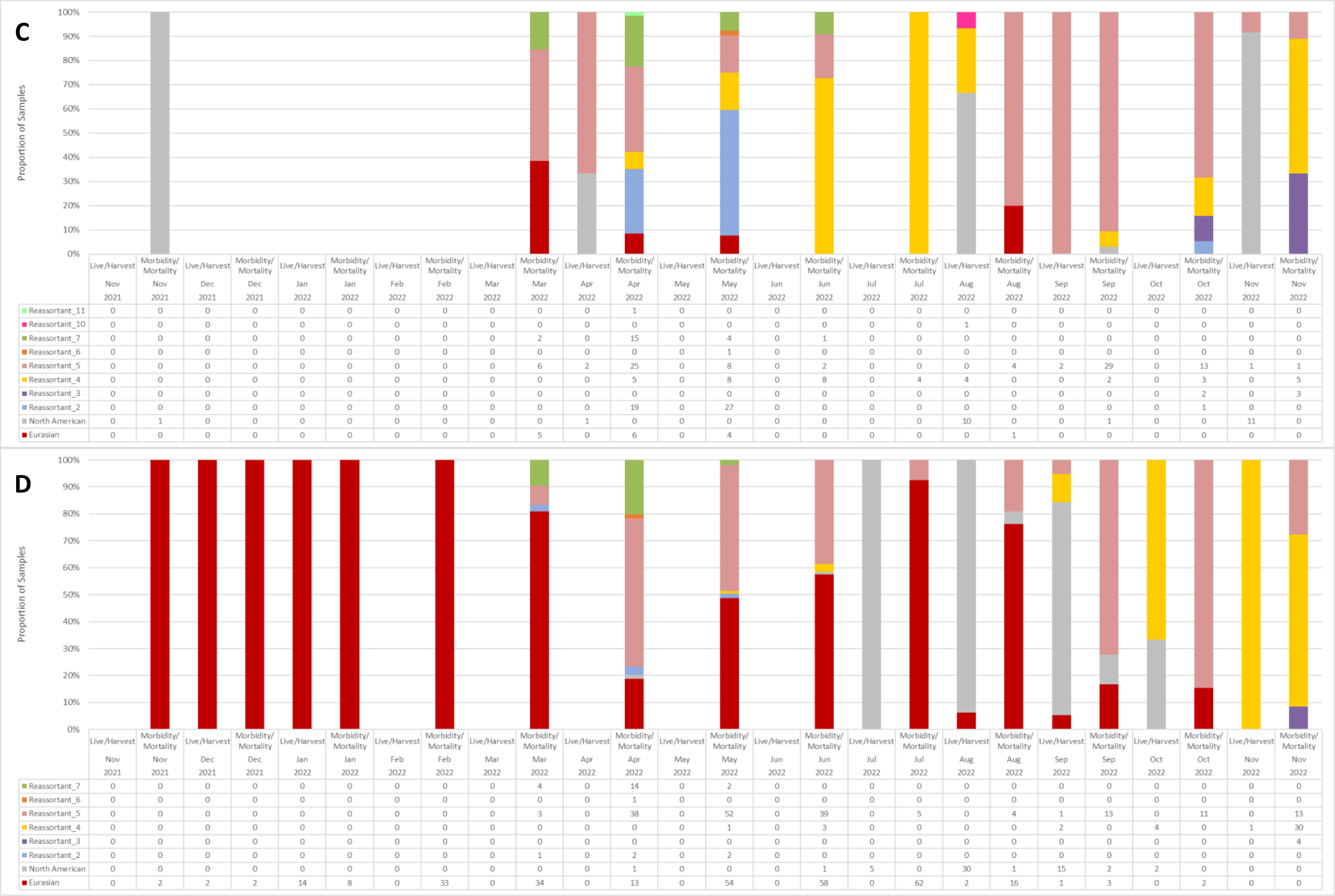
Avian influenza viruses characterized from wild birds sampled during the first year of the outbreak (November 2021 – November 2022) in the A) Pacific flyway, B) Central flyway, C) Mississippi flyway, and D) Atlantic flyway, stratified by surveillance component, and time. Viruses detected included both Eurasian, North American, and ten reassortant viruses the genetic composition of which are described in Table 3.

**Fig. 7.**
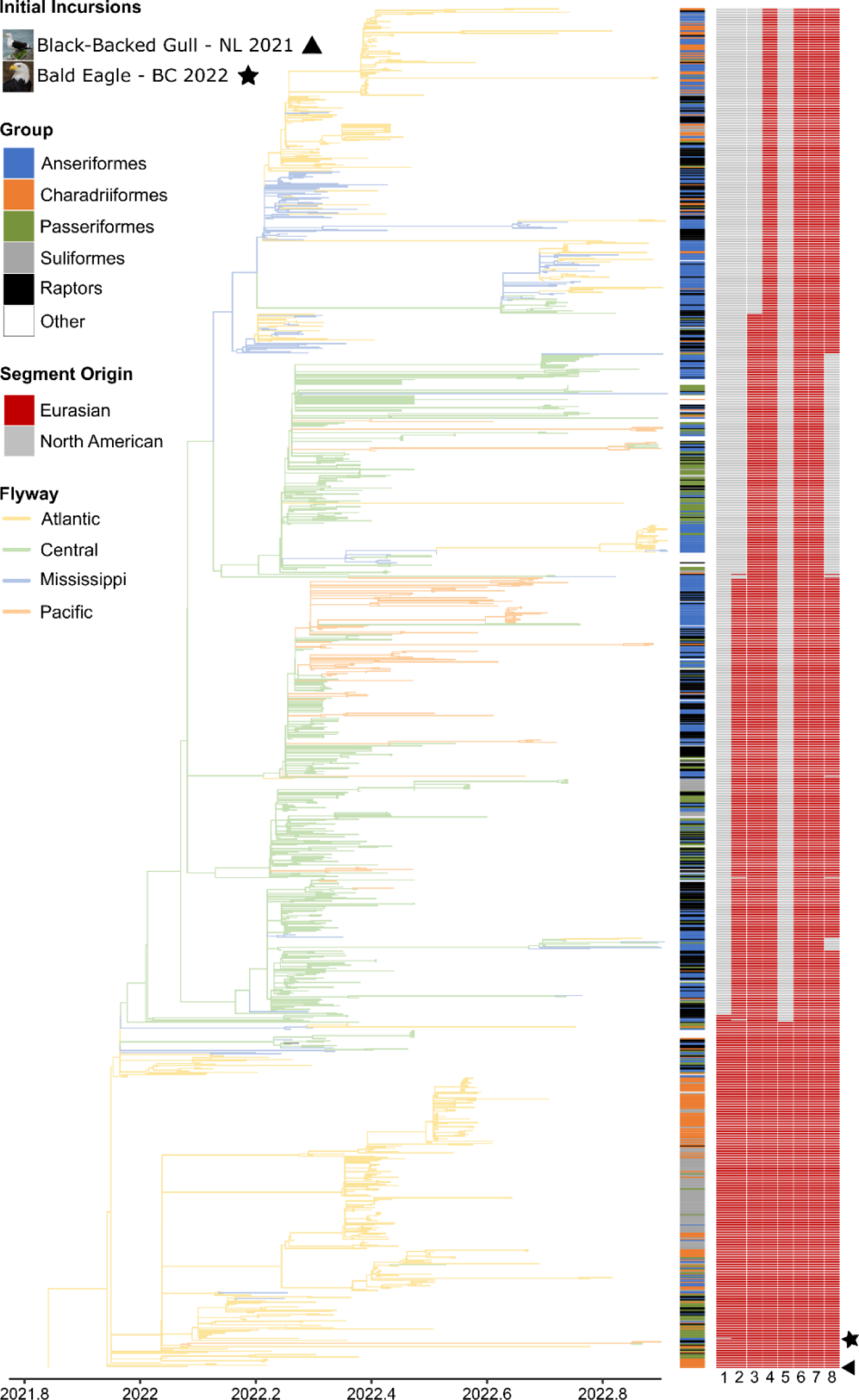
Time-calibrated maximum likelihood phylogenetic tree of 1,166 HPAIV H5N1 complete viral genomes collected from wild bird hosts between November 2021 – 2022. Tree branches are coloured according to the flyway in which the sample was collected. Host taxonomic order is represented by coloured bars at tree tips. The geographic origin of genome segments 1 – 8 (either Eurasian or North American) for all tree tips are represented by red and grey bars (from left to right: PB2, PB1, PA, HA, NP, NA, M, NS). First H5N1 2.3.4.4b detections following the initial incursions into Canada are annotated.

**Table 3.** The geographic origin of avian influenza virus genome segments (Eurasian – EA or North American – N Am) for genome constellations detected through wild bird surveillance in Canada between November 2021 – November 2022.

| Genome Constellation | n | Gene |  |  |  |  |  |  |  |
| --- | --- | --- | --- | --- | --- | --- | --- | --- | --- |
|  |  | PB2 | PB1 | PA | HA | NP | NA | M | NS |
| Eurasian | 341 | EA | EA | EA | EA | EA | EA | EA | EA |
| Pattern 2 | 367 | N Am | EA | EA | EA | N Am | EA | EA | EA |
| Pattern 3 | 20 | N Am | EA | EA | EA | N Am | EA | EA | N Am |
| Pattern 4 | 212 | N Am | N Am | EA | EA | N Am | EA | EA | N Am |
| Pattern 5 | 272 | N Am | N Am | N Am | EA | N Am | EA | EA | EA |
| Pattern 6 | 2 | N Am | EA | EA | EA | EA | EA | EA | EA |
| Pattern 7 | 54 | N Am | N Am | EA | EA | N Am | EA | EA | EA |
| Pattern 8 | 4 | EA | EA | EA | EA | N Am | EA | EA | EA |
| Pattern 9 | 1 | EA | N Am | EA | EA | N Am | EA | EA | EA |
| Pattern 10 | 1 | N Am | N Am | N Am | EA | N Am | N Am | N Am | N Am |
| Pattern 11 | 1 | EA | EA | N Am | EA | EA | EA | EA | EA |
| North American | 99 | N Am | N Am | N Am | N Am | N Am | N Am | N Am | N Am |

Detections of Eurasian-origin virus occurred throughout the full time period and in every flyway, but most Eurasian HPAIV in Canada occurred in sick and dead birds in the Atlantic flyway (Figs. 5, 6, and 7). Only a small proportion of viruses sequenced from live and harvested birds were Eurasian, and all were detected in the Atlantic flyway in August and September 2022 (Fig. 6). Following the first detection of reassortment in March 2022 in the Atlantic flyway, the proportion of Eurasian viruses detected decreased in April, increased again every month until July when the majority of detections were Eurasian virus, and then decreased again to zero by November 2022 (Fig. 6). Charadriiformes and Suliformes hosts made up the greatest proportion of Eurasian virus detections. The Eurasian virus, more so than reassortant viruses, showed phylogenetic segregation by host group, as viruses sequenced from Suliformes tended to be genetically distinct from those sequenced from Charadriiformes (Fig. 7).

The most commonly detected genome constellation overall was Pattern 2 (n=367), which is the most closely related to the Eurasian virus of the main genome constellations detected. Both Pattern 2 and the Eurasian H5N1 were distributed across Canada, however Pattern 2 was widely distributed in western Canada, as it was the most common genome constellation found in the Pacific and Central flyways (Figs. 5 and 6). Pattern 2 (including viruses from the Pacific flyway referenced as clusters 1, 2, 4, and 5 in Andrew et al., submitted) was first detected in the Central and Atlantic flyways in March 2022, and in April 2022 in the Mississippi and Pacific flyways (Fig. 6). Pattern 2 detections decreased to zero in both the Atlantic and Mississippi flyways by June 2022; however, they remained prominent in the Central and Pacific flyways (Fig. 6). Like the Eurasian virus, Pattern 2 showed genetic distinctions by flyway, as viruses collected from the Central flyway cluster separately from those from the Pacific flyway (Fig. 7). However, unlike the Eurasian virus, there is limited phylogenetic structure by host, as viruses from each host group form much smaller monophyletic groups than those infected with the Eurasian virus (Fig. 7).

Pattern 4 (including viruses from the Pacific flyway referenced as cluster 3 in Andrew et al., submitted) was among the most commonly detected genome constellations across Canada (n=212). This reassortant, which is one of the most divergent genome constellations from the Eurasian virus (Fig. 7), was first detected in April 2022 in adjacent Central and Mississippi flyways and represented a high proportion of detections particularly in the Central flyway through to November 2022. Pattern 4 was detected in the Atlantic flyway in a small proportion of samples starting in May but became dominant by October in live and harvested birds, and by November 2022 in sick and dead birds. A small number of viruses with Pattern 4 were detected in early June 2022 (note: Andrew et al., submitted report these as occurring in late May 2022) in the Pacific flyway and then again in November 2022 in the majority of collected samples (Fig. 6). Like the other reassortants, Pattern 4 also showed phylogenetic segregation by flyway, but also included a large cluster of viruses that almost exclusively infected Anseriformes in the Pacific flyway.

Pattern 5, which was among the most commonly detected genome constellation patterns (n=272), was only found in the Mississippi and Atlantic flyways, and continued to be detected through to November 2022 in high proportions in both flyways, including in live and harvested birds (Table 3; Fig. 6). It was the most commonly found genome constellation in the Mississippi flyway overall and peaked in the spring (March to April) and fall (September to October) of 2022 (Fig. 6). In the Atlantic flyway, it was second only to the Eurasian H5N1 virus, peaking in spring (April to June) and in fall (September to November), often outnumbering the Eurasian H5N1 within those time periods (Fig. 6).

The remaining reassortant patterns (3 and 6-11) were relatively uncommon in Canada, representing only 6.5% of sequenced viruses (Table 3, Figs. 6 and 7).

The highest peaks in diversity of genome constellation detections in a given flyway (e.g., >4 patterns) occurred in April and May in the Central, Mississippi, and Atlantic flyways, with additional increases in the number of patterns relative to preceding months in the fall (October and November) in the Pacific and Mississippi flyways (Fig. 6). These peaks in diversity often coincided with peaks in total numbers of detections (Fig. 6).

North American LPAIVs (n=99) were detected across all flyways, but the majority were detected in August and September 2022 in live and harvested birds in the Atlantic flyway (Fig. 5 and 6).

To date, a total of 28 different LPAIV subtypes (i.e., HxNx combinations) were detected in wild birds during the 2021-2022 outbreak event including H2N3, H2N9, H3N2, H3N6, H3N8, H4N2, H4N6, H4N8, H4N9, H5N2, H6N4, H6N8, H7N3, H7N4, H7N5, H7N7, H7N8, H9N2, H9N4, H9N8, H10N7, H11N2, H11N3, H11N9, H12N5, H12N6, H13N6, and H16N3.

## Discussion

The incursion of H5N1 HPAIV of clade 2.3.4.4b into Canada resulted in unprecedented detections in asymptomatic wild birds and large-scale wild bird mortality, affecting a wide range of species. Based on the authors’ collective knowledge, no other infectious disease, including the previous HPAIV incursion in 2014-15, has caused this magnitude of mortality in such a large diversity of bird species in Canada. While not the manuscript’s primary focus, characteristic lesions were associated with HPAIV-positive PCR results in most wild birds that underwent gross and histopathological examination, supporting the assumption that the majority of HPAIV-positive dead birds died as a result of infection. However, because we rely heavily on opportunistic reporting of mortality events, the data presented here provide only conservative estimates of the scope and scale of HPAIV-associated wild bird mortality in Canada. The field capacity and resources necessary to complete structured surveys are prohibitive during multiple and large-scale mortality events occurring simultaneously across the country.

### Wild Birds: Victims and Reservoirs

In the first year of the 2021-2022 outbreak, H5N1 HPAIV was detected in 1,710 sick or dead wild birds from 80 species across 12 taxonomic orders. Most of the sick and dead HPAIV-positive birds submitted for testing were Anseriformes (primarily geese) and raptors (owls, hawks, eagles), followed by corvids and Charadriiformes (primarily gulls and murres). However, the largest recorded mortality events, during which only a subset of carcasses get submitted for testing, occurred in Canada Geese and Snow Geese, and during the breeding season in colonial nesting species in eastern Canada, including Northern Gannet, Common Murre, and Common Eider. Where wild bird mortality was notable throughout the first year of the 2021-2022 outbreak with a wide taxonomic distribution, distinct peaks in detections among asymptomatic wild birds were observed in the spring and fall primarily in dabbling ducks. Peaks in AIV prevalence in dabbling ducks during spring and fall have been well-described for LPAIVs (38, 39) and correspond with northward wild bird migration and southward migration in conjunction with an influx of naïve juveniles, respectively. Increased abundance and density of wild migratory birds during migration facilitates viral transmission through close contact and environmental contamination.

Anseriformes and Charadriiformes, in particular, have been recognized as reservoirs for LPAIVs (40, 41). Our data from Canada along with other studies (9), indicate that candidate wild bird species may also act as reservoirs for H5N1 HPAIV. Amongst apparently healthy birds sampled through live or hunter-harvested bird surveillance, dabbling ducks had the highest prevalence of both HPAIV (8.4%; specifically American Black Duck, Mallard, their hybrids, and Northern Pintail) and LPAIV (21.3%). This was not surprising for LPAIV, as dabbling ducks have one of the highest LPAIV prevalence worldwide (41), and the largest number and diversity of AIV subtypes isolated globally (42). There were also no large-scale mortalities and few HPAIV detections in sick and dead dabbling ducks, corroborating that these species exhibit less morbidity and mortality compared to other Anseriformes (43). Exposure to LPAIVs has the potential to provide some level of heterosubtypic cross-protective immunity against HPAIV (44). Captive studies have demonstrated that pre-exposure to specific LPAIVs can confer partial cross-protective heterosubtypic immunity to other LPAIVs (45–47), as well as HPAIVs (44, 48, 49), which can result in reduced viral loads, duration of shedding, and, in the case of HPAIV, reduced morbidity and mortality. With their high prevalence of both HPAIV and LPAIV, together with their large population sizes (at least 25 million dabbling ducks; USFWS 2022), dabbling ducks are likely candidates as reservoirs of HPAIV in Canada.

Conversely amongst Anseriformes, geese, diving ducks and sea ducks appeared to be highly susceptible to morbidity and mortality. High numbers of sick or dead Canada Geese and Snow Geese tested positive for HPAIV, and HPAIV detections in apparently healthy geese were low compared to dabbling ducks, with only one Canada Goose testing positive out of 1,427 tested, and 125 (5.1%) Snow Geese positive out of 2,475 tested. Large mortality events were also reported in Common Eider. Of 96 Common Eider tested through sick and dead bird surveillance, 60 were HPAIV-positive, however none tested positive for HPAIV through live bird surveillance. Thus, Canada Geese and Common Eider do not appear to be strong candidates as reservoir species and have the potential to be significantly impacted by this virus if population level immunity does not develop or is not sustained over time. Conversely, Snow Geese have the potential to play an important role as sources of transmission and spread, particularly given their large population sizes (e.g., the mid-continental Snow Goose population was estimated at over 16.2 million (+/-1.6M) in 2022; USFWS, 2022), their gregarious behaviour during migration and breeding, the long distances traveled during migration which include arctic breeding grounds (i.e., potential areas of flyway overlap, including with trans-Atlantic migrants), and the significant overlap in ranges and habitats with other waterfowl species including dabbling ducks.

Charadriiformes have been proposed as potential candidates for the spread of AIV within and between colonies or foraging sites during the breeding season (50, 51), as well as over long distances during migration (4). Gulls, including several species identified as candidates for the movement of HPAIV from Europe to Canada in 2021 (2, 4), were the first cases detected in Canada and were often the index cases detected as the virus moved north. However, the contribution of Charadriiformes as a reservoir species is less clear than for dabbling ducks. While several species of Charadriiformes, particularly murres, gulls, and terns, were susceptible to HPAIV-related mortality, HPAIV detections through live and hunter-harvested surveillance were generally low. Notably, a self-limiting HPAIV outbreak with low mortality was reported in summer 2022 among Herring Gulls on Kent Island, New Brunswick, Canada (52). Similarly, a large proportion of apparently healthy Common Murre tested positive for HPAIV within a short time frame from a single colony experiencing an active outbreak (McLaughlin et al., submitted), yet no mortalities were observed based on re-sightings days later (J. Cunningham, personal communication). In contrast, Common Murre mortality events were reported at several other colonies in the Atlantic region during the same time period (Avery-Gomm et al., in prep), suggesting colony-level differences in virus dynamics and susceptibility.

Beyond gulls and Common Murre, certain seabird species, such as the Northern Gannet, exhibited remarkable susceptibility to HPAIV. Large-scale mortality was observed in a number of colonial nesting seabird species during the breeding season in Atlantic Canada and across the North Atlantic (53). Colonial nesting behaviour, characterized by dense populations, a high degree of social interaction, and shared foraging areas, can facilitate extensive transmission among conspecifics, leading to focal and large-scale die-offs following introduction of highly transmissible pathogens like HPAIV (54). However, there is evidence of exposure and survival in some of these highly susceptible colonial nesting species (53). While an in-depth analysis of the impacts to seabirds in Atlantic Canada is the focus of Avery-Gomm et al., in prep and falls outside the scope of this manuscript, we wish to emphasize the importance of continued targeted AIV surveillance in these populations in order to understand the interacting mechanisms driving species- and colony-level differences in virus dynamics, transmission, and susceptibility. Ongoing serologic surveillance can also contribute to expanding our knowledge of exposure and survival in these populations, advancing our understanding of heterosubtypic immunity, and enhancing our ability to forecast mass mortality following exposure to H5Nx HPAIVs.

Podicipediformes, or grebes, are diving waterbirds with previously demonstrated susceptibility to HPAIV (55), as corroborated by the current results. However, there is little evidence that Podicipediformes play an important role as LPAIV or HPAIV reservoir hosts with the caveats that these species are generally not well-studied. In our surveillance, Horned grebes (*Podiceps auritus*) were the only grebe species for which apparently healthy individuals were sampled. Among these individuals, there were no detections of HPAIV or LPAIV. Although there were relatively few samples (n=83), they were collected during the same month (June) and in the same flyway (Central) as reported grebe mortality, which was primarily observed in Eared (*P. nigricollis*) and Western grebes (*Aechmophorus occidentalis*). These colonial species have similar habitat preferences to dabbling ducks, especially Eared grebe, which rely on large shallow ponds with dense vegetation during breeding (56). Therefore, in areas of range overlap, there is an increased likelihood of exposure to HPAIV-contaminated habitat coupled with increased risk of transmission related to colonial nesting dynamics. In comparison, there was no reported HPAIV related mortality in Horned grebe, which may be less likely to occur in proximity to dabbling ducks, because they are highly territorial during breeding and are more likely to nest in isolation on smaller ponds with open water (57). As Podicipediformes are identified as priority species for conservation and stewardship in one or more locations in Canada (58) and two species appear in Schedule 1 of the *Species at Risk Act* (36), a better understanding of factors influencing HPAIV-related mortality events for birds in this order is warranted.

Raptors and corvids have demonstrated a pronounced susceptibility to HPAIV during the current outbreak and in the previous HPAIV outbreak in North America (59). The underlying reason is not known but is likely related to the route and dose of exposure. The most likely routes of exposure are through scavenging of infected carcasses and, in the case of raptors, through predation of infected prey (60) (Andrew et al., submitted). Infected prey that are displaying signs of weakness or abnormal behaviour may be preferentially targeted (61–63). While there are no samples in the national dataset from apparently healthy raptors and corvids, the role that these species play as reservoirs is likely to be minimal given that most species are relatively solitary. They may, however, play a role in subsequent spread to conspecifics at shared roosting or feeding sites, where some species can occur in high numbers, or to offspring during the breeding season (e.g., through parental feeding of infected prey items). The current outbreak reflects a significant shift in HPAIV dynamics, highlighting the dual role of wild birds as victims and reservoirs of this virus. Based on the data presented, there are differences in species susceptibility between and within wild bird taxonomic orders emphasizing the importance of a species-level approach to data interpretation and conclusions. The observed taxonomic and temporal patterns are also important to interpret in the context of a novel AIV, to which Canada’s migratory bird populations were immunologically naïve. Although widespread transmission should result in the development of immunity, and consequently reduced infection and mortality, the duration and extent of this immunity remains uncertain. Mortality events may continue to be pronounced in highly susceptible species, as a high case fatality rate may limit transmission and delay population-level immunity. Factors like food scarcity and extreme weather events can further impact the health and resilience of populations, rendering them more susceptible to mortality following HPAIV infection. This is of particular importance because many of the species identified here as highly susceptible to HPAIV share the characteristics of being relatively long-lived with low annual reproduction and high levels of parental care. Mortality of adults during the breeding season, as seen for the majority of seabirds, sea ducks, grebes, and raptors, would also indirectly impact reproductive success through increased nest failures from reduced hatching success or increased mortality of nestlings. The combination of increased mortality and decreased reproduction can result in significant population-level impacts, particularly for species or populations that are vulnerable, or are already experiencing multiple concurrent stressors (e.g., reduced food abundance or quality, increasing industrial or agricultural activity, urban encroachment, and other large-scale environmental changes associated with climate change). Concurrent stressors can also impact the ability of many of these populations to recover from mortality and reproductive failure associated with HPAIV.

### Genome Constellations

Reassortment is a recurring phenomenon among LPAIVs within wild waterfowl populations (64, 65), and H5 subtypes of clades 2.3.4.4 and 2.3.4.4b have demonstrated a high propensity to reassort with LPAIVs (44, 66). The co-circulation of HPAIV and LPAIV among wild bird reservoir species (e.g., dabbling ducks) increases opportunities for mixed infections and the emergence of reassortants (64). This is consistent with observations in the year following the first HPAIV incursion into Canada, where increased detection rates of new genome constellation patterns coincided with periods of increased LPAIV prevalence and concentrated wild bird abundance on the landscape during the spring and fall of 2022. In addition to these temporal patterns, there was evidence of geographic structuring of genome constellations at the flyway scale, whereas distinct geographic trends were not evident at the ecoprovincial scale within the Pacific flyway (Andrew et al. submitted). These temporal and spatial relationships underscore the dynamic interplay between virus prevalence, wild bird reservoir abundance, and movement (i.e., migration timing and pathways) in shaping reassortment dynamics. With the continued circulation and spread of H5N1 HPAIV and reassortants across Canada, homotypic and heterotypic immunity in wild bird reservoirs will also likely impact HPAIV dynamics (67), influencing the frequency and diversity of viral reassortants. Therefore, longitudinal surveillance targeted during periods of concentrated reservoir abundance, coordinated at the flyway scale, and incorporating serologic sampling, will collectively be needed if we wish to track and understand the complex, dynamic and rapid evolution of this virus.

The majority of reassortants detected in the first year since emergence resulted from the exchange of internal gene segments with North American LPAIVs, which is consistent with reports from the USA (68). The majority of sequences detected in Canada in the first year of surveillance post-incursion were categorized into four broad genome constellations including Eurasian H5N1 and Patterns 2, 4, and 5. The two most frequently detected genome constellations detected in the USA from December 2021 to April 2022, as similarly observed in Canada, were Eurasian H5N1 (genotype A1 in (68)) and Pattern 2 (genotype B2, B3.1, and B4), however Pattern 7 (B1.1 and B1.2) was also among the most common patterns found along with Pattern 4 (B3.2), and Pattern 5 was not detected in the USA in that time period (68).

Interestingly, the persistence of Eurasian virus was particularly notable in the Atlantic flyway throughout the first year following the first incursion. Although Eurasian viruses were sporadically detected in other flyways, their presence was transient, and they were quickly outnumbered by reassortant viruses. It is not clear what ecological, evolutionary, or viral factors were driving this persistence in the Atlantic flyway. Potential drivers could include variation in the prevalence and composition of LPAIVs subtypes circulating within the Atlantic region compared to other flyways, differences in survival following infection with subsequent impact on opportunities for virus reassortment, or species-specific interactions unique to the Atlantic flyway. However, sampling biases in the composition of species, locations, and timing, in addition to diagnostic considerations (e.g., samples yielding lower PCR cycle threshold values were more likely to result in higher quality sequence data and subsequent inclusion in analyses), mean that observed patterns reflect available sequences and are therefore unlikely to represent the complete diversity and distribution of viruses present in wild bird populations.

### Surveillance Components and Sampling Limitations

It is important to note that while live and hunter-harvested (i.e., ‘active’) and sick and dead (i.e., ‘scanning’ or ‘passive’) wild bird surveillance methods can be complementary and contribute data from different subsets of wild birds (i.e., those that do and do not survive infection), each surveillance method has limitations and biases that are critical to understand to contextualize the results presented here (69).

The majority of sick and dead wild bird carcass submissions are opportunistically submitted by members of the public and therefore originate from more populous areas of Canada (Fig. 2). Geographic proximity to diagnostic centers and higher human population densities increase the likelihood of carcass detection and submission (70). Biases in species detectability (e.g., size, habitat with dense vegetation vs. open parkland) and the likelihood of submission based on social (e.g., perceived as a nuisance vs. highly valued) or other factors (e.g., disparate levels of awareness between communities) can also influence which samples are processed through this surveillance component (71). Therefore, absence of detection through sick and dead bird surveillance does not imply the absence of infection and mortality.

Live and hunter-harvested wild bird surveillance is also opportunistic in that it is carried out in conjunction with existing banding and monitoring programs. By strategically targeting wild birds during periods and in areas of high abundance, banding and monitoring programs are well-aligned with locations and time periods expected to have increased AIV prevalence. However, these programs are often conducted over short time frames (days or weeks), only during certain months, and limited to focal areas. Consequently, this can limit our ability to detect infection, which for AIV consists of a relatively short viral shedding period (72), and to track changes in incidence and prevalence within these high-risk areas and time periods. Despite these limitations, the continued integration of both sick and dead as well as live and harvested wild bird surveillance remains crucial to understand HPAIV dynamics in wild birds.

## Supporting information

Canada Interagency Avian Influenza Implementation Plan

## Acknowledgements

We extend our sincere appreciation to members of the public who reported sick and dead birds, rehabilitation centers and migratory game bird hunters who reported observations and contributed samples, and conservation officers and other staff who responded to calls throughout the course of the outbreak. We would also like to thank the Nunatsiavut Government, NunatuKavut Community Council, Miawpukek First Nation as well as the Nunavik Research Center of Makivvik and the Cree Board of Health and Social Services for their role in either the collection of carcasses or harvested bird samples submitted by local harvesters and conservation officers. These valuable inputs have greatly enriched the depth and quality of our work. This manuscript was prepared on behalf of all partners that contribute to Canada’s Interagency Surveillance Program for Avian Influenza Viruses in Wild Birds, including Canadian Wildlife Health Cooperative, and Federal, Provincial, Territorial, Indigenous, and Academic partners involved in wildlife, domestic animal, and human health. See Table S2 for acknowledgement of additional collaborators that have contributed to the collection and curation of these data.

## Additional Supplementary Material

Video 1. Time series of sick and dead wild birds confirmed to be highly pathogenic avian influenza virus (HPAIV) positive in Canada between November 2021 and December 2022. Taxonomic grouping represented by coloured symbology.

Supplemental Document. Canada’s Interagency Surveillance Program for Avian Influenza Viruses in Wild Birds: 2022-2023 Implementation Plan.

## Supplemental Material

**Fig. S1.**
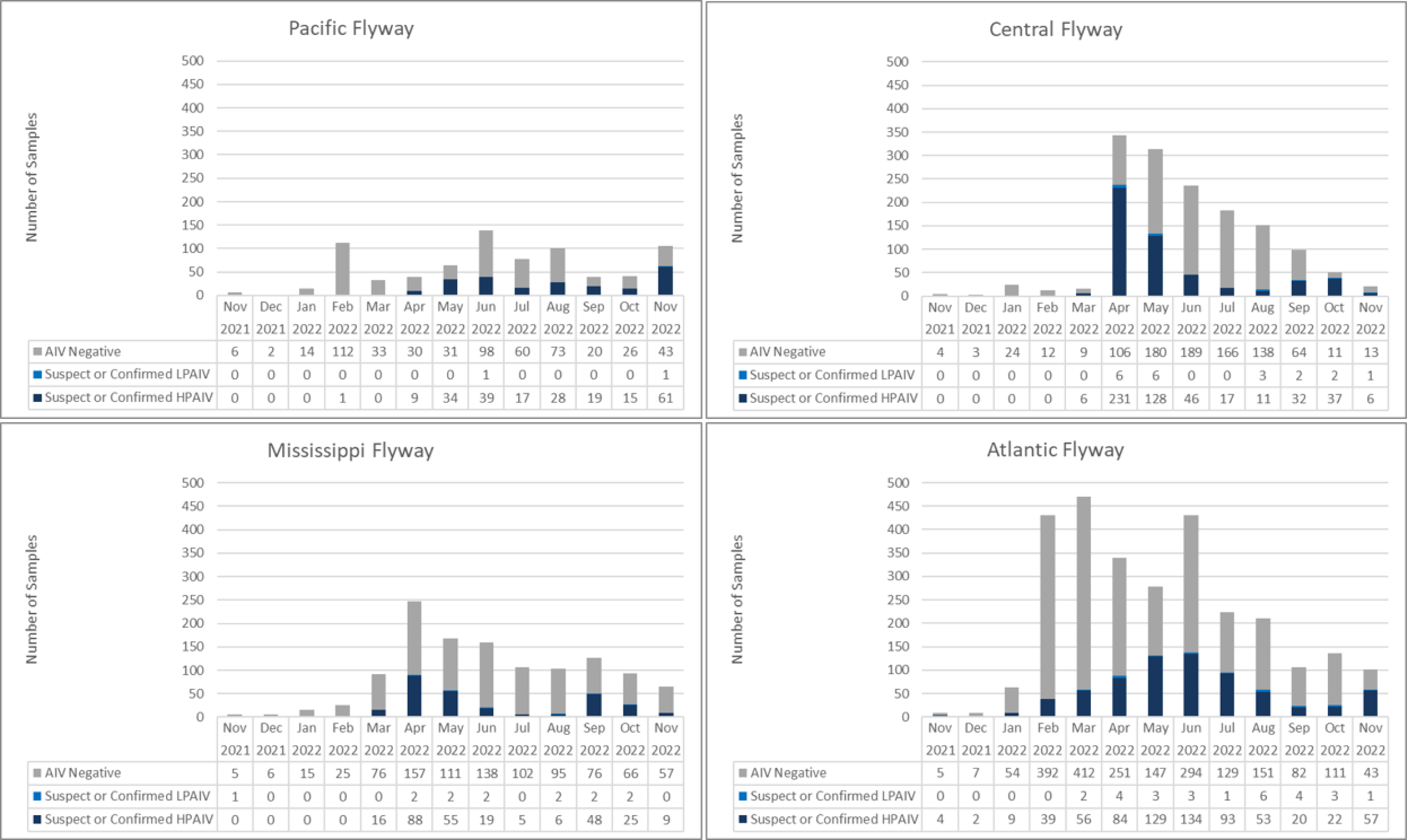
Sick and dead wild birds tested for avian influenza virus between November 2021 and November 2022 across Canada stratified by flyway.

**Fig. S2.**
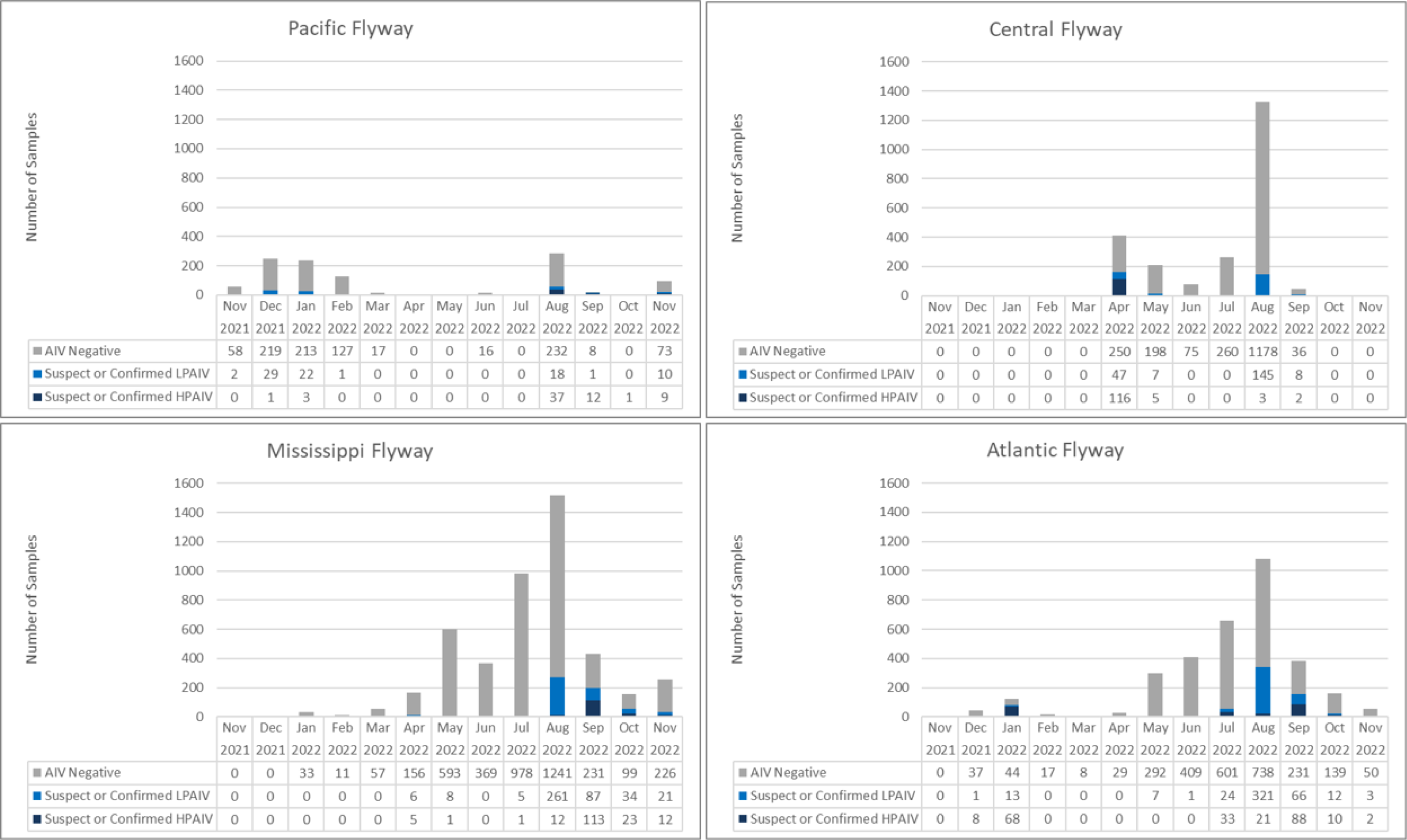
Live and hunter-harvested wild birds tested for avian influenza virus between November 2021 and November 2022 across Canada stratified by flyway.

**Fig. S3.**
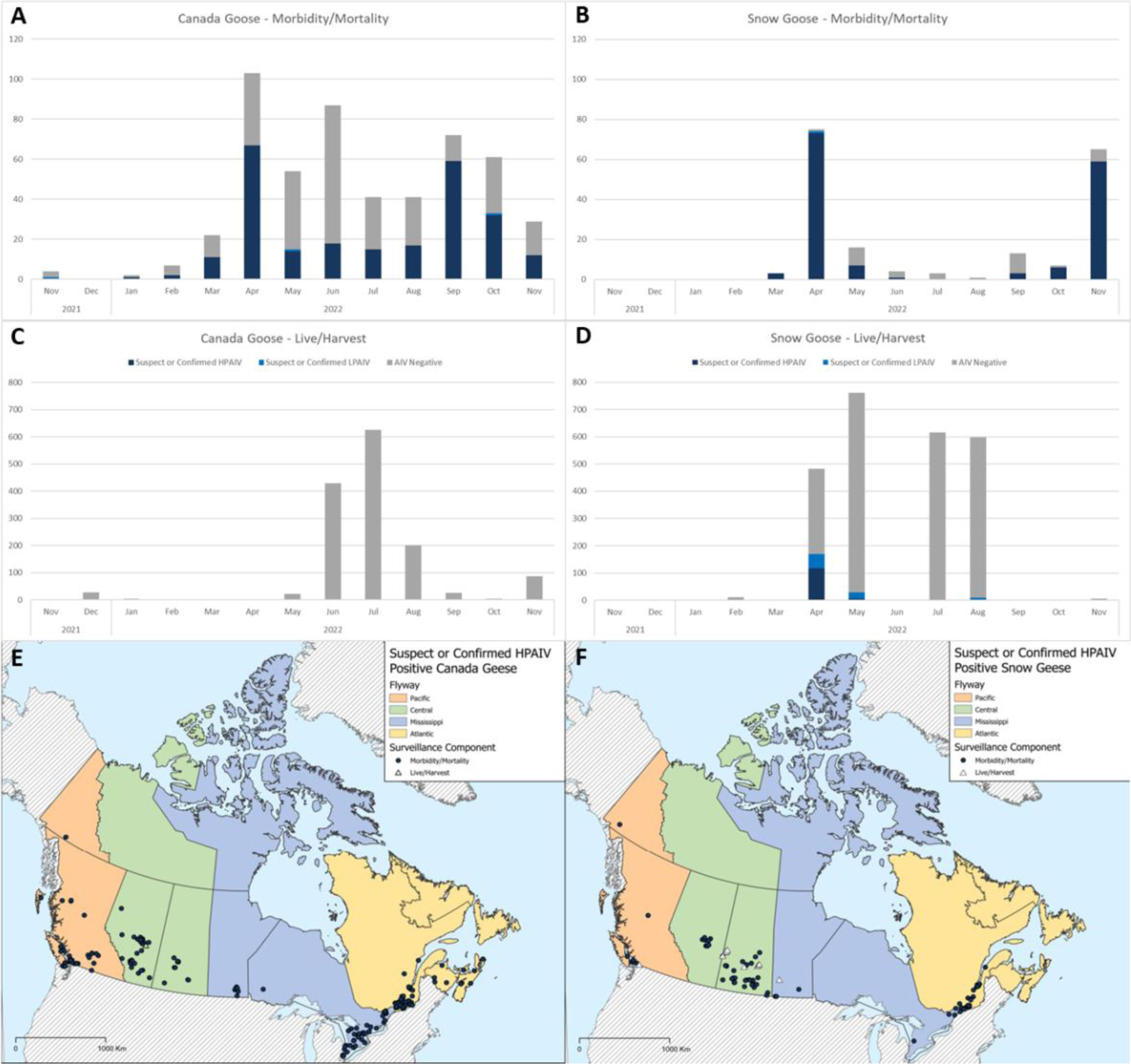
Canada Geese (A, C) and Snow Geese (B, D) tested for avian influenza virus between November 2021 and November 2022 across Canada stratified by surveillance component (i.e., sick/dead and live/hunter harvest). The location of suspect and confirmed highly pathogenic avian influenza positive Canada Geese (E) and Snow Geese (F).

**Fig. S4.**
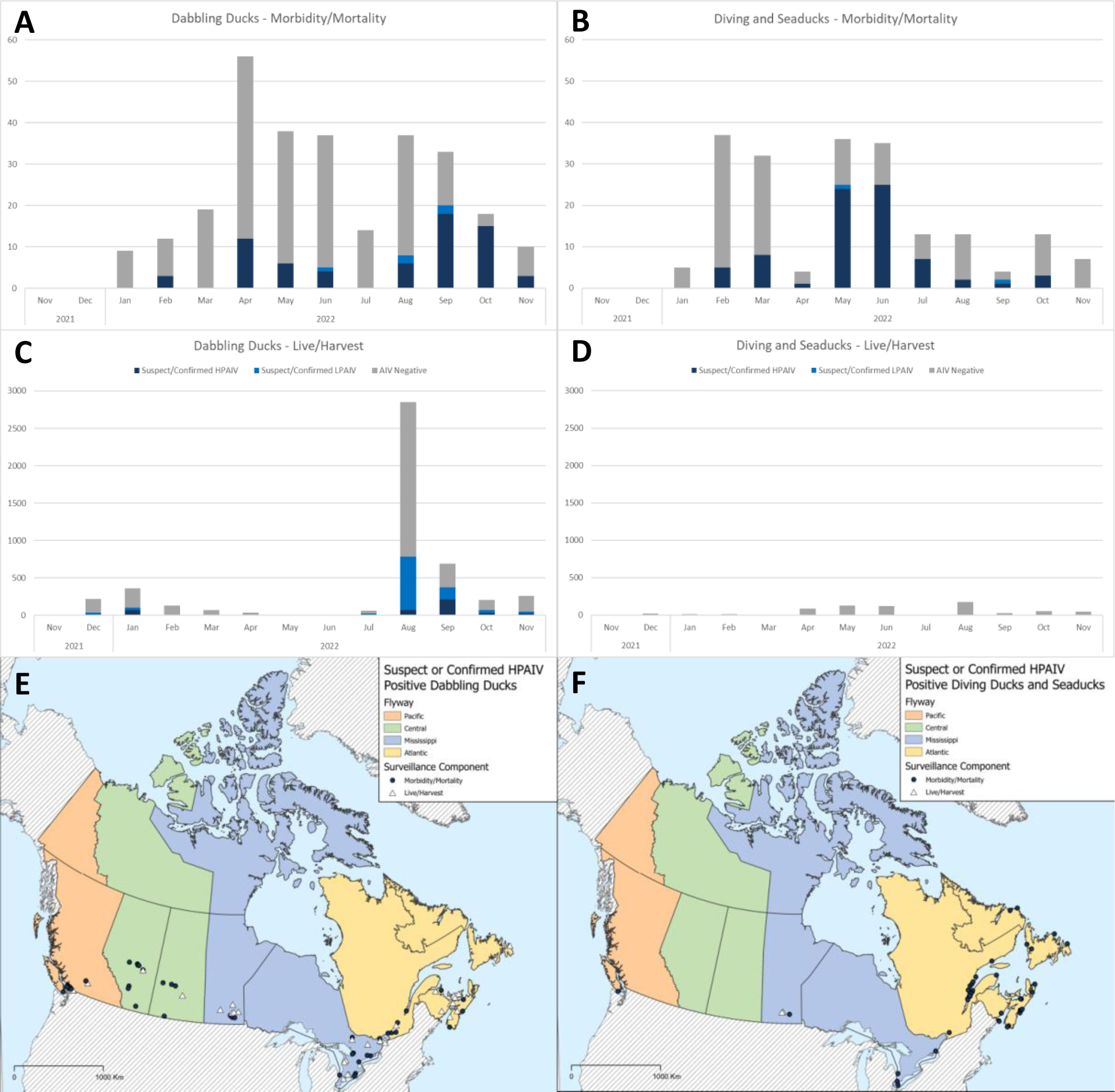
Dabbling ducks (A, C) and diving ducks and seaducks (B, D) tested for avian influenza virus between November 2021 and November 2022 across Canada stratified by surveillance component (i.e., sick/dead and live/hunter harvest). The location of suspect and confirmed highly pathogenic avian influenza positive dabbling ducks (E) and diving ducks and seaducks (F).

**Fig. S5.**
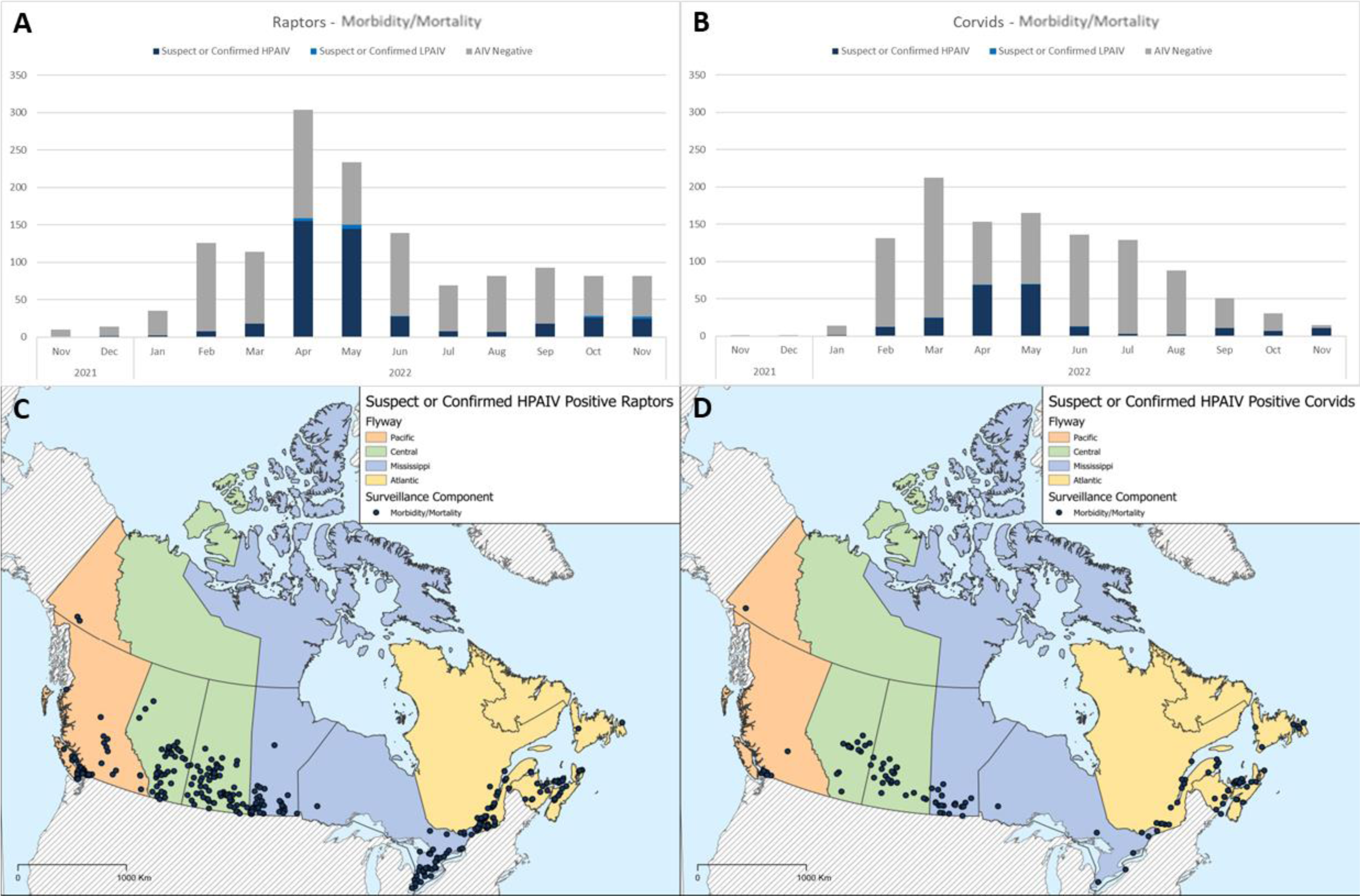
Raptors (A) and corvids (B) tested for avian influenza virus between November 2021 and November 2022 across Canada through morbidity/mortality surveillance. The location of suspect and confirmed highly pathogenic avian influenza positive raptors (C) and corvids (D).

**Table S1.**
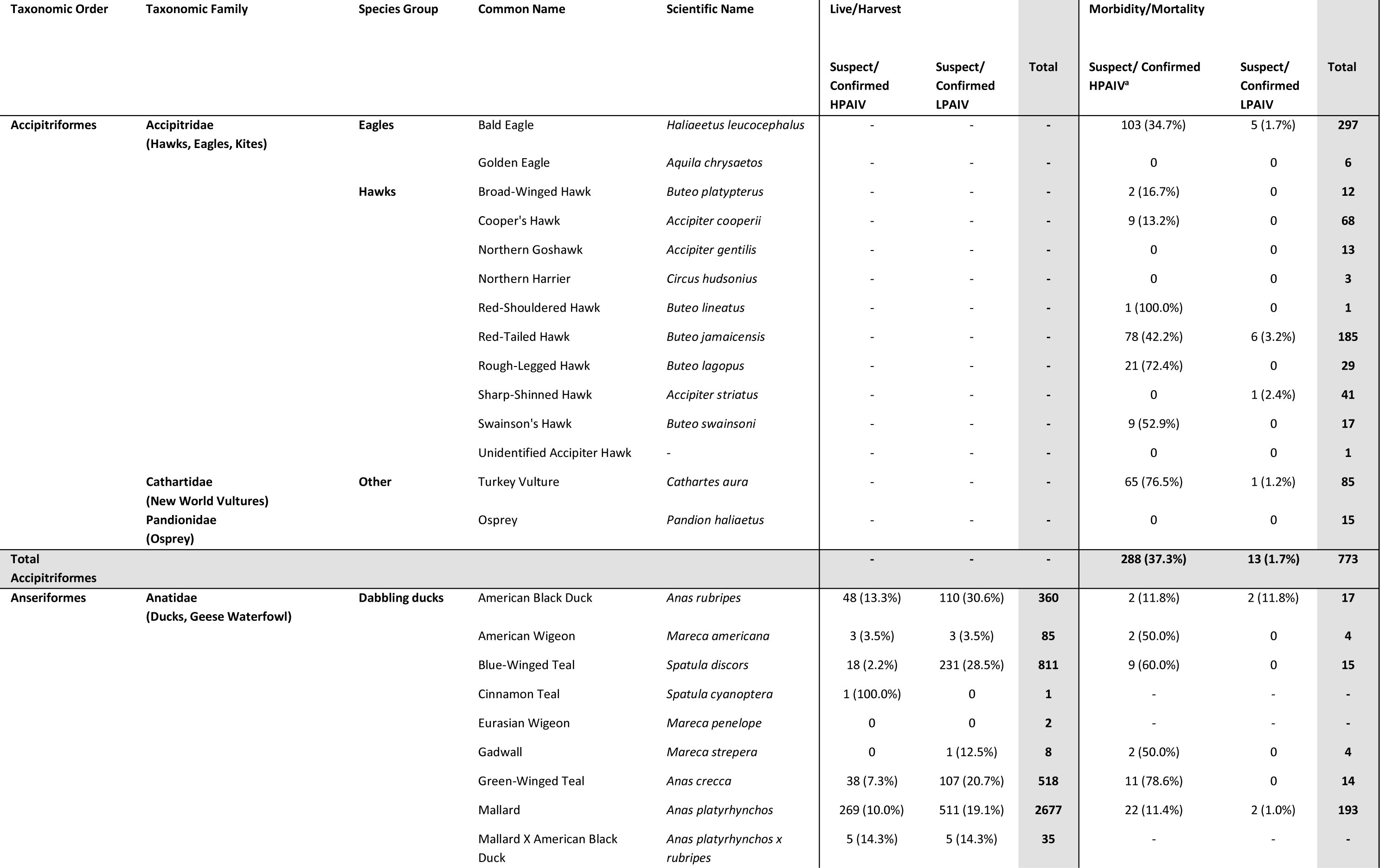
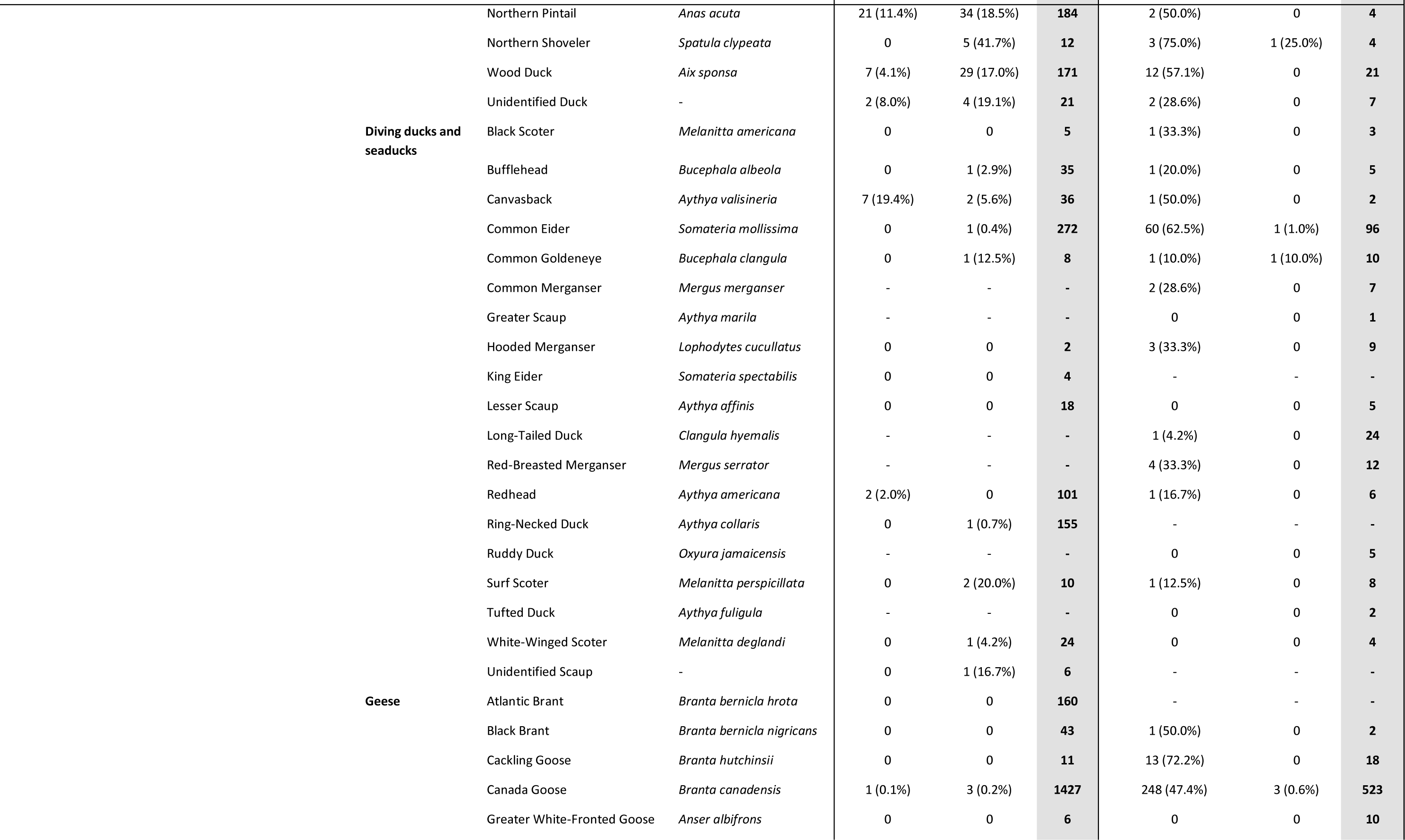
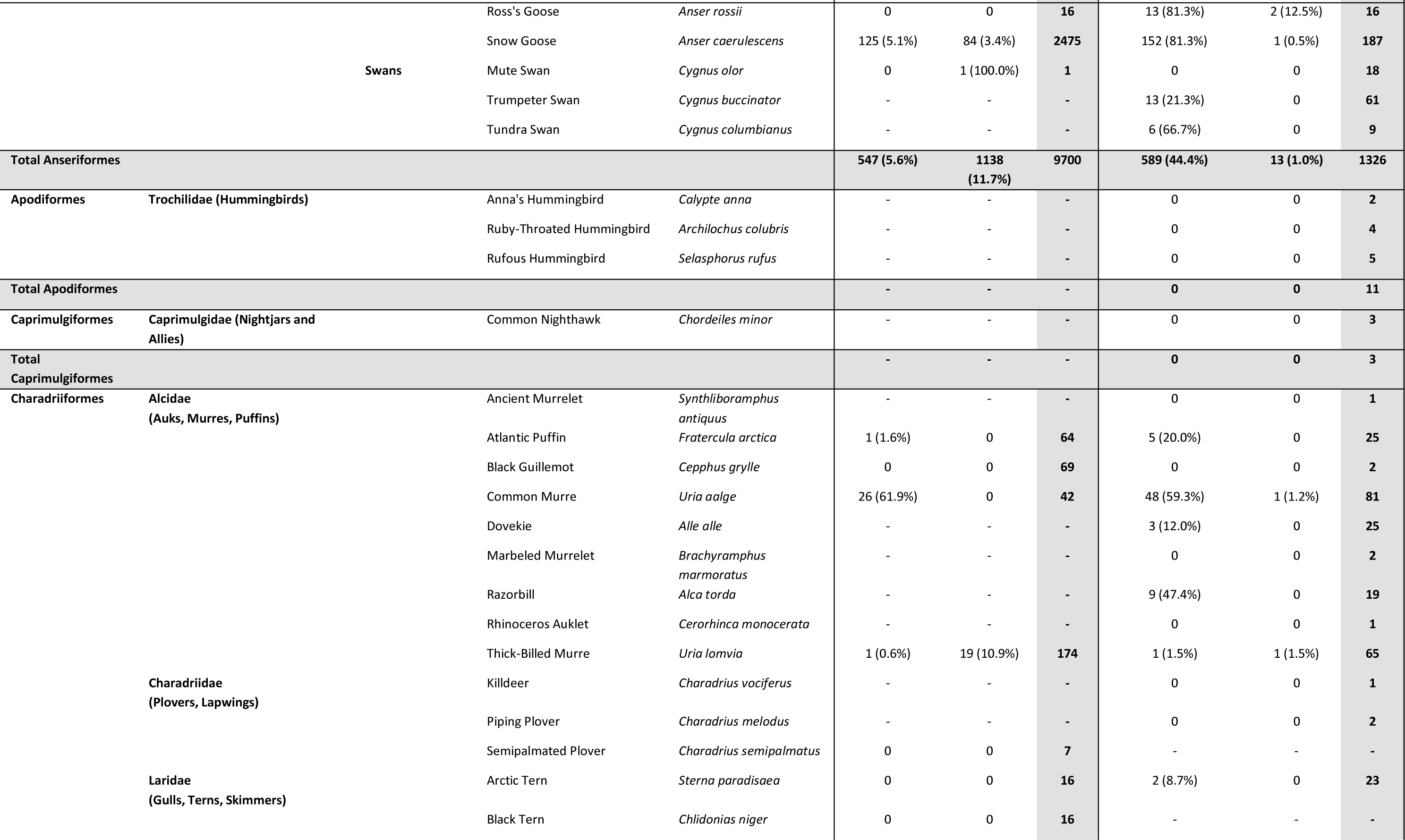
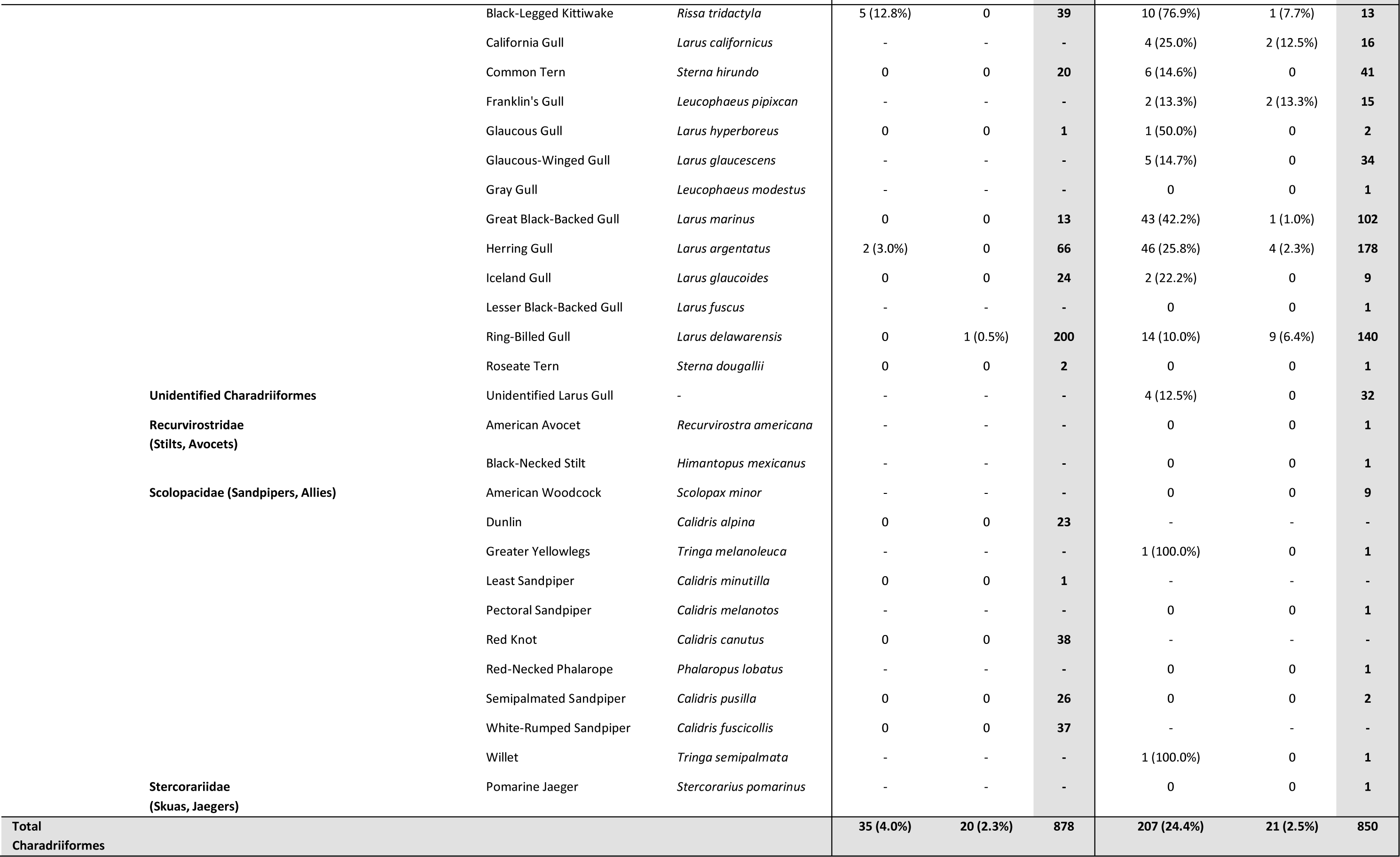
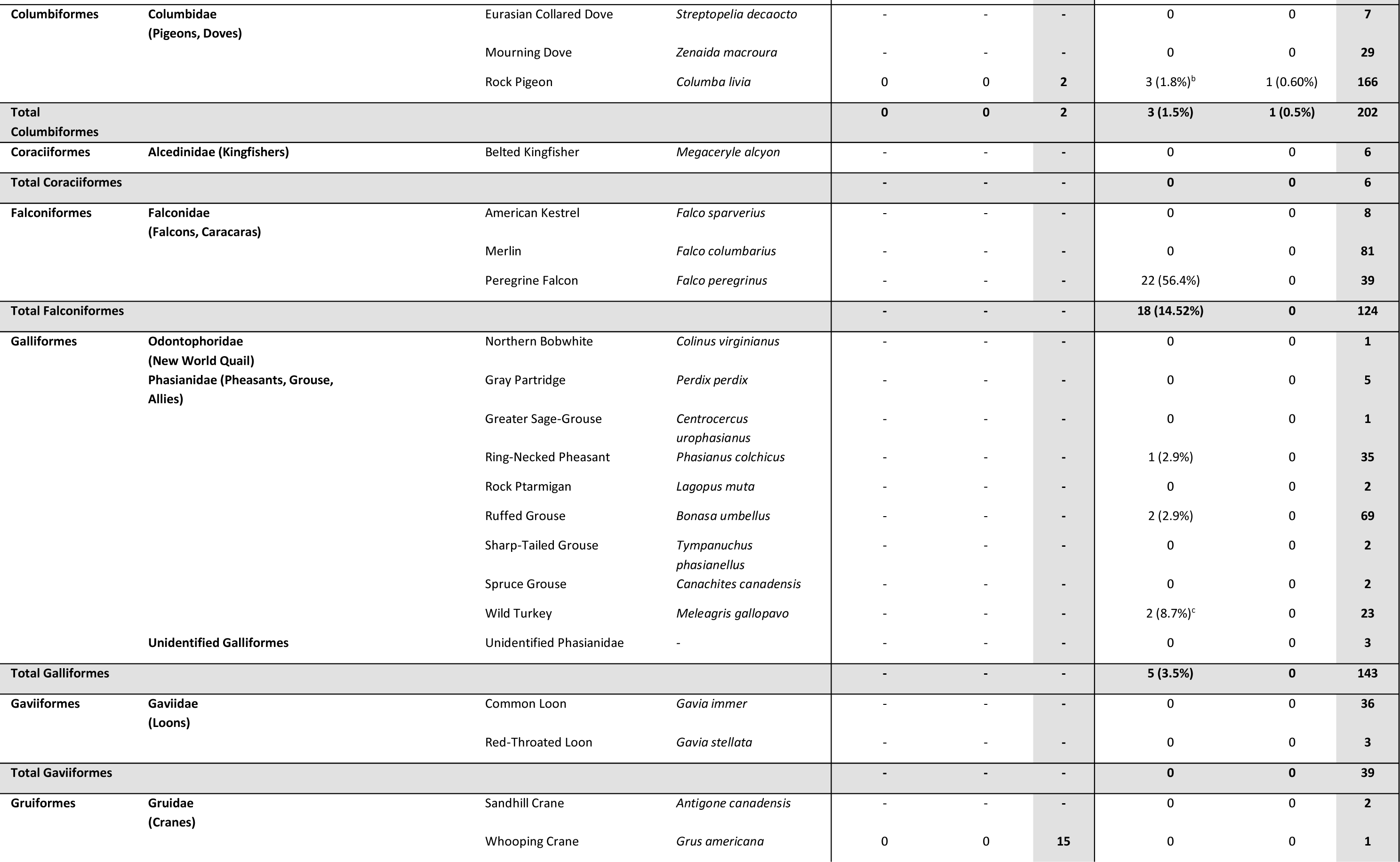
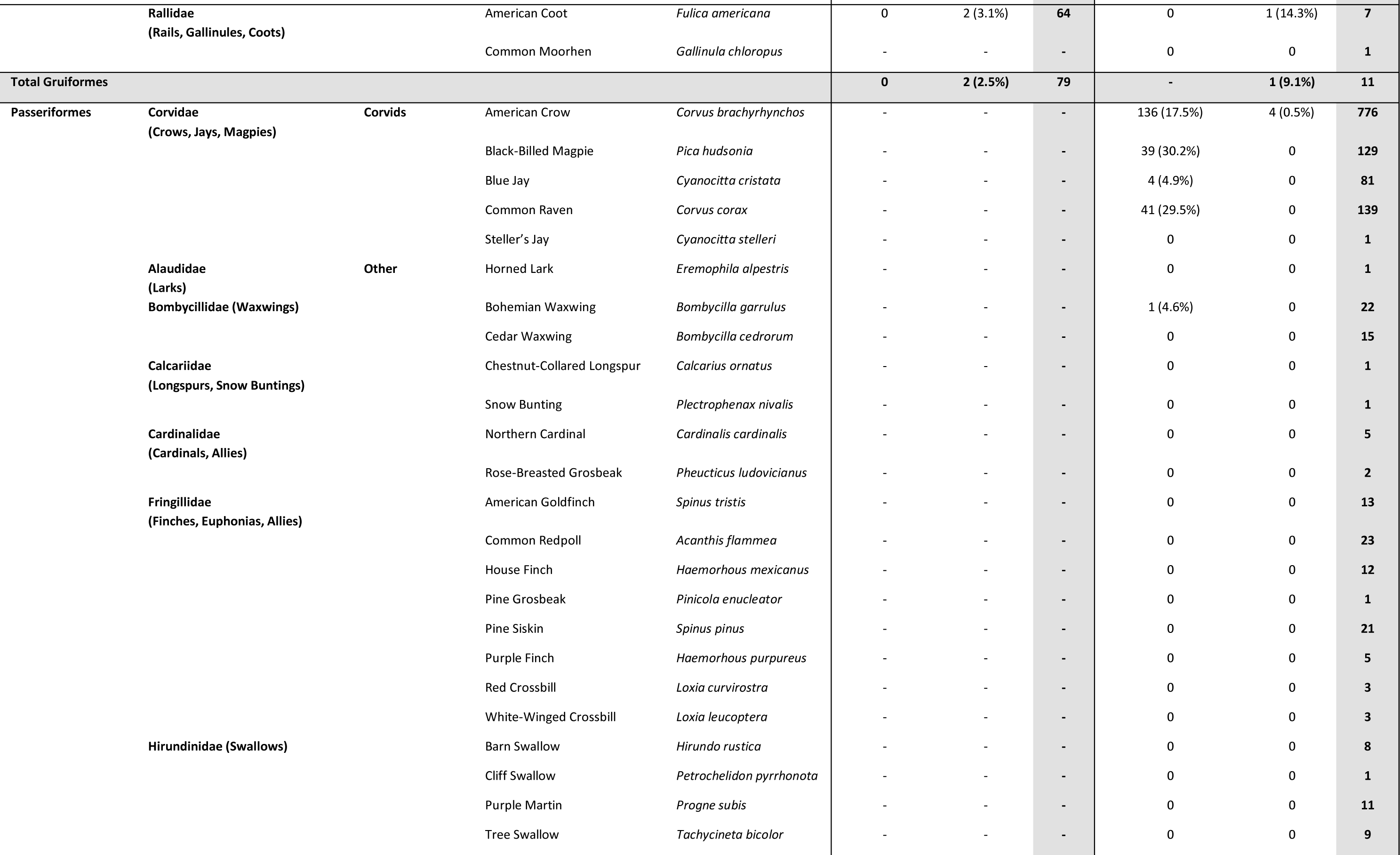
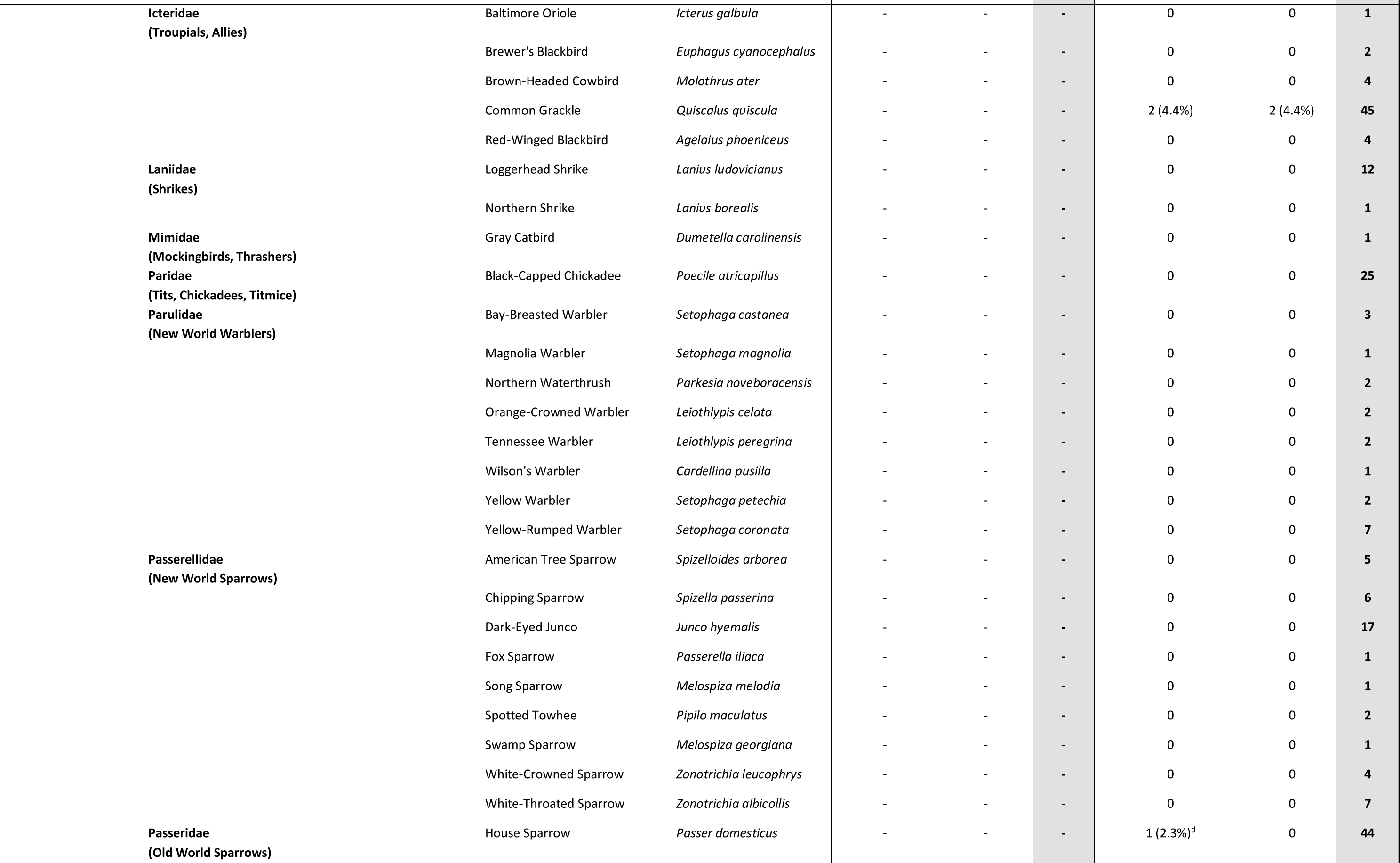
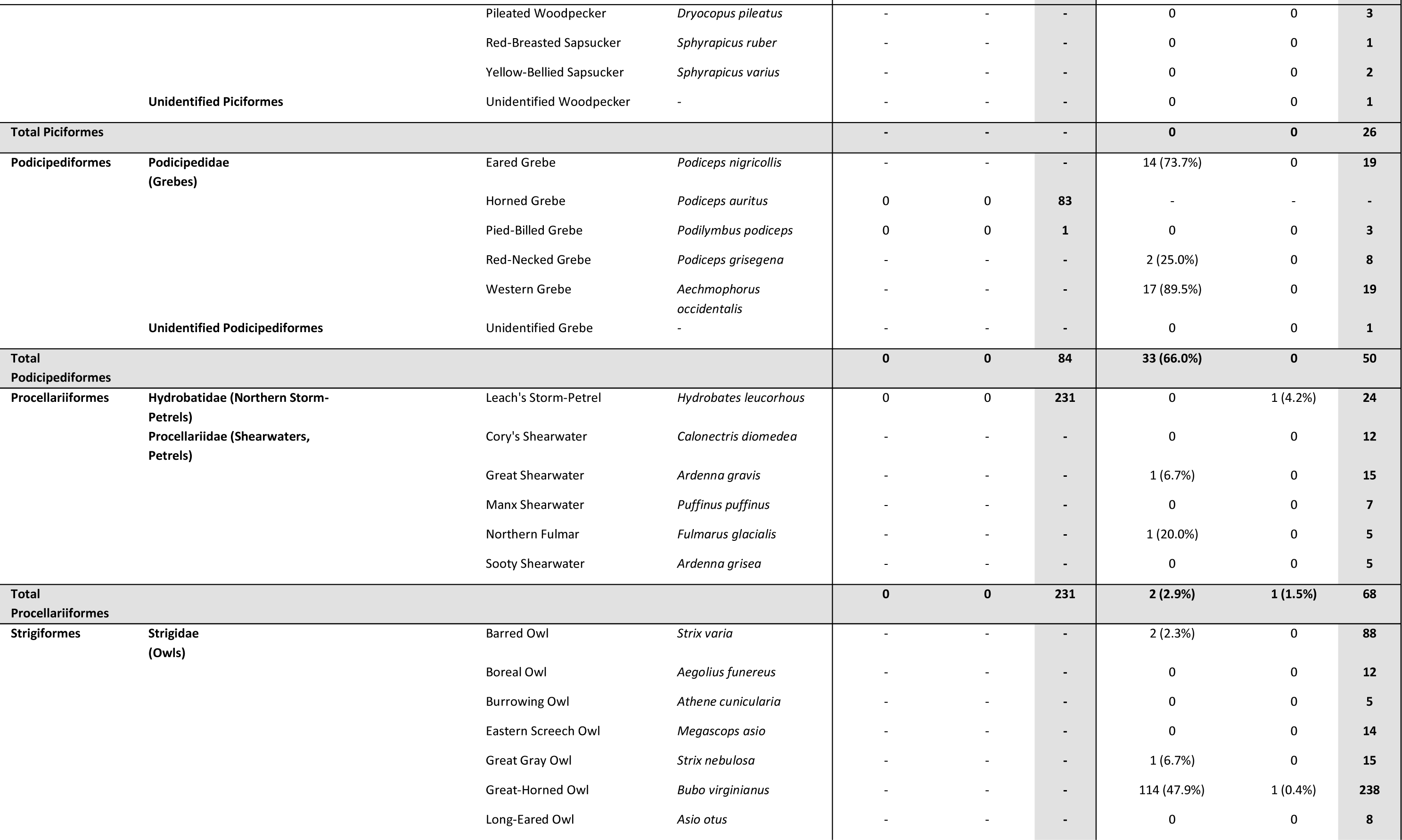
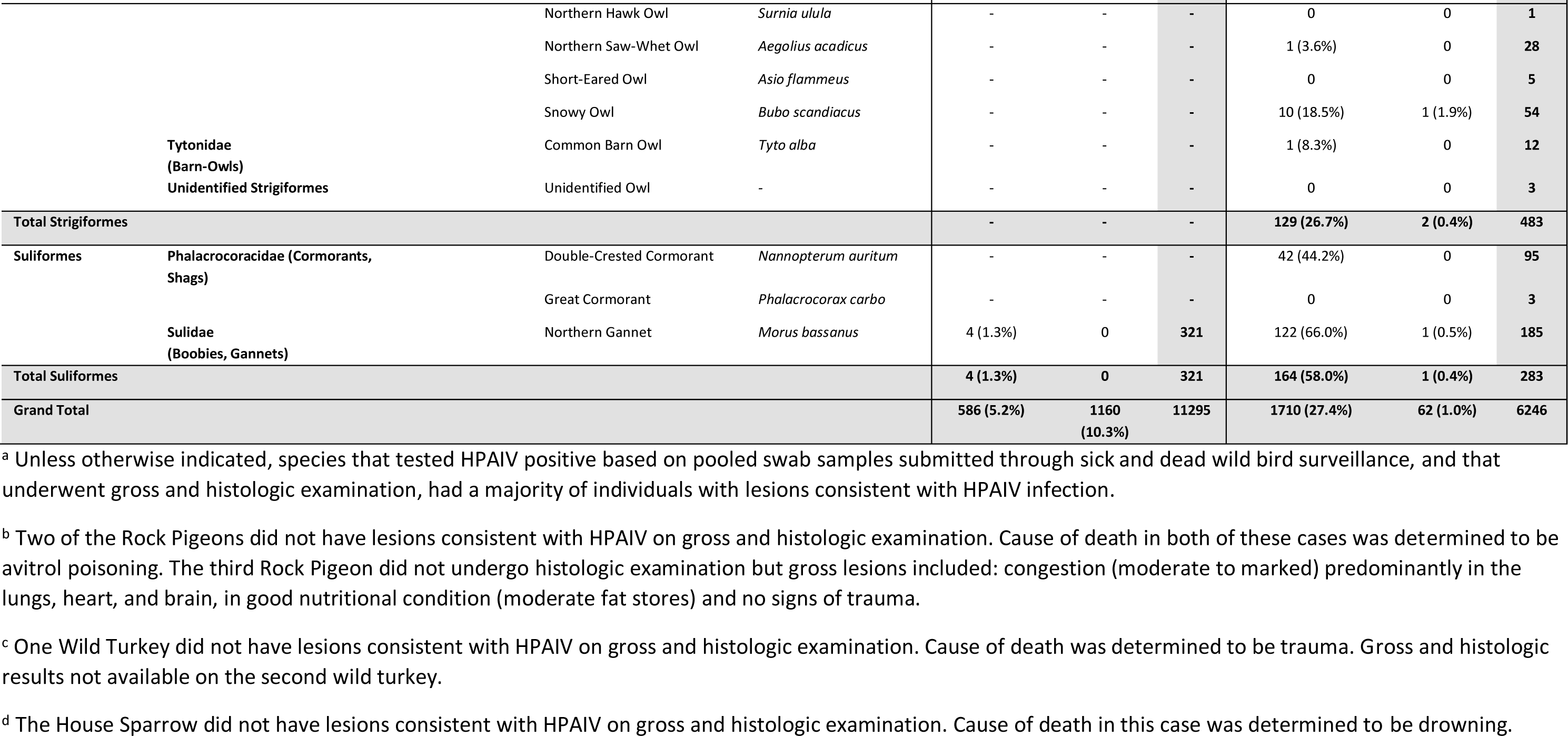
Number of live and hunter-harvested and sick and dead wild birds submitted for testing and suspect or confirmed highly pathogenic avian influenza virus (HPAIV) or low pathogenicity avian influenza virus (LPAIV) positive in Canada between November 2021 – November 2022.

**Table S2.**
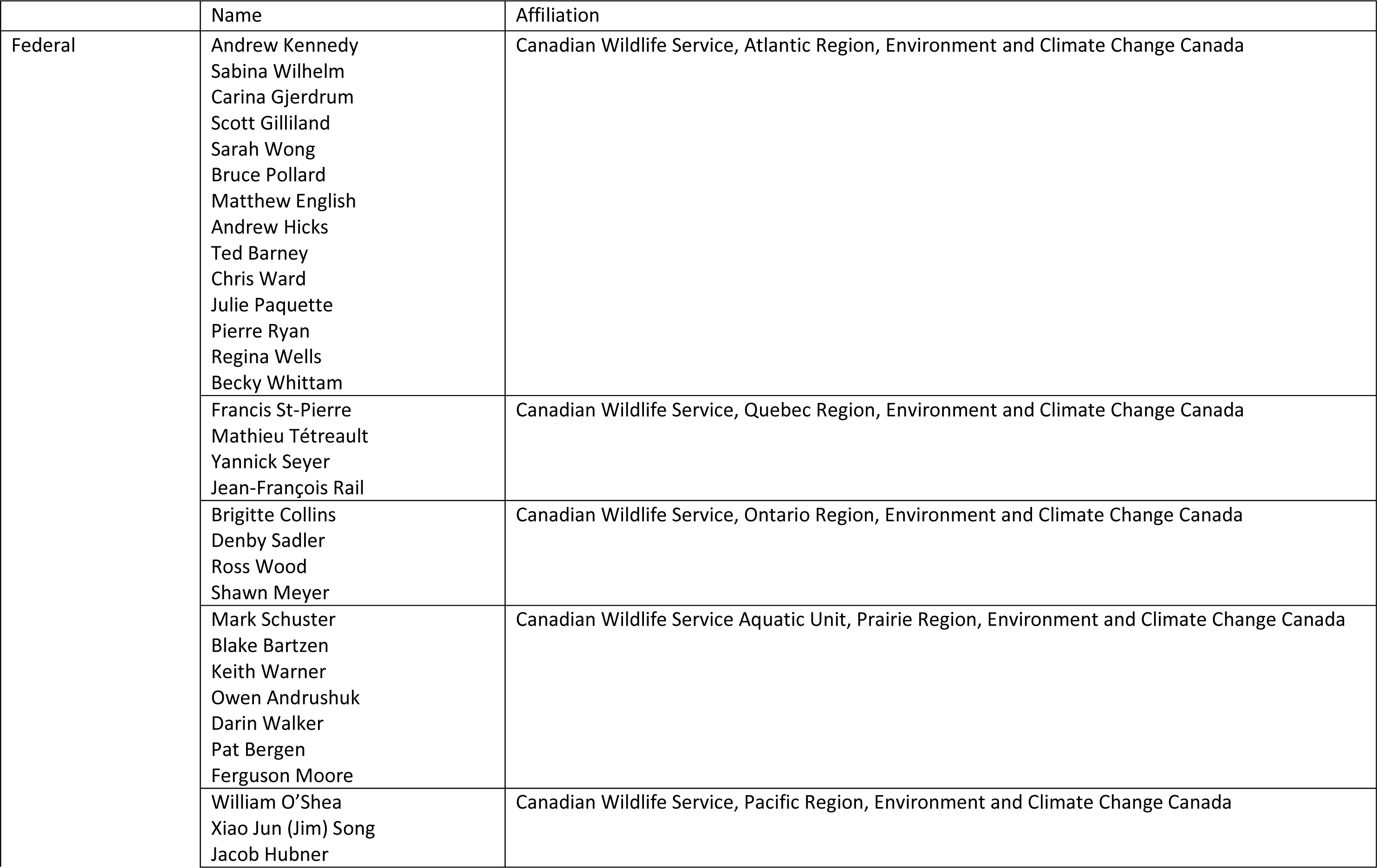
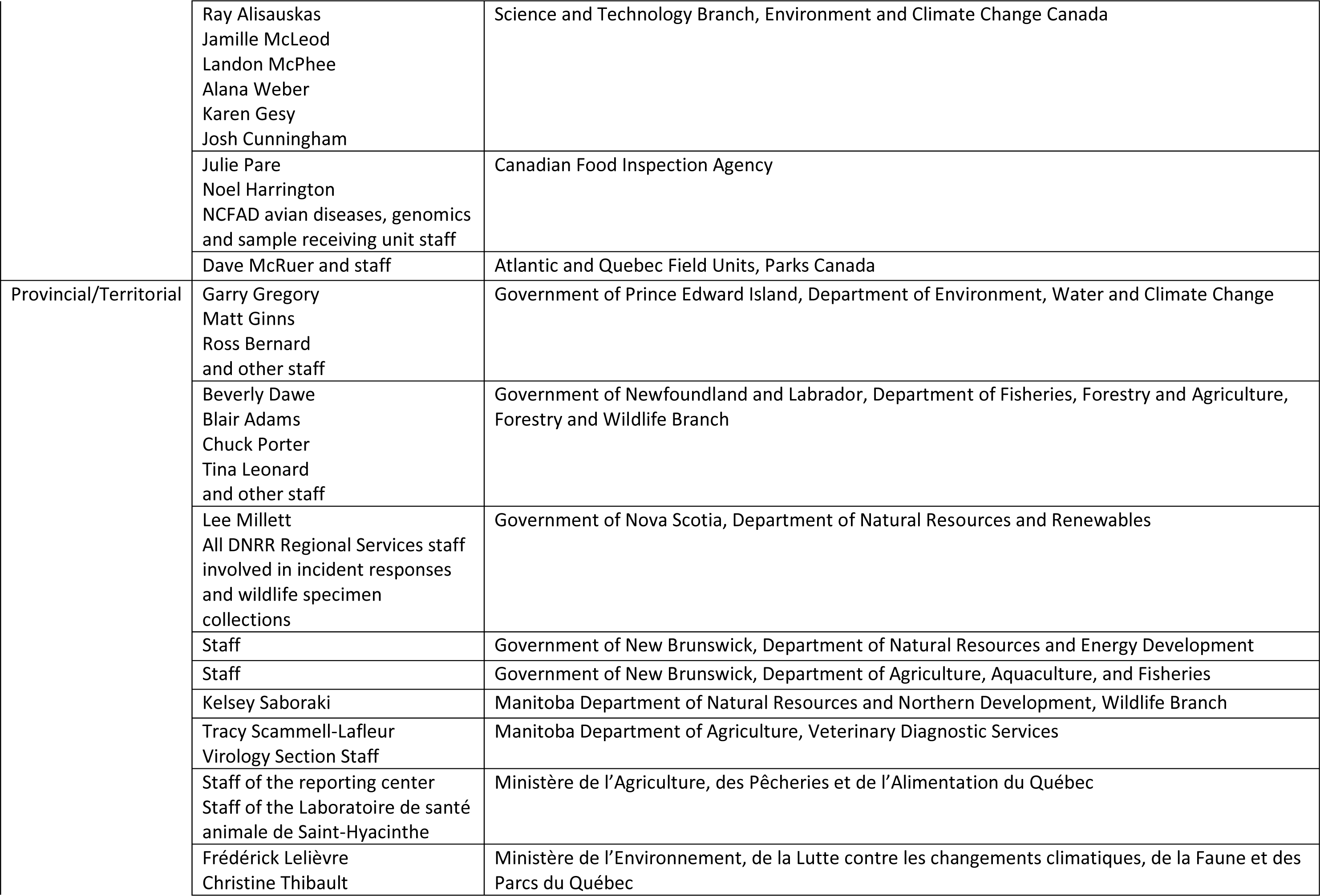
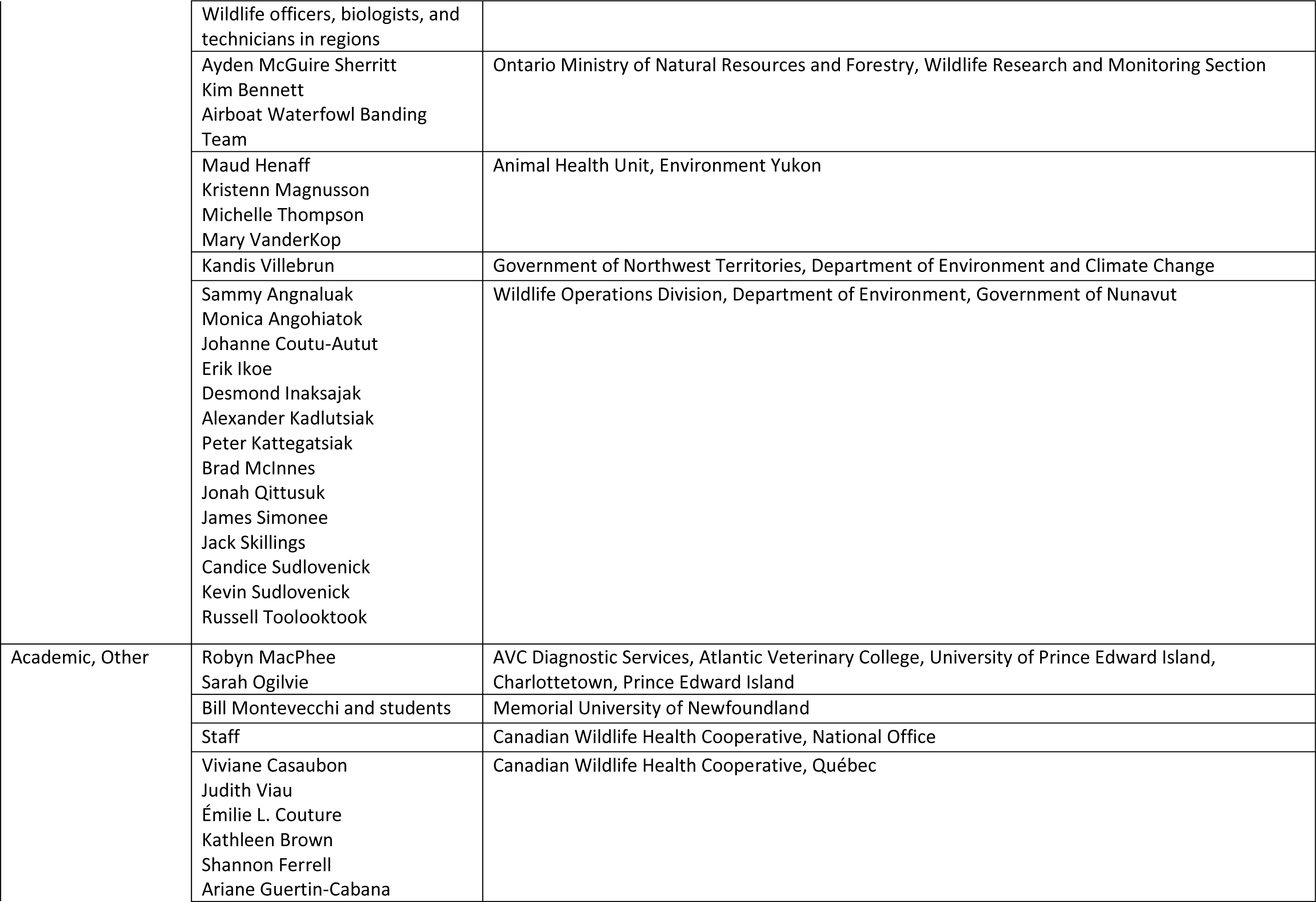
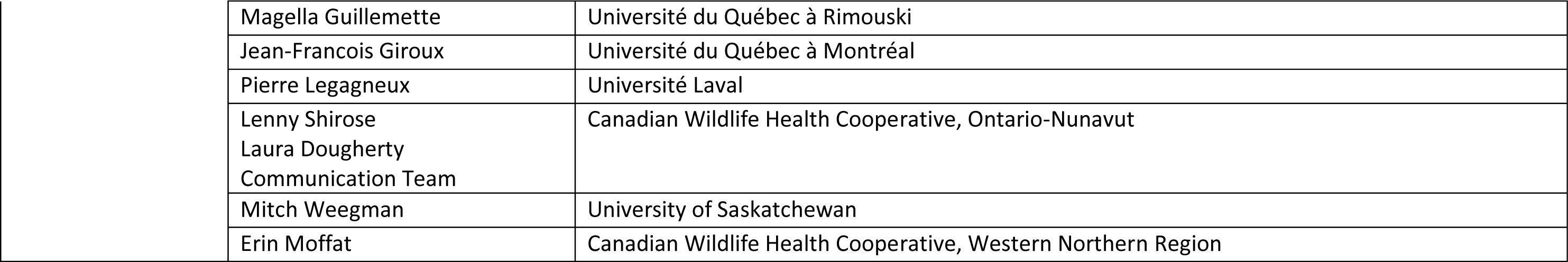
Acknowledgements of collaborators that have contributed to the collection and curation of these data but are not listed as co-authors.

