## Supplementary material for "Avian influenza viruses in wild birds in Canada following incursions of highly pathogenic H5N1 virus from Eurasia in 2021/2022": Canada Interagency Avian Influenza Implementation Plan

### Canada's Interagency Surveillance Program for Avian Influenza Viruses in Wild Birds: 2022-2023 Implementation Plan

This document has been prepared by a joint effort of ECCC-STB and ECCC-CWS. This document has been reviewed by the Wild Bird AI Surveillance Working Group which consists of partners listed in Appendix A of this document.

Date: August 8, 2022

### 1. Intent

The intent of this document is to serve as guidance for the 2022-2023 implementation of Canada's Interagency Surveillance Program for Avian Influenza Viruses in Wild Birds. The development of a stand-alone strategic plan document is still an active area of discussion, therefore, key strategic priorities have been incorporated in this current version of the guidance document where deemed appropriate.

This document is one of several developed by Environment and Climate Change Canada (ECCC) in 2022 in response to the emergence of H5N1 Highly Pathogenic Avian Influenza Virus (HPAIV) in Canada in December 2021. A list of additional ECCC Protocols, Policies, and Guidance documents can be found in Appendix I.

### 2. Status of Avian Influenza in Wild Migratory Birds and Other Wildlife

A state of the science document has been prepared by ECCC, and can be reviewed for detailed background information including information on HPAIV in the global and North American context, potential impacts and implications, and management options.

The dynamics (emergence, spread) and impacts of HPAIV in wild birds is a rapidly evolving situation. Up to date information can be found online through Canada's Avian Influenza dashboard, the USGS Wildlife Health Information Sharing Partnership (WHISpers) portal, and the USGS map of HPAIV in North America. A list of additional relevant links can be found in Appendix B.

In summary, H5N1 HPAIV has been detected in all four North American flyways in Canada and the USA (Atlantic, Mississippi, Central, Pacific), and spread rapidly within and between flyways during the early spring migration months. Between December 29, 2021, and May 12, 2022, Canada confirmed HPAIV (H5N1 or H5 reassortments) in wild birds in all ten provinces. To date, confirmed or suspected results have been observed in numerous species comprising waterfowl, seabirds (e.g., gannets, murres), raptors, gulls, corvids, grouse, pheasants, pigeons, and others. Early in the outbreak multiple mortality events caused by HPAIV have been detected across Canada, including in snow geese, ranging from individuals to groups of 100 in the Prairie provinces, and in green-winged teal (approximately 15) and red-breasted mergansers (approximately 60) in southern Ontario. As of May 27$^{th}$, multiple and large scale morbidity/mortality events were reported in northern gannets in the Atlantic region and Quebec. Large mortality events associated with HPAIV have also been reported in the United States (e.g., > 1000 lesser scaup in Florida).

As of July 22$^{nd}$ 2022, confirmed or suspected HPAIV (H5N1) results have also been reported in mesocarnivore species (including red fox kits, striped skunk, American mink, and harbour seals) in at least 6 provinces in Canada and in a number of states in the United States.

### 3. Background

Canada's Interagency Surveillance Program for Avian Influenza Viruses in Wild Birds (formerly referred to as Canada's Interagency Wild Bird Influenza Survey) has been undertaken by the Government of Canada and Canada's provinces and territories since 2005. Historically, the surveillance program had been coordinated on behalf of government agencies by the Canadian Cooperative Wildlife Health Centre

(CCWHC), now the Canadian Wildlife Health Cooperative (CWHC), and formed part of national and global efforts to detect avian influenza viruses (AIVs) of significance to wildlife, domestic animals, and human health. When funding and interest in surveillance of live wild birds for AIV decreased (in 2012, and again in 2017, Appendix C), surveillance efforts across Canada dwindled, with the exception of ongoing research/surveillance programs that continued in the Prairie Provinces and other regions (e.g., Atlantic provinces, British Columbia). In 2015, funding and interest in AIV surveillance in live wild birds briefly increased associated with detections of HPAI in wild birds in British Columbia and in poultry facilities in British Columbia and Ontario. The CWHC dead bird surveillance component has continued to operate throughout this period.

In early 2021, given the significantly increased reports of HPAIV H5N1 throughout Eurasia, ECCC (both Canadian Wildlife Service (CWS) and Science & Technology Branch (STB)) and the Canadian Food Inspection Agency (CFIA) began discussions to coordinate and enhance surveillance for AIV in live birds across Canada in the summer and fall of 2021, in anticipation of an incursion of the virus into Canada. Funding, coordination, and support for 2021-22 surveillance was provided by ECCC-CWS and significant in-kind support from ECCC-STB and NCFAD-CFIA. ECCC was able to employ and build on the previously established network, with the support of interagency partners, to collect and analyze more than 2300 samples across Canada in the summer and fall of 2021. Efforts for live bird surveillance in spring 2022 are currently being coordinated through a joint effort between ECCC-CWS and STB, and will form part of 2022-23 surveillance efforts.

#### 4. A One Health / Collaborative Approach is Necessary in 2022-2023

The term "One Health" recognizes connections between people, animals, plants, and the environment. In a One Health approach, multiple sectors communicate and collaborate to address shared health threats (World Health Organization (WHO) 2021). The two core objectives of the OIE Wildlife Health Framework are to manage the risk of disease emergence at the human-animal-ecosystems interface, and to protect wildlife health. The Convention on Migratory Species (CMS) and the United Nations Food and Agriculture Organization (FAO) co-convened a Scientific Task Force on Avian Influenza and Wild Birds, and published a statement on H5N1 Highly Pathogenic Avian Influenza in poultry and wild birds on January 24, 2022. The statement emphasizes several important points:

- that wild birds, including those that are globally threatened, have been significantly impacted by this virus, and are not merely vectors of transmission;
- that intensified surveillance and biosecurity measures are critical to reduce risks of transmission between wild birds and poultry (in both directions);
- that any mitigation responses must not cause harm to wildlife or the environment, but focus on minimizing risks to wildlife and sites from poultry and humans;
- that One Health approaches are essential for all aspects of monitoring, responding to, or mitigating this virus, including science-based communications, coordination, surveillance, and research on the epidemiology and impacts of this virus in wild bird populations.

Given that HPAIVs have potential impacts on wildlife, domestic animal, and/or human health, it is essential for Canada to take a One Health approach to investigating, detecting, and managing outbreaks. Plans for surveillance, investigation, management, and communications must involve multiple jurisdictions. This current surveillance plan document is part of a broader series of HPAIV surveillance documents (Appendix I); input into the current plan has involved inter-agency dialogue/engagement.

The potential for the emerging strains of HPAIV to become established in migratory bird populations, spread to multiple species and populations, evolve, mutate, and combine with other AIVs to generate novel viruses, which can spill back into domestic animal populations or to humans is currently unknown. Therefore, timely detection and genomic analysis of positive samples, particularly those in proximity to commercial or backyard flocks, would benefit from federal/provincial/territorial wildlife, agricultural, and public health authorities tasked with demonstrating that all avenues for viral spread are being investigated. In Canada, multiple jurisdictions are responsible for the management of response(s) to wildlife diseases at the human-animal interface, and thus rapid communications, data sharing, and collaboration on surveillance, research, and management activities will be beneficial to wildlife, public health, agriculture, conservation agencies, and Indigenous communities. Exclusion of specific agencies could hamper their ability to exercise due diligence and meet regulatory responsibilities. Interagency One Health HPAIV meetings have taken place since January 2022 to help achieve the outcomes highlighted in this document.

### 5. Rationale

Highly pathogenic avian influenza virus (HPAIV) H5N1 subtype clade 2.3.4.4b is a newly emerging virus in Canada and North America, with significant potential impacts on wildlife and domestic animal health. Since December 2021, the virus has affected an unprecedented number of wild birds and mesocarnivores and has impacted commercial, small flock, and other captive poultry facilities. The virus has also made at least two separate incursions into Canada since March 2022 and appears to be spreading rapidly within and between flyways. There is considerable overlap and exchange between the four main migratory bird flyways in North America, and the potential for further spread across Canada (and across North America) during spring and fall migration is significant. Based on data generated from Europe, Asia, Africa, and North America, this current H5N1 HPAIV appears to be very different from previous HPAIVs in how it behaves in wild birds including:

- Larger susceptible host range compared to previous HPAIVs (more species affected)
- Significant geographic range expansion of the virus with the largest extent of intercontinental spread detected compared to any previous HPAIVs
- Highest levels of wild bird (and mesocarnivore) mortality compared to previous HPAIVs in Europe, Asia, and North America
- Higher prevalence in asymptomatic wild bird populations than previously observed for other HPAIVs, and increasing evidence for maintenance, and rapid evolution of the virus in asymptomatic wild bird populations (primarily dabbling ducks), suggesting the potential for this virus to become endemic in some wild bird species that could serve as reservoirs and sources of exposure to susceptible wild birds and other wildlife species and poultry

The current extent of spread of this virus in Canada is unknown, but is expected to spread rapidly based on experiences in the eastern hemisphere. Hence, increased vigilance, as well as sustained and enhanced surveillance for HPAIVs across Canada are essential to understanding, evaluating, and mitigating the risk to wildlife, domestic animal, and human health.

Increased vigilance and enhanced surveillance activities were implemented (and are ongoing) in response to the initial incursions in the Atlantic provinces and, more recently, in Yukon, British Columbia, Ontario, Quebec, Manitoba, Alberta and Saskatchewan. Key reasons for the need for enhanced interdepartmental and interagency support and coordination in 2022-2023 towards a sustained and long-term Interagency Surveillance Program for Avian Influenza Viruses in Wild Birds (hereafter referred to as the Surveillance Program) include:

#### 5.1. Protection of Canada's wildlife

Given the range of bird species affected by this H5N1 strain, there is significant concern, nationally and internationally, regarding its potential impacts on biodiversity. With the initial incursions of the virus in wild birds on Canada's east and west coasts, further spread has rapidly occurred during spring migration. This virus will likely continue to spread rapidly within and between migratory flyways, particularly in locations of overlap where numerous species of migratory birds from multiple flyways converge and intermingle. The virus will likely continue to spread and evolve during fall migration as populations of birds, now with an influx of immunologically naïve juveniles, fly south to overwintering habitat. The scale of mortality events in wild birds in the eastern hemisphere, the USA, and Canada, the fact that affected locations include areas of international conservation importance for waterbirds, and the fact that globally threatened species have died from the virus, all indicate that this virus has the potential to pose risks to migratory birds in Canada, with significant conservation implications, particularly for species that are endangered, threatened, or vulnerable (Species at Risk; e.g., red knots, whooping cranes). In addition, there has been an unprecedented number of confirmed and/or suspected cases of HPAI H5N1 in mesocarnivores (e.g., red fox kits, striped skunk, American mink, harbour seals) in both Canada and the USA. However, it is still unknown whether the virus is transmissible among wild mammals.

Implementation of the Surveillance Program in 2022-2023 is necessary for understanding the geographic and seasonal spread of this emerging virus, understanding the range of Canadian species impacted, identifying potentially susceptible species or populations, and monitoring the evolution and changes in the dynamics of the virus as it continues to circulate through wild bird and mammal populations. The data generated are important for informing management decisions associated with mitigating or preventing negative consequences to wildlife, preventing spread in migratory birds, and protecting vulnerable or threatened populations or species.

#### 5.2. Protection of Canada's Poultry Industry, Export Trade in Poultry and Poultry Products, and Food Security

Data generated by the Surveillance Program will inform federal and provincial/territorial (P/T) regulatory bodies responsible for agricultural health, as well as poultry industry stakeholders, to help them make decisions about preparedness, biosecurity, and biosafety protocols in response to detections of H5N1 HPAIV or other AIVs of significance that may be circulating in wild birds locally, regionally, or within flyways. Current and credible data on H5N1 HPAIV and other notifiable AIVs circulating in wild birds in Canada can help prevent and reduce the substantial socioeconomic, export, and food security impacts of future outbreaks in commercial poultry as well as small flocks in Canada. Conversely, the absence of such data is an important risk factor for larger socioeconomic, export, and food security considerations.

#### 5.3. Protection of Human Health and Providing Public Assurance

The current H5N1 HPAI virus (clade 2.3.4.4b) which is emerging and spreading in Canada and the USA is still primarily an avian health issue. Based on available information published by the CDC (H5N1 Bird Flu Poses Low Risk to the Public (cdc.gov)), the risk to the public is considered to be low, however some individuals may be at higher risk of exposure due to job-related, recreational, or cultural activities or practices. The virus does not appear to be as pathogenic or as transmissible to humans from birds compared to earlier strains of HPAIV H5N1. Nevertheless, AIVs can rapidly mutate and recombine with other AIVs to generate novel viruses of significance, which may have higher capability of infecting mammalian species, including humans and/or an increased ability for transmission. The current strain appears to be impacting an unprecedented number of mesocarnivore species (i.e., red fox, striped skunk, American mink, harbour seals) in both Canada and the USA. Thus, surveillance in wild birds and other wildlife species is necessary not only to track geographic and seasonal spread of the virus, but also to track changes in the viral genome over time for traits that can signal higher zoonotic potential. (**Note**: *The US Centers for Disease Control and Prevention (USCDC) conduct research on zoonotic influenza viruses of public health concern. In conjunction with this work, the USCDC, in collaboration with other partners, has developed an Influenza Risk Assessment Tool (IRAT) that assesses the potential pandemic risk posed by influenza A viruses that currently circulate in animals but not in humans. A summary of results of Influenza A viruses assessed using IRAT, including the current strain, is available here*).

Due diligence and public assurance are also needed to demonstrate that potential avenues for spread in wild birds and populations of other wildlife (e.g., mesocarnivore), are being investigated, particularly given the heightened risk to domestic poultry and the potential for heightened public perception of risk (despite the current evidence that this lineage of virus poses a low risk to humans). Large-scale mortality caused by HPAIV could impact hunting and other economically and culturally important uses of wild birds and eggs, including those of traditional importance to Indigenous people. It is of particular importance to help provide assurance and build confidence in the safety and sustainability of cultural practices and traditional food systems, as well as to meet constitutional obligations to ensure Indigenous populations have access to wild foods (section 35 rights under the Constitution Act, 1982).

#### 5.4. Risk Analysis and Risk Communication

The risks posed by H5N1 AIV (clade 2.3.4.4b) and other AIVs of concern to Canada's wildlife and poultry industry, and the health of Canadians, can be best assessed on the basis of current information about the range of AIVs circulating within wild bird populations and other wildlife (e.g., mesocarnivores); the distribution of the virus between species and regions; their virulence; spatio-temporal variation in the virus among species; and the extent of interchange of viruses with Asia, Europe, Africa, and the Americas. Sustained, expanded, and timely surveillance of wild birds and other wildlife is essential to obtaining this information and will improve Canada's situational awareness of HPAI viruses and other AIVs of significance to wildlife, domestic animal, and human health. Surveillance will also generate information that is necessary for evaluating, reducing, or managing risk of transmission at the interfaces between wild birds, other wildlife, domestic animals, and humans.

#### 5.5. Strengthen and Build Upon Canada's Capacity for Surveillance for AIVs and Other Pathogens of One Health Significance

Expansion of Canada's Interagency Surveillance Program for Avian Influenza Viruses in Wild Birds will play a critical role in improving Canada's national AIV surveillance capacity, including field, laboratory, data management/analysis, and communications capacity, and will help inform decisions or approaches to management efforts or government policies. Strengthening Canada's capacity for surveillance by addressing current gaps and weaknesses, and sharing resources and capacity among agencies and jurisdictions, will be key to providing data and situational awareness in a timely and coordinated fashion, and will provide the foundation necessary for the establishment of sustainable and consistent P/T and national surveillance plans.

#### 5.6. International Contributions and Obligations

In the current climate of concern regarding potential pandemic HPAIV, Canada is obligated to contribute to the global understanding of AIVs in wild birds, as wild birds are the definitive global reservoir for AIVs. The World Organization for Animal Health (WOAH, formerly known as the OIE) Terrestrial Animal Health Code recommends that country members conduct surveillance of wild bird populations, including both morbidity/mortality and live bird surveillance (Chapter 10.4 of the OIE Terrestrial Code). As has been clearly demonstrated by the separate incursions of H5N1 HPAIV (clade 2.3.4.4b) on Canada's east and west coasts, Canada is geographically situated such that transmission of HPAIVs from the eastern hemisphere via transoceanic or transcontinental bird migration is possible. Furthermore, the rapid northward spread of HPAIV from the USA into Canada during spring migration highlights the importance of North American migratory bird pathways as routes of virus transfer between the USA and Canada. Therefore, Canada also must consider potential routes of virus transfer from the USA, Mexico, and Central and South America northward into Canada, and from Canada southward to the USA and the rest of the Americas. An established surveillance plan in Canada's wild birds can best achieve early detection of foreign HPAIVs of concern arriving into the Americas. Surveillance will allow Canada to meet national as well as international obligations for surveillance and reporting of H5N1 HPAIV (clade 2.3.4.4b) and other AIVs of significance. Data generated by Canada during fall migration can also assist our US and Mexico counterparts in making informed decisions on expanded or targeted surveillance in wild birds and in enhancing biosecurity protocols in their agricultural industries.

### 6. Canada’s Interagency Surveillance Program for Avian Influenza Viruses in Wild Birds: 2022-2023 Objectives

**Note***: P/T AI surveillance and implementation plans may also be developed with additional and/or adapted region-specific objectives*

- To detect the presence, distribution, and spread of H5N1 HPAIV (clade 2.3.4.4b), other AIVs of national or international significance to wildlife, domestic animal, and human health, in wild birds and other wildlife in Canada, and the timely detection of any new/emerging strains.
- To characterize AIVs detected in Canadian wild birds and other wildlife in order to:
    - Track the evolution of H5N1 (clade 2.3.4.4b) or other AIVs of concern as they circulate and spread among wild bird populations, and across the wild bird-poultry interface, in order to support early detection of mutations, recombinations, and reassortments that may impact inter-species transmission risk including zoonotic risk;
    - Compare AIVs detected in Canadian wild birds and other wildlife to those detected in domestic animals or humans in Canada and internationally.
- Compare the relative risk of H5N1 (clade 2.3.4.4b) transmission pathways in order to target interventions to the highest risk pathways.
- Detect wild bird and other wildlife morbidity and mortality caused by HPAIV infection.
- Identify the range of wild bird species and other wildlife that are impacted by HPAIV H5N1 or other AIVs of significance.
- Investigate the impacts of infection on (selected) populations of wild birds.
- Identify vulnerable species or populations at risk that may require implementation of mitigation/prevention strategies.
- Establish sustainable and consistent P/T and national surveillance protocols, and build upon and strengthen Canada’s integrated interagency, multi-jurisdictional network of field, laboratory, regulatory, and communications capacity needed to rapidly carry out AIV sampling, testing, genomic characterization, and reporting, on large volumes of samples during non-outbreak periods, as well as under emergent (i.e., outbreak) conditions.
- Establish reporting/communication protocols and platforms that can be effectively used by local, P/T, Indigenous, and national and international partners, particularly in response to detections of HPAIV H5N1 or other AIVs of concern, and to harmonize communications across Canada.
- Build on Canada’s inventory and archive of AIVs from wild birds and other wildlife to permit rapid retrospective analysis in response to disease outbreaks and to contribute to rapid epidemiological assessment.
- Meet national and international obligations for surveillance and reporting of avian influenza viruses.
- Use the data generated by the surveillance plan for modelling and/or to prioritize future targeted surveillance or research activities (e.g., to help inform surveillance targets for subsequent years, understand population level impacts on key species of wild birds, model the

dynamics and spread of the virus through bird populations and through migration, and to understand/predict changes over time).

### 7. Proposed Surveillance Components

#### 7.1. Morbidity and Mortality Surveillance for AIVs in Wild Birds and Other Wildlife

##### 7.1.1. Benefits:

Investigating morbidity and mortality events and diagnosing causes of morbidity and mortality through enhanced surveillance, including necropsy evaluation in conjunction with AIV testing, will improve our ability to detect the spread of HPAIV in susceptible wild bird populations and other wildlife (e.g., mesocarnivores). Enhanced morbidity/mortality surveillance will also improve our understanding of the **range of wild bird species and other wildlife** affected by this strain of H5N1 HPAI (or other AIVs of concern) and **the impacts of infection on** different species. Lastly, surveillance will improve our ability to identify other potential infectious or non-infectious causes of disease or mortality in wild birds and other wildlife in Canada.

##### 7.1.2. Approach:

Provincial and Territorial governments take primary responsibility for organizing the detection and collection of wildlife carcasses, their conveyance to participating veterinary diagnostic laboratories, and associated reporting.

- Procedures and the scale of activity may differ among P/Ts.
- CWHC regional centres will assist in these efforts as resources allow and on the request of P/T governments.
- Ensure that personnel collecting, submitting and testing wild carcasses are aware of and practice appropriate biosafety practices.

**Note***: The location of P/T reporting hotlines for morbidity/mortality events in wildlife has been provided in Appendix B.*

It is recommended that each P/T work with local P/T and federal partners in wildlife, agriculture, and public health sectors to establish protocols for communications, detection, collection and shipment of samples, as well as AIV testing and necropsy evaluation of morbidity/mortality events in wild birds and other wildlife by participating diagnostic labs (e.g., CWHC, P/T diagnostic labs, or university/commercial diagnostic labs). As H5N1 HPAIV continues to spread across P/Ts, interagency partners who have experienced outbreaks are encouraged to share their protocols and communications approaches so that they can then be adopted/modified by other P/Ts experiencing outbreaks. Protocols for communications and for detection, collection, and shipment of wildlife carcasses in each province and territory will aim to balance the need for enhanced surveillance (see below), and the limits on funding, resources, staff, and capacity in the field and in laboratories. In general, it is recommended that morbidity/mortality events be investigated for HPAIV regardless of the time of year, wild bird or other wildlife involved, or the number of samples already collected in the P/T. However, the prioritization of samples may be required in the event that a high volume of morbidity/mortality events overwhelm field efforts and diagnostic lab capacity during an outbreak. An example approach to the prioritization or triaging of diagnostic submissions of wild birds is available in Appendix D.

##### 7.1.3. Performance targets for morbidity and mortality surveillance:

The following performance targets are provided to guide actions, however, the ability of participants to meet these targets will be affected by their available capacities and budgets.

1. Diagnostic and communication targets
    a. Diagnostic testing – The goal is to minimize the time intervals between finding and submitting wild carcasses to laboratories, and between necropsy examination and completion of AIV related PCR tests and virus characterization. In previous years of enhanced morbidity and mortality surveillance in wild birds, the time frames recommended were as follows.
        i. The time between bird submission and confirmation of the presence of sub-type of H5 or H7 sub-type recommended not to exceed 2 weeks.
        ii. PCR testing of wild birds found dead recommended to be completed within five working days or sooner after the bird is received by the laboratory.
        iii. All samples that are positive by PCR test for H5 or H7 viruses will be sent to the National Centre for Foreign Animal Diseases (NCFAD), Winnipeg.

    The above timeframe targets are recommended for 2022-2023 in wild birds and other wildlife, where possible.

    b. Reporting results – The results of all diagnostic testing should be entered into the CWHC Interagency Surveillance database (where P/T-CWHC partnerships exist, and to be explored as an ideal approach in the instances where they do not) and wild bird results displayed on Canada's Avian Influenza dashboard (inclusion of results for other wildlife species to be explored as an ideal future direction). In previous years of enhanced morbidity and mortality surveillance, the time frame recommended for reporting was within 2 working days of initial PCR testing at regional laboratories and within 1 week of final confirmatory testing. A similar timeframe is recommended for 2022-2023 where possible.

2. Sample size, distribution, and species targeted

Given that the current strain of HPAIV has shown increased levels of mortality among wild bird populations in Europe, Asia, Africa, and North America to date, anticipated numbers for dead birds submitted for diagnostic evaluation in 2022 will significantly exceed submissions in previous years of morbidity/mortality surveillance.

Given increasing reports of detections in wild mesocarnivores (e.g., red fox, striped skunk, American mink, harbour seals), the range of species screened for AIVs should be expanded.

a. Exceed the wild bird sample sizes typical of the last 3 years of the program
    i. Appendix C outlines the number of birds tested for AIV as part of morbidity/mortality surveillance in previous years of Canada's Interagency Surveillance Program for Avian Influenza Viruses in Wild Birds, based on data extracted from the CWHC database.

b. Outreach efforts are encouraged to increase diagnostic submissions and enhance detections of morbidity and mortality events in wild birds and other wildlife (e.g., mesocarnivores, phocids):
    i. Implement regular reporting and collection procedures among ECCC-CWS and P/T field crews of wildlife carcasses that are found during regular program implementation;
    ii. Implement practices to increase vigilance among members of the public (e.g., social media, use of CWHC online reporting tool to report sick/dead birds);
    iii. Engage wildlife rehabilitation facilities and wildlife veterinary clinics to report morbidity and mortality events in wild birds and other wildlife that are submitted, and to increase testing on admission, as well as isolation of potentially infected individuals;
        1. **Note**: *ECCC has guidance related to intake of birds to wildlife rehabilitation facilities during a HPAI outbreak – Appendix I*

###### 7.1.4. Budget requirements:

The data in Appendix C have been provided for planning and budgetary purposes based on historical trends and anticipated increased submissions for wild birds in 2022-2023, and associated need for augmented necropsy and laboratory testing support. These numbers do not account for the anticipated increase in wild carcass submissions and diagnostic testing for other impacted species.

**Note**: *The number of wild bird carcasses tested for AIV through CWHC necropsy submissions during previous increased surveillance efforts due to HPAI events globally (2006-2009, and 2015) ranged from* ***an additional*** *1000-3200 per year. The number of birds submitted to the CWHC in 2022 to date (as of July 15th, 2022) is approximately 3.5 times the total submissions reported in 2021 (Appendix C).*

All P/Ts are asked to contribute to enhanced capacity for morbidity/mortality surveillance as part of the P/T in-kind contribution to the surveillance plan in 2022-2023 (via the CWHC, P/T labs, or other mechanisms). Federal agencies will also be asked to contribute. In some cases, it may be necessary to subsidize the field collection and shipping of specimens. Funding will be required to support increased diagnostic capacity/costs of participating diagnostic labs, to cover necropsy costs (gross necropsy and histopathology) and AIV testing. By supporting partners to enhance capacity to perform full necropsies to confirm cause of disease or mortality, we will be better able to assess the impacts of HPAIV on a broader range of wild species, and rule out other causes of morbidity and mortality.

##### 7.2. Surveillance for AIVs in Live and Hunter-harvested Wild Birds and Other Wildlife

###### 7.2.1. Benefits

Ongoing long-term surveillance for H5N1 HPAIV and other AIVs of concern in live and hunter-harvested wild birds and other wildlife in Canada will identify key species of wild migratory birds that are asymptomatic carriers; will provide essential information on the apparent prevalence, geographic distribution, seasonality, and spread; and will allow us to monitor and track the evolution of this virus over time. This information is critical for understanding, modelling and predicting the spatiotemporal spread of the virus in Canada; identifying areas of concern and potential hotspots (e.g., areas of flyway intersection or high densities of birds); estimating impacts of infection on survival (e.g., in conjunction with analysis of band recovery data); tracking genetic changes to the virus (genomic surveillance) that could signal increased or decreased virulence or transmissibility to birds or mammals including humans; and evaluating whether H5N1 HPAIV becomes endemic within wild migratory bird populations in Canada. The above data are important for informing management decisions associated with mitigating or preventing spread in migratory birds, and protecting vulnerable or threatened populations or species.

##### 7.2.2. Approach

Live wild animal surveillance can be conducted through the sampling of apparently healthy birds and other wildlife in conjunction with ongoing banding efforts or research studies led by interagency partners in ECCC, P/T partners and academia, and through sampling of hunter-harvested, or trapped wildlife.

1. Surveillance in live wild birds and other wildlife

Data generated from the surveillance of live wild birds and other wildlife will be key to providing the information outlined in section 7.2.1 (Benefits). Furthermore, given that the majority of wild birds sampled using this method will be banded, we can conduct band-recovery analyses to model the impacts of asymptomatic infection on band recovery rate and survival, particularly over multiple years of surveillance data.

Target species would be those as outlined below, with a primary focus on dabbling ducks and geese, and to a lesser extent other Anseriformes (e.g., diving ducks, sea ducks), Charadriiformes (e.g., gulls, shorebirds), and other species known to be affected by and/or carry the virus, depending on the region as well as sampling opportunities (e.g., mesocarnivores, cranes). The list of species is adaptive and changing as we learn more, within the scope of available sampling opportunities.

A standard operating procedure (SOP) for AIV sampling in live wild birds that was developed by ECCC in collaboration with CFIA and CWHC is available in Appendix G. The protocol provides information on the methodology and materials needed for AIV sample collection from wild birds, preferred storage options and conditions, and sample shipping and lab submission information. This SOP may need to be modified for use, as necessary, or for each region. It is important to ensure that personnel sampling, submitting and testing bird samples are aware of, and practice, appropriate biosafety practices and that all sampling is done in compliance with appropriate permitting and animal care procedures.

A similar SOP is not currently available for AIV sampling in live wild mammals but upon development, could be included as an appendix in an updated version of this plan.

2. Hunter-harvested birds and other wildlife (spring/fall)

Sampling of hunter-harvested birds and other wildlife (e.g., mesocarnivores) for AIV will allow us to supplement surveillance efforts in regions, locations, or time periods where sampling through live-capture is not possible. Furthermore, it will provide valuable information that will allow us to assess the presence of AIV in important migratory birds and other wildlife at the human-wildlife interface. It will allow us to 1) assess a wider geographic area beyond existing monitoring or research programs, particularly in northern or remote locations, and 2) assess AIV strains that people are most likely to come into contact with. Importantly, hunting and harvesting is often carried out in seasons where little banding or research efforts take place (e.g., spring), thus this represents a seasonal sample that is not possible via opportunistic sampling through research and monitoring programs (as outlined in 2.1). This type of sampling is recommended to be carried out in a variety of ways, including, but not limited to:

- Partnering with P/Ts and regional programs to collect AIV samples through existing relationships with hunters (e.g., sampling in northern Quebec via northern research nodes).
- Undertaking sampling through other science programs that currently work with hunters to sample wildlife (e.g., contaminants monitoring via hunter harvest in Nunavut and Nunatsiavut).
- Establishing sample drop-off locations for hunters to submit samples via regional offices and activities (e.g., similar to drop-off programs for chronic wasting disease monitoring and oil spill reporting for oiled birds).

***Note****: Known live wild bird and hunter harvested sampling efforts have been reviewed and the species/locations captured through these efforts which have been identified as target species/locations, are available in a table in Appendix F. This table is not a comprehensive list of all target species/locations, rather it lists those target wild bird species which are expected to be sampled (in sufficient numbers) by existing field operations planned for 2022-2023. This list is evergreen and will continue to be populated/revised as more information becomes available, in alignment with the objectives as outlined above and in consideration of the available resources (e.g., budget and capacity). It is requested that any additional field opportunities not represented on this list, be submitted to [redacted].*

***Note****: A list of existing sampling efforts is not currently available for other wildlife*

In general, the selection of target species, locations, and time periods for live and hunter-harvested surveillance should be conducted using risk-based surveillance approaches:

***Note****: The following provides general guidelines, however, these criteria should be adapted to the regional context and in consideration of region-specific knowledge*

1. Target Species

- Should focus on species that may potentially be critical for the transmission, spread, maintenance, and evolution of this HPAIV and other AIVs of concern (e.g., Anseriformes, with an emphasis on dabbling ducks). This includes species that are generally (or often) asymptomatic when infected, as they are more likely to become reservoirs for this HPAIV, and may play a role in the generation of novel AIVs of concern through mutation, recombination and reassortment.
- May also consider other species that appear to be vulnerable to infection (based on current morbidity and mortality data; e.g., gulls, cranes, seabirds, mesocarnivores) to understand whether these species survive infection, and whether the virus has the potential to circulate in these populations.
- Both sexes and age categories should be included, however in wild birds, given that hatch-year ducks are 2-5 times more likely to be positive for LPAIV compared to adults prior to fall migration (Papp et al, 2017; Nallar et al, 2015; Nallar et al, 2016), an increased emphasis on sampling of hatch-year birds should be considered in order to increase the likelihood of detection in a particular location, species, or population.
- Most hunter/harvest collections of migratory birds will be waterfowl across Canada, with some other species included that will vary by region. This includes Anseriformes and Charadriiformes (e.g., murres and other species of seabirds in NL and Nunavut). Given that both of these groups (Anseriformes and Charadriiformes) are highly susceptible to the current strain of HPAIV, hunter/harvester collections of these groups should be prioritized.

2. Target Locations

- Should include locations that are potential key areas or “hot spots” for transmission and spread across Canada. This can include targeting locations that have high wild bird population densities, a high diversity of species that intermingle, that may have multiple flyways overlapping, which often occurs on large wetlands or watersheds. Multiple studies conducted in Canada using previous surveillance data have demonstrated that infection with LPAIV is positively associated with waterfowl population densities regionally and across Canada (Papp et al, 2017, Nallar et al, 2016).
- In some instances, consideration for targeting sites that are near locations of importance to endangered, threatened or vulnerable species or populations to determine whether HPAIV has spread near important breeding areas, stopover sites or staging areas.
- Another consideration is targeting locations of importance to the interface between wild birds, domestic poultry, and human health, depending on the objectives. A recent study using satellite-marked waterfowl revealed that the occurrence of AIV outbreaks in commercial poultry in North America was associated with increased waterfowl occurrence or residence time in the vicinity of farms (Humphreys et al 2020).
- Finally, surveillance at other locations, e.g., in response to positive detections in wildlife or poultry (see other surveillance strategies below)

3. Target Time Periods or Seasons

- Sampling campaigns should focus on critical times during which waterfowl or other potentially susceptible species gather in large numbers or densities, and/or share sites with multiple species, where they moult or stage prior to migration, or at key stopover sites during migration. Different periods of time or seasons within the year and across multiple years are important for detecting seasonal and annual patterns.
    - **Spring migration** – to evaluate the distribution and track the spatiotemporal spread of virus as migratory birds move northward from the USA into Canada, and for some species, into Canada’s North.
    - **Post-breeding season just prior to fall migration** – to evaluate the distribution and spatiotemporal spread among wild birds, particularly with the influx of immunologically naïve juveniles into the population. A study led by ECCC examined Canada’s wild bird AIV surveillance data in dabbling ducks sampled from 2005 to 2011, and demonstrated that peaks in LPAIV infection occurred in late August to end of September in Eastern Canada (Ontario, Quebec, and Atlantic Canada), and in mid-August in the prairie provinces (Papp et al, 2017). Another study led by ECCC, in collaboration with USDA-APHIS and USGS, combined wild bird influenza surveillance data from Canada and the USA to examine spatiotemporal patterns of AIV infection in blue-winged teal (a long distance migratory dabbling duck) at the continental scale, and demonstrated the apparent prevalence of AIV in this species peaked during late summer staging (July-August), and that seasonal cycles and spatial variation in the apparent prevalence of AIV are largely driven by the dynamics of AIV infection in hatch year birds (Nallar et al, 2015).
    - **Fall migration** – to evaluate the distribution and track the spatiotemporal spread of virus as migratory birds move southward to the USA, Mexico, and further south.

- There can also be surveillance at other periods of time, e.g., in response to positive detections in wildlife or poultry (see other surveillance strategies below)

##### 7.2.3. Performance targets for live and hunter harvested animal surveillance

The following performance targets are provided to guide actions, however the ability for participants to meet these targets will be affected by their available capacities and budgets.

1. Diagnostic targets and communications
    a. All tests should be completed and results entered into the CWHC Interagency Surveillance database and displayed on Canada's Avian Influenza dashboard Survey database within 4 weeks of field collection.

2. Sample Size and Distribution

Surveillance of live, apparently healthy birds should be enhanced compared to previous years, in spring (where possible) and in late summer/fall in Canada. Given that there have been at least two separate incursions of H5N1 HPAIV into Canada, and given the rapid spread of the virus across Canada and the USA, we recommend sampling in all four flyways. **Note***: Appendix C outlines the number of birds tested for AIV as part of live bird surveillance in previous years of Canada's Interagency Surveillance Program for Avian Influenza Viruses in Wild Birds (formerly referred to as Canada's Interagency Wild Bird Influenza Survey), based on data extracted from the CWHC database.*

Surveillance of other wildlife (e.g., mesocarnivores) should be initiated where possible.

Opportunistic sampling will be the focus for live wild bird and other wildlife surveillance in 2022-2023. To capitalize on ongoing activities across the country and maximize our efforts, sampling should be conducted opportunistically in conjunction with waterfowl banding activities and research/surveillance studies led by various interagency partners in ECCC, the P/Ts, CWHC, and academia.

A power analysis was performed to estimate sample size requirements to detect the presence or absence of HPAIV in wild bird populations in Canada. The results of the analysis are summarized below, divided into two scenarios according to the range of wild bird species sampled. The detailed power analysis is provided in Appendix E.

**Scenario 1:**

Species: Heavily focused on waterfowl species with inclusion of small numbers of other relevant species in relevant locations

Objective: Detect the presence or absence of HPAIV assuming roughly 2% prevalence (as reported in the USA at the time the power analysis was completed), with a 95% confidence interval

Rationale: The total sample size represents 200 samples collected from selected waterfowl species (dabbling ducks, geese) at multiple sites within in each flyway (using criteria described in section 2.1), and based on waterfowl population estimates from the latest status report.

Total sample size: at least 8,400

**Scenario 2:**

Species: Waterfowl, other Anseriformes, and Charadriiformes

Objective: Detect the presence or absence of HPAIV assuming roughly 2% prevalence (as reported in the USA at the time this document was prepared), with a 95% confidence interval

Rationale: Scenario 1 does not include an emphasis on other species (e.g., Charadriiformes or other potential target species). If sampling efforts of Charadriiformes (or other species) are increased, then sample size estimates for live bird AIV surveillance in Canada increase according to the same rationale provided in scenario 1.

Total sample size: at least 11,600

#### 7.3. Environmental Surveillance for AIVs

##### 7.3.1. Benefits

The collection of environmental samples (e.g., feces, sediment/water samples) can supplement or even replace live animal sampling efforts for time periods or regions in which live animal sampling is limited, or in conjunction with sampling depending on specific objectives within a particular region.

Fecal sampling (through collection of fresh fecal material with a swab into a vial containing viral transport medium, similar to cloacal swabbing) can be used to supplement live and hunter-harvested wildlife surveillance. This method can be particularly useful during spring sampling and fall migration when there are few banding or research activities taking place. Fecal sampling in spring, particularly in advance of any detectable mortality event, can serve as a method for early detection of the presence of the virus in populations. Selection of key sites would follow criteria outlined in section 7.2.2.

Genomics-based approaches (targeted resequencing) to analyze environmental samples (e.g., water/sediments from ponds used by waterfowl) can also be used to supplement live bird sampling efforts. This approach has been shown to be useful for detecting the presence of strains even prior to strains being detected in wild birds using a site, particularly at sites where the prevalence may be too low to detect in wild birds (Himsworth et al, 2019). This method may be applicable for investigating transmission pathways between wild birds (or environment) to and from poultry farms during outbreaks (e.g., see section 2.2.4). This method could also be considered for identifying and characterizing AIV strains present in important migratory bird habitats or zones in order to inform modelling about future outbreaks, as AIV is known to be able to overwinter in soil and water matrices (Morin et al. 2018; Mihai et al. 2011; Shoham et al. 2012).

##### 7.3.2. Approach and Performance Targets

The approach and performance targets are still an active area of discussion. Key areas of focus include:

- The development of a list of key environmental sites to be sampled according to season
- The development of protocols for the sampling of feces, water and soil, along with collection of measurements on relevant environment variables such as temperature and pH
- Use of these environmental data to understand the role of the environment in transmission between poultry (commercial and backyard) and wild birds (and other wildlife), and the potential for overwintering of the virus.

#### 7.4. Other Surveillance Strategies: Targeted Animal and Environmental Surveillance Surrounding Positive Detections

##### 7.4.1. Benefits

Targeted sampling of waterfowl, and potential "bridging hosts," on or around commercial farms can be considered in response to the detection of HPAIV in wild birds in the region of commercial poultry operations, or in response to the detection of HPAIV on a poultry farm, to further understand the bidirectional pathways of transmission at the wild bird-poultry interface. This can inform biosecurity measures to reduce transmission of H5N1 HPAIV or other AIVs of concern from wild birds (or the environment) to poultry, and vice versa, thereby potentially preventing outbreaks in poultry as well as in wild birds (depending on the direction of transmission).

##### 7.4.2. Approach

Sampling around positive detections (either domestic or wild species) should be done with partners including P/Ts, CFIA, and industry. Each situation will be different, and enhanced/targeted surveillance should be carried out within the context of the positive detections. This type of sampling should be adaptive to enhance surveillance in new species or locations where HPAIV positives have been detected. This includes positive domestic premises to better understand routes of transmission that are currently evading biosecurity measures that have already been implemented. This type of sampling should also consider avian species that may be asymptomatic vectors, and non-avian species that could be potential carriers or mechanical vectors of AIV. The purpose is to better understand how HPAIV may enter and be maintained in a region, specifically if this applies to domestic poultry farms where biosecurity measures are in place, but HPAIV infections are still occurring.

**Target premises/locations to consider for enhanced/targeted surveillance**

- Any HPAIV positive domestic bird facility and its surroundings.
- Any site where a large-scale unusual mortality event of migratory birds has occurred with a confirmed HPAIV positive detection.

**Sampling types for enhanced/targeted surveillance**

- Target susceptible bird species at the poultry-wildlife interface in locations with positive detections (e.g., ducks and waterfowl).
- Consideration of adding new avian species (potential "bridging species") depending on location or concerns relevant to the region. For example, passerines and corvids could be considered where they may contribute to the movement of HPAIV near locations with positive detections, as they may serve as carriers or mechanical vectors.
- Mammalian sampling may be considered where mammals may interact with soil or feed that birds are then exposed to (i.e. barn cats, rodents, and other potential bridging species).
- Environmental/soil sampling can be carried out where animals may be moving soil, feathers or fur around an area (e.g. for nesting material).
- Water/sediment sampling of wetlands or ponds on properties, especially if frequented by aquatic birds, and if the water is used by the farm.

***Note:*** *A targeted surveillance initiative is currently being designed in response to stakeholder queries, that will inform whether barn swallows nesting on barns in areas with HPAIV could and are likely to, act as a "bridging host" at the wild bird-poultry interface. Currently, this surveillance is expected to take place in Ontario and Nova Scotia via existing partner programs that are focused on barn swallows in agricultural landscapes.*

#### 8. Sample Diagnostics, Results Reporting and Communication

Samples from wild bird species to be tested for influenza will consist of both a cloacal and an oral swab combined in a single vial containing virus transport medium using appropriate sampling protocols. Participating laboratories will test each sample by PCR for the matrix protein gene, and matrix positive samples will also be tested by PCR for H5 and H7 subtypes (**Note**: *in some instances all samples may be tested by PCR for H5 and H7 subtypes*). Currently at a minimum, all samples that are positive for H5 or H7 should be sent to the National Centre for Foreign Animal Diseases (CFIA, Winnipeg, MB) for further viral characterization (e.g, subtyping, virus isolation, sequencing).

A list of regional laboratories (i.e., Canadian Animal Health Surveillance Network (CAHSN) partner laboratories) can be found in Appendix H.

The results of all Interagency Avian Influenza Surveillance activities in wildlife species should be submitted to the CWHC which maintains a national database of AIV sample collection information and results for all participating program partners. The CWHC sends out bi-weekly reports via email and the information is also uploaded on their website. The data from wild birds will also be displayed on Canada's Avian Influenza dashboard which is updated weekly. Links to the CWHC and Canada's Avian Influenza dashboard can be found in Appendix B.

All collaborators should strive to communicate results in a timely fashion to ensure the data reflect the current situation.

The Interagency Wild Bird Surveillance Working Group will generate an annual report that summarizes the surveillance activities, testing results and surveillance performance relative to the targets outlined in the 2022/23 Implementation Plan.

The coordination and establishment of a system to share data across the US border in an efficient and timely manner is an active area of discussion (e.g., link between the Canadian Avian Influenza dashboard and WHISpers).

### Appendix A: Anticipated Partners

This surveillance plan is intended to build and enhance Canada's One Health capacity through collaboration among wildlife, agriculture, and public health agencies within federal and provincial/territorial, and Indigenous governments and with the Canadian Wildlife Health Cooperative, academia, hunting/trapping organizations, and other organizations.

**Primary Federal Government Partners:**

- Environment and Climate Change Canada
    - Science & Technology Branch
    - Canadian Wildlife Service
- Canadian Food Inspection Agency
- Public Health Agency of Canada
- Parks Canada Agency

**Primary Provincial/Territorial Government Partners:**

- Provincial/Territorial Departments responsible for Wildlife and Protected Areas
- Provincial/Territorial Departments responsible for Agriculture (Animal Health)
- Provincial/Territorial Departments responsible for Public Health

### Appendix B: Important Links and Contact Information

| Name | Link |
|---|---|
| Canadian Wild Birds HPAI Dashboard | National Avian Influenza - Wild Positives (arcgis.com) |
| US Wildlife Health Information Sharing Partnership Event Reporting System (WHISPers) | WHISPers (usgs.gov) |
| Canadian Food Inspection Agency (CFIA) | Canadian Food Inspection Agency - Canadian Food Inspection Agency (canada.ca) |
| Public Health Agency of Canada (PHAC) wild birds and avian influenza handling guidelines | Wild birds and avian influenza – Handling guidelines - Canada.ca |
| Canadian Wildlife Health Cooperative (CWHC) | CWHC-RCSF :: Canadian Wildlife Health Cooperative - Réseau canadien pour la santé de la faune |
| United States Department of Agriculture, Animal and Plant Health Inspection Service (USDA APHIS) | USDA APHIS \| Avian Influenza |
| US Centers for Disease Control and Prevention (USCDC) Influenza Risk Assessment Tool (IRAT) | Information on the tool: Influenza Risk Assessment Tool (IRAT) \| Pandemic Influenza (Flu) \| CDC<br><br>Results: Summary of Influenza Risk Assessment Tool (IRAT) Results \| Pandemic Influenza (Flu) \| CDC |

**Provincial/Territorial reporting hotlines for AIV morbidity/mortality events in wildlife can be found at the following link:**

Avian influenza in wild birds - Canada.ca

#### Appendix C: Number of Wild Birds Tested for AIV as Part of Morbidity/Mortality and Live Bird Submissions in Previous Years of Canada's Interagency Surveillance Program for Avian Influenza Viruses in Wild Birds (formerly referred to as Canada's Interagency Wild Bird Influenza Survey)

***Note**: The data in the table below have been provided for planning and budgetary purposes based on historical trends and the anticipated need for increased submissions of wild birds for 2022-2023, compared to historical submission numbers. The data for morbidity/mortality surveillance was obtained from the following link: CWHC-RCSF :: Canadian Wildlife Health Cooperative - Réseau canadien pour la santé de la faune*

| Sampling year | Morbidity/mortality surveillance in wild birds | Live bird surveillance |
|---|---|---|
| 2005 | 165 | 4319 |
| 2006 | 2863 | 9687 |
| 2007 | 3354 | 7435 |
| 2008 | 3019 | 2261 |
| 2009 | 2407 | 4875 |
| 2010 | 1845 | 5885 |
| 2011 | 1864 | 4858 |
| 2012 | 1664 | 951 |
| 2013 | 1392 | 1218 |
| 2014 | 1544 | 1324 |
| 2015 | 2288 | 2999 |
| 2016 | 1304 | 3990 |
| 2017 | 1207 | 1187 |
| 2018 | 1003 | 728 |
| 2019 | 806 | 1038 |
| 2020 | 840 | 278 |
| 2021 | 940 | 2319 |
| 2022 year to date (as of July 15$^{th}$) | 3503 | Sampling currently underway |

### Appendix D: Example Prioritization Decision Tree for Morbidity/Mortality Surveillance in Wild Birds

An example approach to the prioritization or triaging of diagnostic submissions during AI surveillance in wild birds. The approach presented was proposed by CWHC (Atlantic region) in early 2022 and included the following guidance:

- Bird species which frequent aquatic or wetland habitats (e.g. ducks, geese, swans, seabirds, gulls, terns, shorebirds, loons, cranes, rails, herons, pelicans, grebes, cormorants, gannets, etc.); sample individuals with evidence of trauma or starvation if capacity allows.
- All raptors.
- Additional scavenger species (crows, ravens, vultures, etc.)
- Deaths of 3 or more individuals of any other species in the same location (with no signs consistent with trauma or with a different disease, e.g., salmonellosis, trichomonosis, etc.).
- All birds with neurological signs.
- Additional species/groups should be added to the list above if confirmed to be infected in other regions of North America.
- Additional species or repeated sampling of specific species in the same location may be needed for additional epidemiological information, or to meet research needs.
- Sampling of wild birds from rehabilitation facilities can also follow these same triage criteria.
- Furthermore, any species listed as threatened, vulnerable, or of concern should also be submitted for diagnosis if found dead.

***Note****: triage criteria may change from the onset of an outbreak to late in the outbreak as the objectives may change*

#### Appendix E: Power Analysis for Live Bird Surveillance

**Purpose:** estimate the number of samples needed in 2022 from live captured birds to evaluate AIV in Canadian populations/species of birds most commonly implicated in the epidemiology of AIV, across all four flyways.

**Summary of findings**

- In order to confirm AIV presence/absence in 2022 across bird species, flyways, and populations with high probability, ECCC should aim to process **at least** 8,400 samples from waterfowl in 2022.
- If additional Anseriformes species and Charadriiformes species are sampled, the target sample size increases to **at least** 11,600 samples.
- In order to assess the prevalence of AIV in migratory bird populations, approximately 4,700 samples from each population numbering one million individuals is needed.

**Assumptions**

- This is for live captured and hunter-harvested birds only
- This does not account for any morbidity/mortality sampling

**Calculating potential sample sizes needed**

Minimum sample sizes required to detect the presence/absence of AIV within a population at 1-2% virus prevalence rates with 90 or 95% confidence intervals, were calculated using the following formula (Sample Size Calculator (sruc.ac.uk)):

$$n = \left(1 - (1-p)^{\frac{1}{NP}}\right) \times \left(N - \frac{NP-1}{2}\right)$$

where:

*p* is the confidence level

*N* is population size

*P* is the prevalence

Based on these results, we estimate that approximately 200 samples should be collected from each sampling effort unit. This accounts for some failed samples, and aims to provide data for varying prevalence and population sizes.

| Population size | Population prevalence of HPAI | Confidence level (%) | Minimum sample size needed |
|---|---|---|---|
| 1,000 | 2% | 95 | 138 |
| 1,000 | 2% | 90 | 108 |
| 10,000 | 2% | 95 | 148 |
| 10,000 | 2% | 90 | 114 |
| 100,000 | 2% | 95 | 149 |

| | | | |
|---|---|---|---|
| 100,000 | 2% | 90 | 114 |
| 500,000 | 2% | 95 | 149 |
| 500,000 | 2% | 90 | 114 |
| 500,000 | 1% | 95 | 298 |
| 500,000 | 1% | 90 | 229 |

**Historic sampling efforts for AIV**

- Between 2005 and 2021 the average number of AIV samples collected from live birds in Canada was 3,058. This is reflective of all efforts reported to the Canadian Wildlife Health Cooperative (CWHC), and does not include all AIV sampling in Canada (e.g., other research studies in Canada).
- The years with higher surveillance efforts when global concerns around HPAI include 2005 to 2011, 2014/15 and 2021.
- The average sample numbers for these years of increased surveillance is 5,405.

**Estimate based on migratory gamebird populations across Canada**

- The current strain of highly pathogenic AIV is being detected at rates of approximately 2% in samples collected from live-trapped and hunter-shot birds in the US in 2022.
- The most recent population status of migratory game birds in Canada provides population estimates of numerous Anseriformes for the year 2019, which can be used to calculate the minimum sample size needed to confirm AIV presence/absence with high probability, across species and spatial scales
- Given the apparent AIV prevalence of 2% reported in the US, it is estimated that 150-200 samples are needed from most populations in order to confirm virus presence/absence with high probability.
- The calculations described above have demonstrated that at 2% AIV prevalence and 95% confidence levels, the statistical power provided by minimum sample sizes of approximately 150 is sufficient to detect AIV in waterfowl populations across Canada (based on estimates for the number of breeding pairs in each population ranging from 2,950 to 9,420,000 individuals).
- Increasing the minimum sample size to approximately 200 for populations would account for sampling and diagnostic errors, such as improper sample collection/storage, sample degradation/contamination, failed PCR reactions, inadequate sample volume/RNA template, and allows for imperfect test sensitivity.
- This population estimate likely overestimates capture sample numbers as not all waterfowl populations can be captured and tested, and underestimates possible sample numbers as it does not capture all waterfowl populations in Canada (i.e. only a few populations in Arctic Canada are included in the report).

**The estimated sample sizes provided below are based on (known) existing programs that include handling of Anseriformes and Charadriiformes with the following considerations:**

- The current strain of HPAI has about 2% prevalence in Anseriformes populations based on data from the US in early 2022.
- Existing programs handle many species, and some in large numbers.
- Based on information collected by CWS on the opportunities to swab in 2022, there are programs existing in each flyway.

- If we take each program as an independent sampling unit, we would have 200 swabs per unit, per species.

**Scenario 1:**

Species: Heavily focused on waterfowl species with inclusion of small numbers of other relevant species in relevant locations

Objective: Detect the presence or absence of HPAIV assuming roughly 2% prevalence (as reported in the USA at the time this document was prepared), with a 95% confidence interval

Rationale: The total sample size represents 200 samples collected from selected waterfowl species (dabbling ducks, geese) at multiple sites within in each flyway (using criteria described in section 2.1), and based on waterfowl population estimates from the latest status report.

Total sample size: at least 8,400

**Scenario 2:**

Species: Waterfowl, other Anseriformes, and Charadriiformes

Objective: Detect the presence or absence of HPAIV assuming roughly 2% prevalence (as reported in the USA at the time this document was prepared), with a 95% confidence interval

Rationale: Scenario 1 does not include an emphasis on other species (e.g., Charadriiformes or other potential target species). If sampling efforts of Charadriiformes (or other species) are increased, then sample size estimates for live bird AIV surveillance in Canada increase according to the same rationale provided in scenario 1.

Based on available information about existing sampling efforts including Anseriformes, Charadriiformes, and other target species across Canada, the following number of samples collected in the following regions is proposed:

- Atlantic – 5 sampling effort units = 1000
- Ontario – 10 sampling effort units = 2000
- Quebec – 10 sampling effort units = 2000
- Prairies – 20 sampling effort units = 4000
- Arctic – 8 sampling effort units = 1600
- BC – 5 sampling effort units = 1000

Total sample size: at least 11,600

### Appendix F: List of Live Wild Bird Regional Sampling Opportunities and Target Species

***Note:*** *Table 1 (summary) and Table 2 (detailed) include a list of known sampling efforts that capture wild bird species/locations which have been identified as target species/locations. This list is not a comprehensive list of all target species or locations, rather it represents those target wild bird species and locations which are expected to be sampled (in sufficient numbers) by existing field operations and/or research projects planned for 2022-2023 that are currently known to us. This list is evergreen and will continue to be populated/revised as more information becomes available and in alignment with the objectives as outlined above and in consideration of the available resources (e.g., budget and capacity).*

*Additional field opportunities that are not represented on this list, can be forwarded to [redacted].*

Table 1. Summary of live bird sampling opportunities by season (hunter-harvested not included) as of May 2022

*[Table contents redacted]*

Table 2. Detailed list of live bird and hunter-harvested sampling opportunities from which Table 1 was generated, as of May 2022

*[Table contents redacted]*

### Appendix G: AIV Sampling Protocol in Live Wild Birds

***Note***: *The following SOP is an evergreen document that is being updated as needed.*

**Avian Influenza (AIV) National Surveillance in Wild Birds:**
**AIV Sampling, Storage, and Shipping to Regional Labs**

**Standard Operating Procedure**
**For partners collecting and submitting avian influenza samples from wild birds, as part of *Canada's Interagency Surveillance Program for Avian Influenza Viruses in Wild Birds,* to regional laboratories for preliminary PCR screening.**

#### 1.0 Purpose

To outline avian influenza (AIV) sampling, storage and shipping procedures for the submission of samples to regional laboratories for matrix (M), H5, and H7 gene preliminary PCR tests. This standard operating procedure (SOP) covers the methodology for AIV sampling collection from wild birds, preferred storage options and conditions, and sampling shipping and lab submission information, and includes a list of materials required in the field.

#### 2.0 Procedures

##### 2.1 Materials Required for Field Sampling

- Swabs (Sterile synthetic applicator swabs to be used for the collection of oral samples and cloacal samples)
- Vials (16x100 mm tubes containing 3.0 mL of virus transport medium - https://www.copanusa.com/wp-content/uploads/2019/08/UTM-Brochure.pdf)
- Cryomarker or permanent marker (Sharpie)
- Ziploc bags
- Hard cooler with freezer packs to keep samples cold during the day
- Latex gloves
- Hand sanitizer
- Other Personal Protective Equipment (PPE) in accordance with your institutional requirements
- Field log sheet

The materials will be shipped to the location received through email communication. A sample field log sheet will be distributed through email.

##### 2.2 Sample Collection Procedure

From each bird, one (1) cloacal swab and one (1) oral swab are to be collected (two [2] swabs per bird) and placed into a one (1) UTM vial.

###### 2.2.1 Cloacal Swabs

a. Choose the appropriate swab size for the species. Open the plastic applicator swab envelope on the 'stick' end (do not touch the sterile polyester-end). **Keep swab as sterile as possible; avoid contact with everything other than what is being sampled**. If the applicator end of the swab touches anything other than the intended sample, discard and use a new swab.

b. Remove the swab from the envelope and **gently** insert into cloaca of the bird (about 1 cm), swab along the mucosa, rotate it to collect a sample of excreta (getting fecal matter onto swab is good), and remove the swab.

c. Insert the swab about 3/5ths into the vial, and **carefully** break the swab off into the vial by prying the swab against the rim of the vial. **Do not use scissors to cut the swabs off**, as this could cause spillage and contamination of the samples with the contents of other vials or swabs (cross-contamination).

- It is important to try to avoid cross contamination as much as possible.

d. Close the tube tightly immediately after putting the swab in it, and keep chilled according to storage instructions in section ***2.3–Sample Storage Requirements.***

###### 2.2.2 Oral Swabs

a. Open the applicator swab envelope from the 'stick' end being careful to keep the swab as sterile as possible as described above. Remove the swab and insert it into back of the mouth, at the level of the larynx and back of the tongue, and gently swab across the back and roof of the mouth, and over **(not inside)** the choanal slit.

b. Insert the swab about 3/5ths into the vial, and carefully break the swab off into the vial by prying the swab against the rim of the vial. Avoid cross-contamination between samples as much as possible as described above.

c. Close the tube tightly immediately after putting the swab in it, and keep chilled according to storage instructions described in section ***2.3–Sample Storage Requirements***.

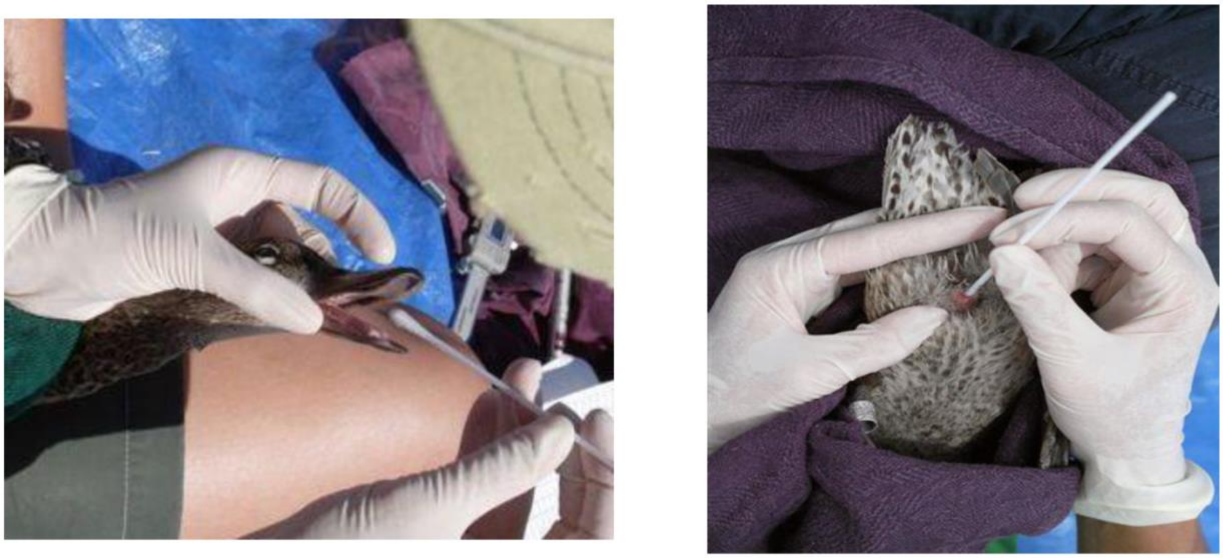

**Figure 1.** Example of oral (left) and clocal (right) sample collection from a single wild bird. For each bird, one (1) swab from the oral cavity, and one (1) swab from the cloaca, are to be carefully placed into a single vial.

###### 2.2.3 Vial Labels

A unique AIV ID/sample number should be written on each tube using a cryomarker or permanent marker. It is important to ensure that the AIV ID/sample number can be matched with the corresponding band number.

Please label each tube according to the following format:

AIV-year-province code-number

e.g., AIV-2021-SK-001

*Note: if you have multiple crews collecting samples in the same province, please coordinate your coding accordingly so there are no duplicates (e.g., one crew takes 1-100, the next 101-200 etc.)

###### 2.2.4 Field log datasheet

Please complete the AI field log sheet by recording:

- **Date**: Record the date in the following format (YYYY-MM-DD).
- **Province**: Record province code.
- **Coordinates**: Record lat/long.
- **Location**: Record the name of the lake/wetland.
- **Banding Station Name**: Record name if applicable.
- **AIV ID/Sample Number**: A unique ID should be written on each tube and recorded on the AI field log sheet.
- **Band Number:** For cross referencing purposes record the band number of the bird you are sampling if applicable.
- **Species**: Record the species of the bird you are sampling
- **Sex**: Record M (male), F (female), or U (unknown).
- **Age**: Record the age of the bird as U (Unknown), HY (Hatch Year), AHY (After Hatch Year), SY (Second Year), ASY (After Second Year).
- **AIV Swab**: Record sample source. O/C (oral and cloacal in one tube), O (oral only), or C (cloacal only). *Note: both oral and cloacal samples are requested where possible.
- **Sampler**: Record initials or name of individual that obtained the sample.
- **Capture Method**: Record how the bird was captured (i.e., airboat, dip net, baited trap, etc.).
- **Notes**: Record any additional notes if applicable.

Please email the AI field log sheet to the AIV surveillance coordination contact.

A printed copy can also be included with the samples during return shipping.

##### 2.3 Sample Storage Requirements

- The sample vials of commercial transport medium (e.g. Copan Universal Transport Medium (UTM-RT®) System, catalog # 3C047N; Puritan UniTranz-RT Transport System, catalog # UT-361) are not required to be chilled or frozen until after use, however they should at minimum be kept at **room temperature** to keep the integrity of the transport medium (e.g., keep in a cooler with freezer packs if in a truck on a hot day).
- Sample vials provided by the Prairie and Northern Wildlife Research Centre in Saskatoon contain a viral transport medium made in-house at the University of Saskatchewan, and must remain cool prior to use. Store vials at 4◦C until sampling.
- Vials **must** be kept chilled after use. A cooler with freezer packs is sufficient during the day. At the end of the day or as soon as possible thereafter, tubes should be transferred to a -20◦C freezer, -80◦C freezer, or if available, a cryoshipper filled with liquid nitrogen.
    - If there are logistical issues that prevent freezing the samples right away or concerns with keeping the samples in a frozen state, samples can remain cool (refrigerated 2-5◦C or on ice) for up to 5 days after they are obtained after which they must be transferred to a -20◦C freezer, -80◦C freezer, or a cryoshipper filled with liquid nitrogen.

- The appropriate Transportation of Dangerous Goods (TDG) training is required to send, carry, or receive a cryoshipper filled with liquid nitrogen. It is understood that cryoshippers are not feasible for many field crews due to the specific TDG training required, their size, and the unavailability of liquid nitrogen.
- AI virus is very susceptible to freeze/thaw cycles therefore it is recommended that all samples (if not being shipped to the laboratory on ice within the same day) be **frozen as soon as possible**. **However, once frozen the samples must remain frozen and never thawed until testing at the laboratory.**
- Ensure that the vials are **tightly closed** after use to prevent leakage, and place securely into the cooler.
    - It is recommended to keep vials in a Ziploc bag that is then placed in the cooler/freezer. Please label the Ziplock bag with collection information (i.e., sampling location, date).

#### 2.4 Shipping Instructions

##### 2.4.1 Shipping Temperature Conditions

**Sample vials must remain frozen during return shipping (or chilled on ice-packs if shipping to the laboratory same-day).** The most important consideration when shipping is to limit the amount of freeze-thaw cycles to preserve sample integrity. Not all shipping and packaging conditions may be feasible, as field crews may not have access to cryoshippers or -80°C freezers. However, the recommended shipping and packaging conditions, in order of most **preferred (1) to least preferred (4)** are:

**1.** Shipping immediately (same-day) on regular ice (i.e. refrigerating and never freezing). AIV samples can remain cool (2-5°C or on ice) for up to 5 days after they are obtained.
**2.** Freezing at -80°C and shipping on dry ice to prevent samples from thawing during shipment.
**3.** Freezing at -20°C and shipping on dry ice to prevent samples from thawing during shipment.
**4.** Freezing at -20°C and shipping on regular ice (ice packs), **ensuring that samples do not thaw during shipping.**

##### 2.4.2 Packaging Samples for Shipping to a Regional Laboratory

**Always review lab submission instructions, as packaging/shipping/submission requirements may differ between labs, and change depending on the current AIV situation and their capacity.**

a. Ensure that vials are tightly closed and securely packed to avoid sample leakage during transport.
b. Place vials in Ziploc bags (separate bag for each location).
    - On each Ziploc bag, write the number of vials contained in the bag, in addition to collection information (i.e., sampling location, date).
c. Wrap Ziploc bags containing vials in 5-6 layers of newspaper if using gel-type icepacks (not needed for dry ice).
d. Place Ziploc bags of vials on dry ice (preferred) or gel-type ice-packs inside a Styrofoam or plastic cooler
    - If using dry ice, use containers that meet standards for Transportation of Dangerous Goods, Type 1B packaging, i.e., watertight inner packaging surrounded by absorbent material, watertight secondary inner packaging and sturdy outer packaging (corrugated cardboard).
e. Securely tape cooler closed.

- If using dry ice, do not seal the lid completely with tape to allow the dry ice to evaporate through the cracks. The packaging must permit the release of carbon dioxide gas and prevent a build-up of pressure that could rupture the packaging. Please consult TDG training materials for additional information.

f. Place cooler into a sturdy cardboard box and surround with cushioning material to prevent damage to the vials or cooler. This cushioning material should be an absorbent material (i.e. newspaper), if gel-type ice packs are used to chill the vials.
g. Place the **completed sample submission form in a window bag/Ziploc bag** and into the cooler or sturdy cardboard box and ensure that the number of samples/containers matches the number indicated on the AIV field log.
h. Securely tape cardboard box closed.
i. To ensure compliance with subsection 13(1) of the Migratory Bird Regulations, any package or container for shipment or transport containing a migratory bird or its parts, nest, or egg must have its exterior clearly marked with:
    - the name and address of the shipper;
    - the number of any permit under which the birds, nests or eggs were taken, and;
    - an accurate statement of its contents.
j. Prior to shipping samples, please email completed field logs to your AIV surveillance coordination contact, as some regional labs require AIV sample IDs to be submitted via email prior to submission.

##### 2.4.3 Shipping Samples to a Regional Laboratory for PCR Screening

a. When you are ready to ship the samples, please email your AIV surveillance contact, who will provide you with:
    - Fed Ex Account information; and,
    - Laboratory address and account information (the pre-screening (initial lab analysis) will be done at provincial labs).
b. Please ship on Monday, Tuesday or Wednesday in order to avoid the shipment being in transit on the weekend.
    - If overnight shipping is not possible, shipping on dry ice or with a cryoshipper is recommended.
    - The appropriate TDG training is required to send, carry, or receive a cryoshipper filled with liquid nitrogen for sample collection.
    - There are also additional shipping requirements for cryoshippers filled with liquid nitrogen, please consult TDG materials and training.
    - If shipping on dry ice, it is recommended to ship by ground as there are additional requirements if shipping by air in Canada. An additional label will be required if you are shipping on dry ice. See comment in section 2.4.2 describing the need for venting.
c. After shipping, please email the waybill to your AIV surveillance coordination contact.

#### 3.0 Additional Sampling Considerations

- Field members are required to abide by regional PPE and Personal Protective Measures protocols provided by your region or agency to accommodate worker safety regarding Avian Influenza, biosecurity and COVID-19 mitigation procedures.
    - Please review and consider the following guidance on health and safety:
        - ECCC THA-SWP #50 Wild Bird Handling

        - Public Health Agency of Canada guidance: https://www.canada.ca/en/public-health/services/flu-influenza/fact-sheet-guidance-on-precautions-handling-wild-birds.html
    - Refer to the handout (“Fieldwork Operations During the COVID-19 Pandemic”) and link (“Wildlife and SARS-CoV-2: Handling Guidelines”) provided for general guidance.
    - Note: A qualitative assessment regarding the risk of transmission of SARS-COV2 from humans to birds during bird-handling activities permitted by Environment and Climate Change Canada was conducted in June 2020. The expert panel considered the risk of birds becoming infected with SARS-CoV-2 through bird handling to be extremely low.
- The birds will be under minor stress for a short duration. All team members handling birds and collecting swabs must be highly skilled with years of experience or have been appropriately trained and under supervision by experienced personnel.
- All birds are handled with the utmost care to ensure that any injury is avoided, and handling time and stress are minimized.
- Everyone in the field crew is responsible for monitoring the immediate health condition of the waterfowl before and after sampling.
- In the low likelihood event that the animal has been severely injured by predators or conspecifics while in traps, or if underlying disease conditions are precipitated by the stress of capture it would be up to the crew leader of the banding station to make the call whether that animal should be euthanized or brought to a wildlife rehabilitation facility, as these activities are conducted under their respective animal care permits. However, should animals be injured during our sampling procedures as stated in this animal care protocol, then the decision to euthanize, release, or bring to a wildlife rehabilitation facility will be made by the crew leader in collaboration with the CWS lead.
- Should animals show signs of severe pain or distress, abort the procedure, place the animal in a dark, quiet place (e.g., covered cage) for a short period of time, and will release the animal upon satisfactory evaluation of its well-being. This includes any animals that have been injured by trapping methods, predators, or conspecifics, or those showing signs of potential overheating (e.g., open-mouthed breathing).
- If you have any questions regarding status coding when submitting the data in Bandit 4.0 please contact the [redacted]

### Appendix H: List of Animal Health Diagnostic Laboratories

The Canadian Animal Health Surveillance Network (CAHSN) is a network of federal, provincial, and university animal health laboratories across Canada. Analysts from network laboratories across Canada have received training from the Canadian Food Inspection Agency National Centre for Foreign Animal Disease (CFIA-NCFAD) on how to use standardized testing protocols to detect four major foreign animal diseases including notifiable avian influenza. A list of CAHSN partner laboratories can be found at the following link:

Canadian Animal Health Surveillance Network - Canadian Food Inspection Agency (canada.ca)

### Appendix I: Additional ECCC Protocols, Policies, and Guidance Documents of Relevance for Surveillance Implementation

**Note***: that the development of AIV related protocols, policies, and guidance documents is still an active area of work. The following table lists relevant documents and their intent. These documents may be in various stages of approval. For more information on any one document please contact [redacted]. These documents are evergreen and will be updated as needed.*

[Table contents redacted]
